# AtSWEET13 transporter discriminates sugars by selective facial and positional substrate recognition

**DOI:** 10.1101/2022.10.12.511964

**Authors:** Austin T. Weigle, Diwakar Shukla

## Abstract

Transporters are targeted by endogenous metabolites and exogenous molecules to reach cellular destinations, but it is generally not understood how different substrate classes exploit the same transporter’s mechanism. Any disclosure of plasticity in transporter mechanism when treated with different substrates becomes critical for developing general selectivity principles in membrane transport catalysis. Using extensive molecular dynamics simulations with an enhanced sampling approach, we select the *Arabidopsis* sugar transporter AtSWEET13 as a model system to identify the basis for glucose versus sucrose molecular recognition and transport. We find that AtSWEET13 chemical selectivity originates from a conserved substrate facial selectivity demonstrated when committing alternate access, despite mono-/di-saccharides experiencing differing degrees of conformational and positional freedom throughout other stages of transport. In summary, our results point to a potentially generalizable finding that selectivity in transporters emerges from molecular recognition events occurring within regions distal from any conserved (non)functional binding sites.

## MAIN

Membrane transporters predominantly control the developmental and metabolic fates of cells through water, ion, and carbon allocation. Transport catalysis does not involve breaking or making of covalent bonds to form some product; instead, membrane transporters undergo large conformational changes to simply change a substrate’s physical position rather than its chemistry.^1,2^ Whether by design or on accident, transporters uptake and export molecules outside of their canonical or known substrate scope.^3^ Still, outside of just a few static crystal structures captured in different conformations, it is unknown exactly how a typical membrane transporter’s alternate access mechanism can be generalized when discriminating between different substrates.

A best-case scenario for approaching this research question would be to involve a relevant model system that transports similar yet structurally distinct substrates and does so without simultaneous cotransport of ions or additional molecules. Surprisingly, there exist few cofactor-independent transporters with readily available crystal structures.^4^ SWEETs (**S**ugars **W**ill **E**ventually be **E**xported **T**ransporters) are universally expressed, cofactor-independent, bidirectional uniporters responsible for sugar transport in plants.^5–7^ Evolved from bacterial SemiSWEETs possessing a single triple helix bundle, plant SWEETs contain two triple helix bundles connected by an inverted transmembrane helix linker (TM4).^8,9^ The evolutionary advent of TM4 introduced topological symmetry through inverted repeats, likely to structurally reinforce the functional need for bidirectional transport activity.^10^ SWEET substrate specificity generally aligns with sequence similarity: phylogenetic clades I, II and IV transport monosaccharides, while clade III prefers sucrose but has also demonstrated glucose and gibberellin transport activity.^11–15^ Their tissue-based expression and substrate specificities for mono- and di-saccharides underscore their significance in maintaining plant physiology through proper sugar signaling. This signaling can be disrupted during plant-pathogen interactions, where pathogens hijack SWEET functionality to redirect sugars away from the plant host. Specifically, clade III SWEET promoters are targeted during *Xanthomonas* infection to alter their expression and facilitate bacterial blight pathogenesis – an etiology currently known to devastate rice yields for African and Asian subsistence farmers.^11,16,17^

Given its substrate scope (glucose, sucrose, and gibberellin), cofactor independence, physiological/societal importance as a clade III SWEET, and available crystal structure (PDB ID: 5XPD),^18^ AtSWEET13 emerges as an excellent candidate for studying how different substrates activate the same membrane transporter. Beginning with the inward-facing (IF) AtSWEET13 crystal structure,^18^ we implemented an adaptive-sampling based regime with classical molecular dynamics (MD) simulations. Similar approaches have proven to be excellent strategies for studying membrane transport protein mechanisms.^19–23^ Herein, we resolve the conformational landscapes depicting AtSWEET13 *apo, holo* glucose (GLC), and *holo* sucrose (SUC) transport cycles. After statistically validating our ~450 μs of aggregate simulation with Markov State Models (MSMs),^24–28^ we identified regions along the AtSWEET13 transmembrane channel critical for differentiating GLC from SUC molecular recognition.

### Resolving the AtSWEET13 transport cycle

AtSWEET13 gating dynamics can be described by the extracellular and intracellular gating distances between K65-D189 and F43-F164, respectively.^18^ In line with prior work on SWEET gating dynamics,^19,20^ the inward-facing (IF) conformational state occupies the lowest free energy state (Figure 1A). This finding corroborates SWEET dependence on biological sugar concentration gradients,^5,11^ as this bidirectional uniporter family can be expected to remain in the IF state until enough sugar has accumulated to require export to sink destinations. Additionally, the MSM-weighted *apo* AtSWEET13 transport cycle includes an hourglass-like (HG) intermediate state which experiences a higher energy barrier for transitioning directly to the outward-facing (OF) state, rather than through the occluded (OC) state (Figure 1B). HG states have been previously observed during the transport cycle of rice glucose transporter OsSWEET2b, where the HG conformation preserves the crux of alternate access by constricting the transmembrane channel pore radius.^19^ Meanwhile, AtSWEET13 conformational change experiences higher energy barriers for transition during *holo* GLC and SUC transport cycles, where access to intermediate states in GLC transport are ~0.9-1.6 ± 0.7 kcal/mol higher, and ~0.2-0.3 ± 0.3 kcal/mol higher in SUC transport, when compared to *apo*. Like the *apo* simulations, both MSM-weighted *holo* datasets indicate the IF conformation as the lowest energy state; however, *holo* simulations uniquely stabilize transitions for an alternate access mechanism passing through an HG conformation while destabilizing an OC-based alternate access pathway. An HG gating distance of ~8-10 Å, rather than an OC aperture of ~5 Å, is likely needed to accommodate the molecular recognition processes for each substrate (Figure 1C). It is important to note that the existence of HG states in *holo* sugar transport does not violate the principle of alternate access, as this common transporter topology maintains a blocked channel to prevent substrate leakage.^19,29,30^ Overall, these findings are consistent with the “free ride” mechanism, where the *apo* and *holo* transport cycles access the same protein conformational changes, albeit at different relative free energies and transition rates.^31^ The relationship between the higher energy barriers for GLC-induced gating versus SUC-induced gating also corroborates experimental evidence that AtSWEET13 has evolved to be a more efficient disaccharide transporter.^18^

**Figure 1.**
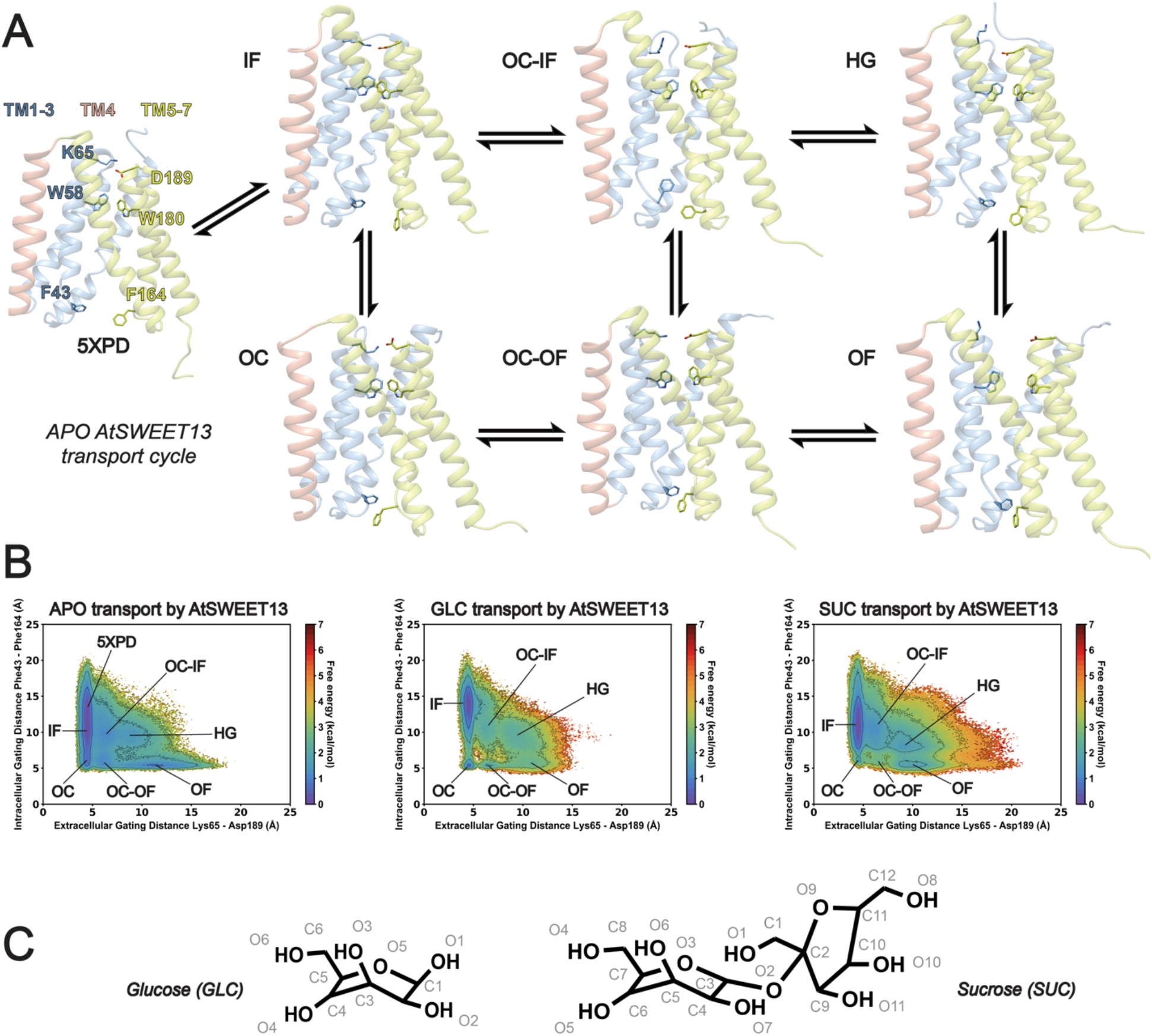
AtSWEET13 transport cycle. (A) *Apo* conformational snapshots for each representative state. (B) Gating landscapes for *apo*, GLC, and SUC transport processes, with key intermediate states labeled based on extracellular and intracellular gating distances. (C) Glucose (GLC) and sucrose (SUC) substrate molecular structures with atoms numbered as parameterized for molecular dynamics simulations.

### Conformational change cannot explain differences in AtSWEET13 sugar transport

To determine exactly how AtSWEET13 differentiates GLC versus SUC transport, free energy landscapes specific to individual substrate atoms were projected versus substrate position and AtSWEET13 gating distances (see Online Methods, Supplementary Figures 3-8). Figure 2 depicts gating difference versus substrate atom position landscapes, where the atom under inspection is the hexose backbone ether oxygen (Figure 2A, B). Upon binding from the intracellular face of the transporter, both sugars must traverse 25 Å up into the transmembrane channel to reach the hydrophobic binding pocket containing W58 and W180. Due to the topological asymmetry of AtSWEET13, where the hydrophobic binding pocket containing W58 and W180 is arranged more closely to the OF-side of the transmembrane channel, much of GLC and SUC translocation occurs while AtSWEET13 is in an IF-favoring conformation (Figure 2A, B). Indeed, as the sugars approach the W58-W180 binding pocket, the intracellular gating distance is ~10 Å greater than the extracellular gating distance for GLC and ~7 Å greater for SUC. Binding events along the IF association pathway for SUC (SUC_1_ → SUC_2_ → SUC_3_) are at least ~1.0 ± 0.1 kcal/mol more stable than as seen for GLC (GLC_1_ → GLC_2_; Figure 2A, B). Both GLC and SUC then approach a stable intermediate state after climbing 25 Å up the transmembrane channel (GLC_3_; SUC_5_), during which point AtSWEET13 commits each substrate to alternate access for the remaining 15 Å of membrane-spanning distance. Projection of raw gating distance data onto these gating difference distance plots reaffirms how alternate access occurs through an HG pathway (Supplementary Figures 1-2). Commitment to alternate access is reflected based on the conformational changes between the GLC_3_ → GLC_4_ states and the SUC_5_ → SUC_6_ states, where SUC_6_ is ~1.2 ± 0.3 kcal/mol more stable than GLC_4_ (Figure 2A, B). These transitions invoke the greatest protein conformational change, where the per-residue RMSD of nonterminal amino acids is surprisingly greater for the GLC HG-OF transition than the same transition for transport of the larger SUC molecule (Supplementary Figure 9). Across all transitions between numbered metastable states, the average per-residue RMSD is also greater for overall GLC transport than for SUC (Figure 2C, D). Noteworthy of these highly fluctuating regions is that the residues involved in the largest extent of conformational change do not reside within the central binding pocket. These residues instead predominantly line the AtSWEET13 periphery, echoing the importance of distant residues for evolved function in other proteins, including membrane transporters.^32–35^ Still, AtSWEET13 maintains high fluctuations within these regions for both GLC and SUC transport reactions, suggesting that movements from these residues are essential for general alternate access regardless of substrate.

**Figure 2.**
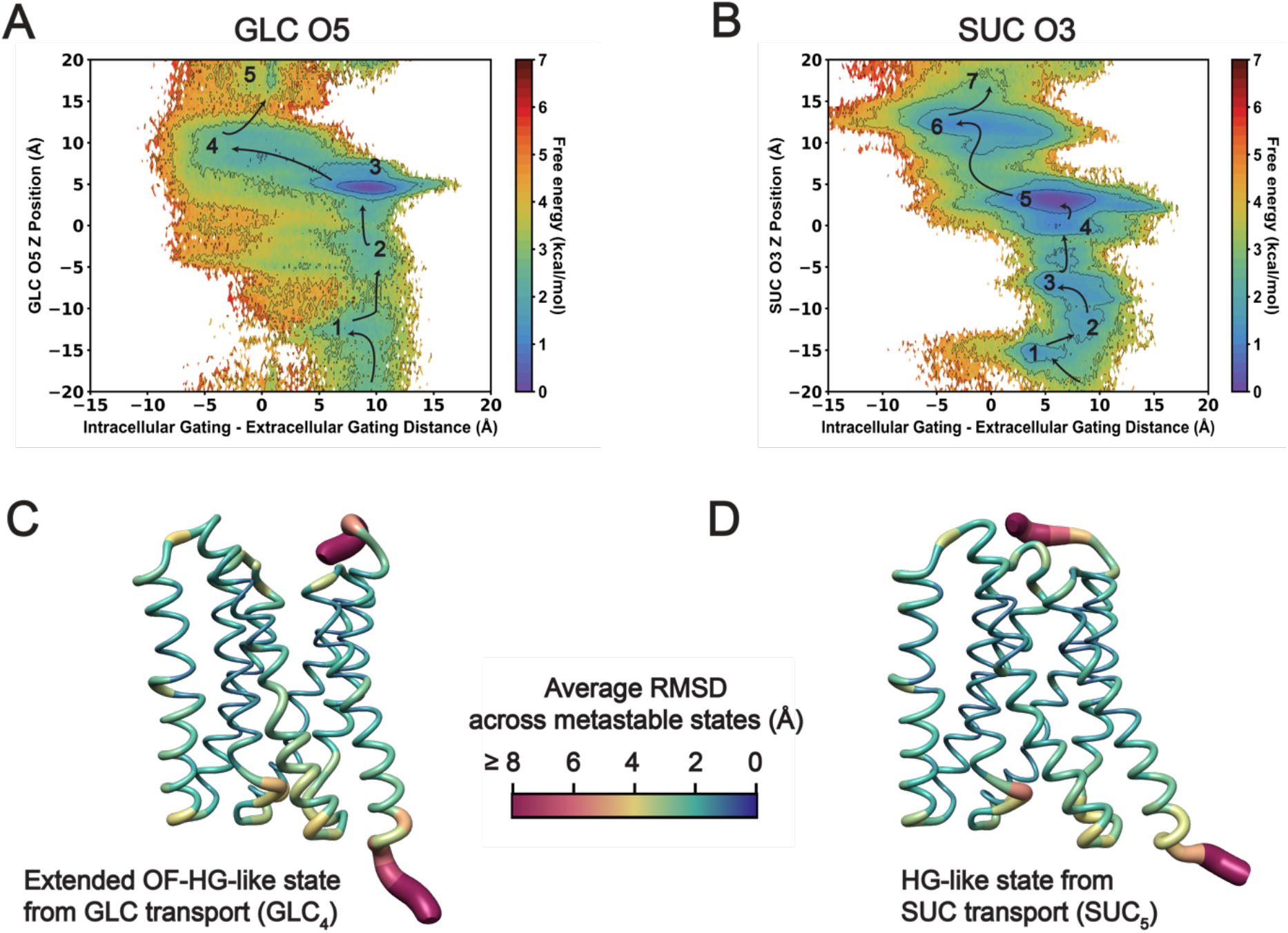
Protein conformational change implicated in sugar transport. (A) GLC transport versus the difference in intracellular versus extracellular gating. AtSWEET13 is in a more IF-like position the more positive the gating difference and a more OF-like position the more negative the gating difference. (B) SUC transport versus the difference in intracellular versus extracellular gating. (C) Protein snapshot of AtSWEET13 upon commitment to alternate access during GLC transport. An average RMSD value is calculated across all numbered metastable states and projected as a worms representation onto the HG-like structure for GLC transport. (D) Protein snapshot of AtSWEET13 in an HG-like conformation during SUC transport, with averaged RMSD values projected as a worms representation, as seen in (C). Transport cycles involving each of the above numbered states are shown in Supplementary Figure 9. Arrows drawn over landscapes are for illustrative purposes to suggest the transport path according to the lowest MSM-weighted free energy transitions as shown.

When projecting the same data using each heavy atom in SUC as the reference substrate atom, the SUC_5_ → SUC_6_ transition is smoothened for fructosyl atoms (Supplementary Figures 5-8). AtSWEET13 preference of SUC transport thus appears to be reflected by the evolved incorporation of the pentose moiety into the disaccharide structure, where its presence helps facilitate commitment to alternate access better than the SUC hexose unit. Additionally, SUC atom positioning throughout the early IF uptake stages of transport does appear to be more stable than seen for GLC, suggesting that transport binding stages apart from the OC and HG states are controlled by different processes for translocation of the two sugars.

Time-lagged independent component analysis (tICA) used to determine appropriate features for MSM construction and reweighting further corroborates how GLC transport is more reliant on protein conformational change (Supplementary Figures 1,11; Supplementary Table 1).^36,37^ Second order tICA decomposition has demonstrated that the first and second time-lagged independent components (tICs) relate to the slowest, or rate-determining, processes for a given biomolecular system.^36^ During initial iterations of MD data featurization, exhaustive attempts were made to ensure that the amount of adaptive sampling carried out sufficiently captured the dynamics of transport. To this end, the aggregate datasets for *apo*, GLC, and SUC transport were described using between 100-1000 residue-residue distances for transmembrane channel residue pairings, along with substrate atom-residue distances to the select channel residues (see Online Methods).

Representations of these more feature-diverse tICA discretization attempts for AtSWEET13 transport were used to shortlist a set of features that would best describe transport dynamics contained within the aggregate dataset in a final MSM (Supplementary Figures 12-17; Supplementary Tables 3-5). A “best” description of transport involves features which properly discretize metastable transport states from one another. Additionally, resulting discretization should not cause free energy landscape reweighting that significantly or artificially deviates from trends seen during raw data collection based on poor clustering. Iterative attempts at MSM construction were made until feature sets satisfied these criteria (see Online Methods).

After feature set reduction, the slowest tICs for monosaccharide transport were highly correlated with 13 protein residue-residue distances whereas just six residue-residue distances were highly correlated with disaccharide transport (Supplementary Figures 1,10). Additionally, tICA landscape projection depicts *apo* and SUC transport as dominated by a singular process while GLC transport is highly correlated along two distinct processes (Supplementary Figure 1). For *apo* transport, tIC1^apo^ signified protein conformational change, as the most correlated features related to protein residue-residue distances. For SUC transport, tIC1^SUC^ reflected glucosyl moiety atom positioning while tIC2^SUC^ reflected fructosyl moiety atom positioning, together suggesting overall substrate translocation is rate-limiting. Lastly, for GLC transport, tIC1^GLC^ reflected protein conformational change while tIC2^GLC^ reflected monosaccharide substrate atom positioning, suggesting greater extents of protein conformational change are required for GLC transport (see Online Methods sections concerning MSM construction and feature selection; Supplementary Figures 13-17).

Despite the increased dependence on protein conformational change for monosaccharide transport, transmembrane channel pore radius between GLC and SUC transport share similar size as well as morphological changes during the transport cycle (Supplementary Figure 9). AtSWEET13 exhibits a similar extent of IF gating aperture when presented with monosaccharide GLC versus disaccharide SUC, despite SUC being a larger molecule. During transport of either substrate, major changes in transmembrane channel pore radius only occur after commitment to alternate access (Supplementary Figure 10). From the perspective of protein conformational change, these findings signify that AtSWEET13 surveys a relatively similar distribution of protein dynamics to commit each native substrate to alternate access. Since the general gating mechanics of AtSWEET13 are predicted to remain relatively the same between the two substrate transport processes, it is more likely that preferential substrate-specific interactions occurring within the transmembrane channel may act as determinants of selectivity.

### Stages of discriminative sugar recognition occur across distinct segments of transmembrane-spanning channel residues

The extent of similarity in AtSWEET13 gating is somewhat expected, as a transport pathway can only be so plastic to maintain transport of different classes of molecules. However, the differences in how energetically favorable intermediate states were along IF association pathways for GLC O5 and SUC O3 are intriguing considering they represent the same ether oxygen along nearly identical hexose units. Atoms along the SUC pentose atom also experienced lower energy barriers during commitment to alternate access (Supplementary Figures S7,8). If the incorporation of a glycosidic linkage is enough to make the same atom between GLC and SUC be no longer perceived as functionally equivalent by AtSWEET13, and if the two monomeric units of SUC experience different energy barriers, then atom-specific molecular recognition may be satisfactory in explaining how the two sugar types are differentiated.

Treating AtSWEET13 as a “physical enzyme”^1,2^ requires taking facial and spatial selectivity into consideration. Accordingly, each GLC and SUC atom was analyzed via a vector representation with respect to its Z position during transport (see Online Methods for AtSWEET13 atom-specific analyses). Briefly, the angle between the atom of interest and the center of mass of the protein along the XY plane (***θ***_xy_) indicates ligand spinning, or spatial selectivity. Conversely, the angle between the atom of interest and the ligand’s center of mass along the XZ or YZ planes (***θ***_xz_ or ***θ***_yz_) indicates ligand flipping, or facial selectivity (Figure 3C). These atom-specific rotational analyses were performed on each GLC and SUC atom (Supplementary Figures S18-35). Based upon these analyses, AtSWEET13 appears to recognize the GLC C3/O3 and SUC C5/O6, as well as the GLC C4/O4 and SUC C6/O5, hydroxyl groupings as roughly equivalent (Supplementary Figure 36). Figure 3B presents the ***θ***_xy_ spinning rotational analyses for GLC C3 and SUC C5 for distinguishing GLC from SUC IF, as well as the ***θ***_yz_ flipping rotational analysis for GLC O5 and SUC O3. Most notably, discriminative binding events occur along specific segments, or “layers”, within the AtSWEET13 transmembrane-spanning channel (Figure 3A).

**Figure 3.**
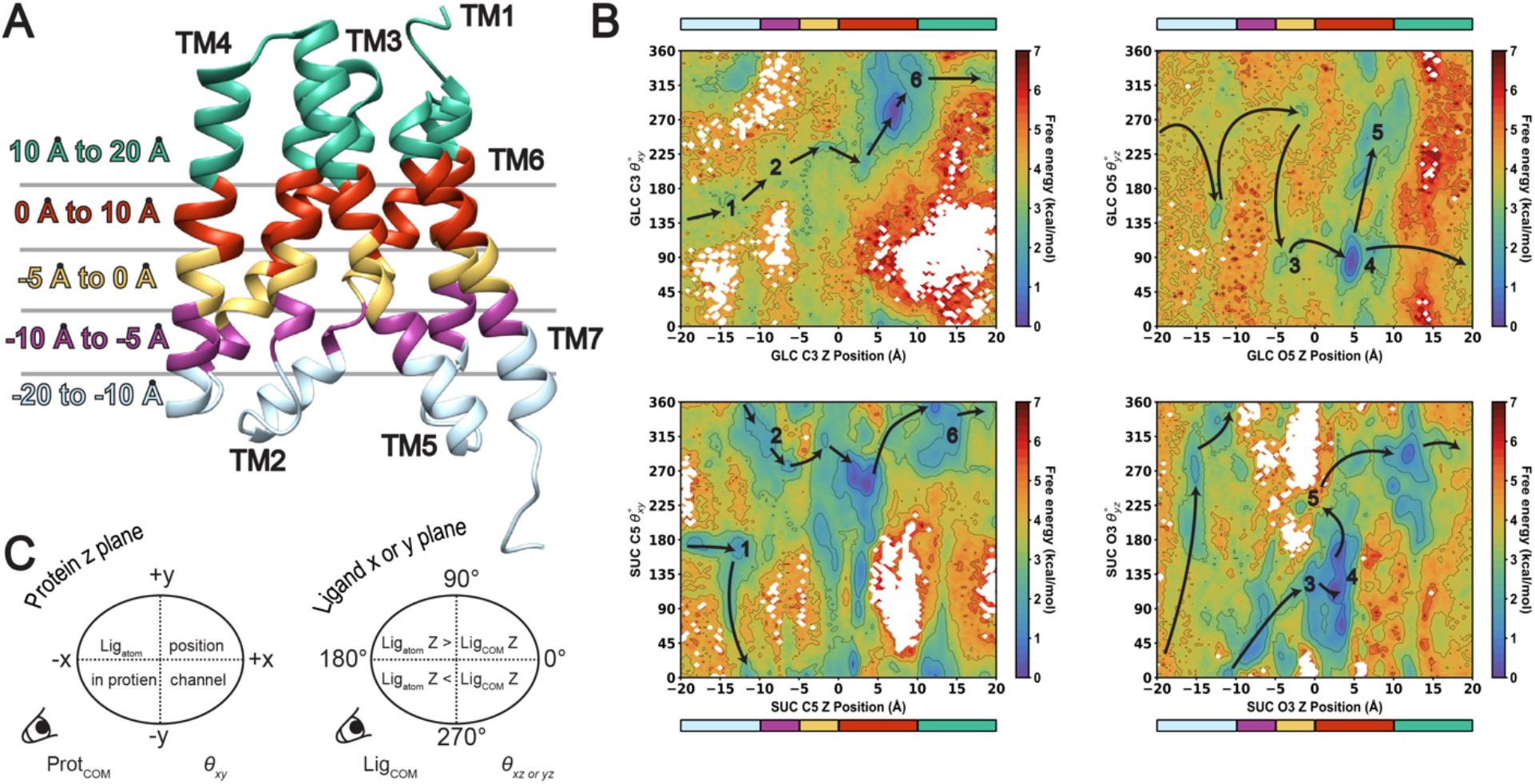
AtSWEET13 layers for discriminating sugar substrates. (A) AtSWEET13 5XPD crystal structure with layered coloring based on residue roles in stage of sugar transport. (B) Atom-specific **θ** rotational plots, measuring ligand atom Z position versus rotational orientation. **θ**_xy_ is shown between nearly equivalent atoms GLC C3 and SUC C5 while **θ**_yz_ is shown for GLC O5 and SUC O3. Colorbars above and below each plot allow direct comparison to the colored 5XPD structure in (A). (C) Schematic explanation for interpreting **θ**_xy_ and **θ**_xz_ or yz calculation. **θ**_xy_ captures the angle between the protein center of mass (COM) and the specific ligand atom coordinates in the XY planes. **θ**_xz_ or yz captures the angle between the ligand (COM) and the specific ligand atom coordinates in the XZ or YZ planes. Arrows drawn over landscapes are for illustrative purposes to suggest the transport path according to the lowest MSM-weighted free energy transitions as shown.

Upon first binding to the IF AtSWEET13 state, GLC C3 and SUC C5 enter along the same face of the protein with 90° ≤ ***θ***_xy_ ≤ 200°, or in between TM4 and TM5 (Figure 3A, cyan layer; Figure 4, GLC_1_/SUC_1_). GLC IF association then must proceed within the range of 135° ≤ ***θ***_xy_ ≤ 225°, as the GLC C3 atom samples transmembrane channel space between TMs 5, 6 and 7 (Figure 3A, purple later; Figure 4, GLC_2_). Although averaging a free energy of ~3.4 ± 0.3 kcal/mol, note that this is the only predicted IF association pathway available for GLC as it travels up the first 15Å up the AtSWEET13 channel, inspiring us to refer to this rotational space as part of the “monosaccharide pathway” (Figure 3B, top left panel, cyan and purple layers). SUC C5 is also able to access the monosaccharide pathway during early IF association with similar free energy penalties as GLC C3, but can also access other regions of the channel at nearly half the free energy cost. SUC C5 instead is more likely to occupy regions where ***θ***_xy_ > 225° (Figure 3B, bottom left panel, cyan and purple layers). In contrast to GLC, the SUC_2_ state therefore leads the GLC-like atom towards the TM6-TM1 cleft, space which the equivalent atom in the GLC structure does not approach during the same stage of transport. Thus, we term this area of AtSWEET13 as part of the “disaccharide pathway”.

**Figure 4.**
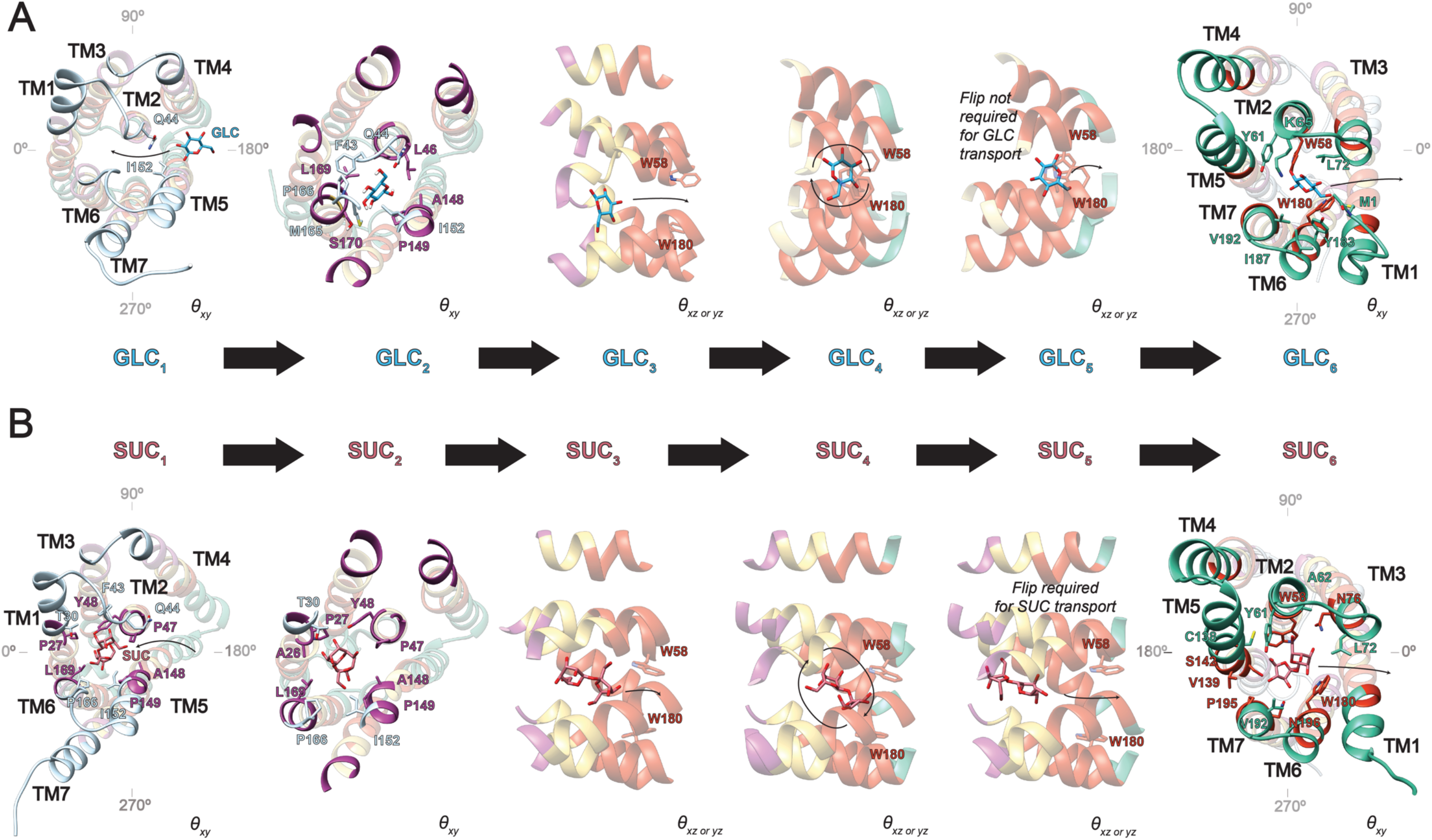
Discriminative sugar transport events for AtSWEET13. States are shown for (A) GLC transport and (B) SUC transport. Each of the numbered states correspond to those enumerated in Figure 3B. Each snapshot is accompanied by a marker for indicating which type of rotational analysis the structure is intended to convey.

Regardless of mono-versus di-saccharide pathway for early IF association, the progression from an OC-IF to a more HG-like state results in experiencing similar rotational freedom. But like the gating difference plots from Figure 2, SUC atoms experience lower energetic penalties as they sample ***θ***_xy_ space and contact each AtSWEET13 TM. One reason for this discrepancy in energetic penalty in transit to the W58-W180 binding pocket is the varying degrees of facial freedom experienced by each substrate (***θ***_yz_; Figure 3B, right panels). During IF association, GLC can invert freely while occupying the TM5-TM6-TM7 cleft between the cyan and yellow AtSWEET13 layers (Figure 3A). Interestingly, SUC atom IF association instead can only flip and cross the ***θ***_yz_ = 180° boundary within just the cyan layer. SUC facial orientation is then relatively fixed when traveling throughout states SUC_2_ → SUC_3_ → SUC_4_ (Figure 3B, bottom right). On the other hand, GLC samples different facial orientations prior to binding to the W58-W180 pocket, where this conformational selection strategy is enabled by the smaller ligand’s size. As SUC expresses less facial freedom, transport can rather focus on rotational substrate positioning such that a productive “pre-transition state” can be available for commitment to alternate access. Increased facial freedom for GLC instead results in variable orientations which likely require correction from more involved protein conformational change. AtSWEET13 is essentially playing a game of Rubik’s cube with the two substrates, where increased entropy in possible GLC facial orientations requires more correction by transmembrane-spanning channel residue interactions to enable a competent substrate orientation for transport catalysis.

Nevertheless, both predicted substrate transport pathways involve committing GLC and SUC to particular facial orientations to allow alternate access (Figure 4; GLC_4_ → GLC_5_ and SUC_4_ → SUC_5_ transitions). SUC transport requires substrate inversion where the hexose unit is ***θ***_yz_ < 180° and leads the disaccharide structure from the IF face of the transporter. Upon reaching the binding pocket (red layer), the pentose unit leads the disaccharide substrate towards OF dissociation as the hexose unit assumes ***θ***_yz_ > 180° (Figure 4B). Monosaccharide GLC can flip throughout what could be perceived as the transition state of translocation catalysis even while bound directly to W58 and W180 (GLC_4_ → GLC_5_; Figures 3B, 4A). Considering that the equivalent ether oxygens between GLC and SUC both have ***θ***_yz_ ≈ 225°, it is possible that GLC flips freely, perhaps signifying attempts to satisfy recognition requirements for either monomeric unit of SUC. Such an explanation is likely the primary reason for why GLC samples the unproductive state GLC_5_, as GLC_3_ already expresses the correct facial orientation to commit directly from GLC_3_ → GLC_4_ → GLC_6_ and final OF dissociation. Substrate departure for GLC and SUC OF dissociation occurs through TMs 1 and 3 (Figure 4). Rotational analyses for each substrate heavy atom are provided in the Supplementary Information (Supplementary Figures S18-35).

Overall, these atom-specific rotational analyses reveal the functional differences between the hexose and pentose units within SUC. Visual inspection of facial and spatial orientational preferences for substrate positioning by AtSWEET13 allow for the rationalization of transportable-sugar structure-activity relationships (Supplementary Figure 36). Specifically, IF molecular recognition of the SUC hexose unit emphasizes that it can be perceived like monomeric GLC by potentially sampling the monosaccharide path, but that this mode of transport is likely to be less probable based upon MSM-weighted relative free energy calculations. AtSWEET13 specialization in disaccharide transport therefore emphasizes the importance of pentose molecular recognition during HG-OF transitions and OF dissociation.

### Exploiting evolutionary context for mechanistic generalization and predictive engineering

Viewed under evolutionary contexts, our proposed mechanism suggests AtSWEET13 can tap into some “primordial memory” to first position substrates based on IF molecular recognition of glucose-like atoms. The increased plasticity in SWEET transport, as well as the ability for AtSWEET13 to invert its substrates in the HG state, is likely a reflection of the inverted symmetry gained through the evolutionary advent of TM4 as SWEETs evolved from SemiSWEETs.^8,9^ With reference to the topological asymmetry of AtSWEET13, it makes sense that a difference between mono- versus di-saccharide transport is the variation in IF association pathways, as a majority of metastable transport events occur while AtSWEET13 is in an IF-like conformation (Figure 3B). Transport catalysis is thus expected to ensue with high efficiency once an intermediate conformational state is achieved with the substrate in an optimal facial orientation. Given the barriers between intermediate and OF states are relatively flat, it can be proposed that the AtSWEET13 substrate transport mechanism mimics the catalysis-promoting strategy of effective intermediate barrier reduction as seen in enzymes utilizing ground state stabilization for rate enhancement.^38^

To make AtSWEET13 more selective for either GLC or SUC would then imply the need for mutations which would lower the effective barrier between the IF ground state and the flat intermediary free energy landscape, or selectively reduce the barriers between intermediate states, in preference of one substrate type. More specifically, selective mutations would eliminate the possibility for monosaccharide IF association pathway sampling or the need for substrate inversion during commitment to alternate access.

In the former case, enhanced SUC selectivity would be predicted as GLC would be unable to associate and bind to AtSWEET13 from the intracellular opening. In the latter case, substitutions nearby the conserved W58-W180 binding pocket could prevent SUC facial inversion necessary for catalysis, only permitting transport of GLC molecules that had preselected the desired facial orientation for alternate access. Such changes to amino acid composition may possibly be reflected in the evolutionary contexts for specialized transport across the different SWEET clades.^11^

## DISCUSSION

Just like chemical enzymes, transporters demonstrate facial selectivity. Our study suggests that chemical selectivity in the AtSWEET13 sugar transporter originates from the protein’s ability to optimally position a ligand for recognition before committing to intermediate transitions and alternate access. So long as a ligand’s functional group(s) can resemble those of a more native or ancestral substrate, and the protein can recognize it similarly, the ligand can bind, initiate conformational transitions, and be transported. Differences in size do influence the ability for different substrates to sample different facial orientations throughout the transport cycle, where facial selectivity appears to be critical for transport catalysis.

Increases in free energy barriers for *holo* AtSWEET13 sugar transport are surprising, but placing the transport process into biological context offers clarity. SWEET transporters are bidirectional, cofactor-independent uniporters that catalyze sugar translocation in agreement with a concentration gradient. Catalytic efficiency from reduced effective barriers between intermediate states in response to concentration gradients would also be coupled to an endergonic, downhill, and immediate, substrate uptake into cells when required. Critical sink destinations like meristematic cells for embryo development; epidermal cells during stem elongation for cell wall deposition and sugar distribution; stigma and ovary cells for aiding nectar secretion; and trichome cells for the development of defensive sugar-based metabolites would benefit from a seemingly on-demand regulation of sugar allocation.^39^

AtSWEET13 alternate access accommodates substrate transport by assuming an HG state where the gating aperture is large enough to house sugar molecules and permit necessary atomistic rotations. Matching its experimentally characterized function, AtSWEET13 transport of GLC is predicted to be a higher energy process than SUC transport. Per the tICA decomposition analysis, monosaccharide transport is highly dependent on both protein conformational change and substrate positioning, whereas disaccharide transport mirrors *apo* transport efficiency in that its transport cycle is highly correlated along a single dimension rather than two (Supplementary Figure 1) While the extent of protein conformational change between GLC and SUC transport follows similar trends, mono-versus di-saccharide transport pathways bifurcate along early molecular recognition events during IF association. These differences in functional behavior across the different stages of AtSWEET13 transport were characterized thanks to the label-free atomistic resolution of MD simulations validated by feature-agnostic MSMs.

Our study offers insights into (1) how cofactor-independent transporters are predicted to accommodate different substrates within the same general transport scheme and (2) how general transport selectivity can be engineered. In the case of predicted sites for engineering selective monosaccharide transport, these residues surround the conserved W58-W180 pairing constituting the binding site. Residues within these regions are responsible for uniquely preparing different substrates so that they may most efficiently approach a transition state conformation suitable for transport. The extent to which these residues predetermine a substrate orientation capable of committing to catalysis can be expected to be a function of their distance from heavily conserved sites and their own degree of conservation within the families of related proteins.^35^ The local and allosteric roles of these identified residue sites in maintaining fundamental differences in selective substrate transport are expected to be targetable provided evolutionary contexts.

Adding to the difficulty of study based off their non-soluble nature,^40^ efforts to reveal membrane transporter selectivity principles are complicated by the absence of local catalytic hallmarks as seen in chemical enzymes. Outside of the canonical approach of identifying transporter global conformational changes, we introduce the importance of substrate orientation during the promotion of translocation catalysis by these “physical enzymes” using MD simulation. Generally, we expect the importance of targeting residues peripheral to conserved sites to be an important strategy in efforts for the design of dedicated transporters or highly selective small molecules. However, the targeting of conserved residue sites is likely permissible provided the allosteric relationship with distal site residues is understood. Ultimately, the goal of engineering transport selectivity is to alter the ability by which critical facial and spatial intermediate substrate orientations become accessible throughout the transport reaction. In the case of SWEET sugar transport, we hope our efforts help contribute towards the development of blight-resistant crops and improved international food security.

## ONLINE METHODS

### Membrane and membrane protein system assembly

For AtSWEET13 simulations, an inward-facing (IF) *Arabidopsis thaliana* AtSWEET13 crystal structure (PDB: 5XPD) was acquired from the RCSB Protein Data Bank (PDB).^18,41^ Using CHARMM-GUI, AtSWEET13 was modeled off residues 1-222, where the thermostable mutations performed to aid in crystallization were reverted.^18,42–44^ The protein was inserted into a realistic, asymmetric model plant plasma membrane following a Maximum Complexity composition.^45^ Addition of all water and ligand molecules for each system was performed using PACKMOL 18.169.^46^ All *holo* AtSWEET13 systems were built using 100 mM of substrate to match published experimental conditions and to shorten simulation time necessary for initial binding.^18^

### System parameterization

All ligands were prepared as PDB files in PyMol, where charge states were determined from pKa calculations using the ChemAxon Chemicalize tool.^47,48^ All ligands were then parameterized by applying the CGenFF36 forcefield using the CHARMM-GUI Ligand Reader & Modeler.^49,50^ CHARMM36 protein and lipid forcefields applied with the psfgen VMD-plugin,^51^ while the PDB2PQR (PROPKA) server was used to determine protein protonation states.^52^ The CHARMM TIPS3P water model was employed, and potassium and chloride ions were used to neutralize the systems using the VMD autoionize package. All systems prepared through VMD were then converted to AMBER format using the ParmEd CHAMBER package.^53^ Hydrogen mass repartitioning was applied to improve statistics by extending single trajectory lengths given available computing wall-clock times.^54,55^

### Simulation details

System preparation was performed as previously described for membrane protein simulations using the AMBER18 engine.^45,56^ A major deviation from the referenced membrane protein simulation protocol is that during production runs, a Langevin thermostat and Monte Carlo barostat were implemented for temperature and pressure maintenance, where the Langevin collision frequency was 2 ps^-1^.^57,58^ AtSWEET13 production runs ran for 80 ns. The SHAKE algorithm was applied to all stages of simulation initialization except for minimization,^59^ while the Particle Mesh Ewald method used for treating long-range electrostatics at a 12 Å cutoff.^60^

### Adaptive sampling protocols

The hallmark of parallelizable simulation seen in adaptive sampling is the ability to select any given state as a starting point for seeding a new trajectory.^61–66^ Starting states are selected through a kmeans clustering of available data given a reaction coordinate of interest, where states occupying the least populated clusters (i.e. least counts clustering) are used for seeding the next round of sampling. The “aggressiveness” of adaptive sampling regimes can be tuned depending on whether clustering and seed selection occurs using the data only collected from a given round or across all recorded rounds, as well as the number of seeds generated from each round. For *apo* simulations, distances between gating residues were used as the reaction coordinates for clustering. Monosaccharide simulations were also performed entirely using distances between gating residues. Adaptive sampling metrics for sucrose (SUC) transport changed depending on the stage of transport progress seen in simulation. At first, distances between gating residues were solely used to promote ligand binding. Once a SUC molecule was bound and no transport progress was noticeable along the free energy landscape projections after ~3 successive rounds of adaptive sampling, the sampling metric was changed. Atom Z position of the closest SUC molecule to binding site residues Trp58 and Trp180 was then used as the adaptive sampling metric until AtSWEET13 entered an OC/HG-like state. Once SUC was bound to Trp58 and Trp180, the sampling metric was again changed to account for both atom Z position and gating distances. AtSWEET13 extracellular and intracellular gating distances were calculated using residue pairings Lys65-Asp189 and Phe43-Phe164, respectively. From clustering of the provided metrics within the present round, a minimum of 200 states were seeded per round of adaptive sampling for each AtSWEET13 system, where restart coordinate files were prepared using CPPTRAJ.^67^ Seeded trajectories were simulated for at least 80 ns in length, depending on available computational resources. Starting from the AtSWEET13 crystal structure IF state, adaptive sampling proceeded until OF states were observed, substrate dissociation occurred, and the raw free energy landscapes were well connected (i.e. observable variation in the raw free energy landscape ceases). The use of MDTraj 1.9.3 for measuring distances, VMD 1.9.3. for visualizing states, and Matplotlib 3.2.0 for visualizing projected landscapes were the same as described previously.^51,68,69^

### AtSWEET13 atom-specific analyses

Because *holo* sugar simulations were run using multiple copies of the same substrate, the atom under inspection belonging to the ligand molecule found within the AtSWEET13 transmembrane channel was determined. The Z position of the specified atom for the substrate copy undergoing transport was then compared to either the difference between the extracellular and intracellular gating distances, or to a rotation angle, theta (***θ***). For the ***θ*** calculations, imagine a unit circle with a line drawn between points ***A*** and ***O***, where ***A*** represents an atom on the ligand and ***O*** represents the graphical origin as the protein (***O_prot_***) or ligand’s (***O_lig_***) center of mass. The angle between ***A*** and ***O_prot_*** in the XY plane (***θ***_xy_) represents the specified atom’s rotational freedom within the transmembrane channel of AtSWEET13, as the resulting vector signifies directional orientation (and thus contact) between the ligand atom and different regions within the transmembrane channel. Meanwhile, the angle drawn between ***A*** and ***O_lig_*** in the XZ or YZ planes (***θ***_xz_ or ***θ***_yz_) indicates the facial orientation of the substrate during different stages of transport (Figure 3C). Each of these analyses was conducted for every heavy atom (carbon or oxygen) found in each substrate scaffold.

### Determination of agnostic MSM descriptors

Although adaptive sampling regimes typically employ classical MD, the selection of seed states per adaptive round to improve the observation odds of conformational transitions is an inherently biased process. Thus, a fundamental objective for implementing the MSM technique is to statistically validate the aggregate ensemble of MD data through the removal of bias introduced during adaptive sampling seed state selection. A major challenge in the building of MSMs is deciding what features or molecular descriptors will be used for initial data clustering. While there is no exact standard as to how this featurization should be performed, it is generally understood that the MD practitioner should attempt to characterize the slowest dynamic modes while minimizing the use of molecular descriptors directly introduced during adaptive sampling.^70–73^ Accordingly, the selection of molecular descriptors for featurization should accurately capture slow dynamical processes relevant to the study objectives while having not been used directly during the adaptive sampling protocol. To this end, representative structures were taken from each relative metastable state along the raw free energy landscapes from select projections. Visual comparison between the monosaccharide and SUC free energy landscapes led to the identification of atoms which seemed critical for molecular recognition between the mono- and di-saccharide transport processes. For GLC, this meant collecting states from all C3, C4, O3, and O4 atom-specific landscapes. Critical atoms for SUC state extraction were identified as C2, C5, C6, C9, C10, C11, C12, O5, O6, and O9 landscapes. Metastable energetic minima states were extracted from these landscapes for each substrate dataset, as well as the general extracellular versus intracellular gating landscapes. From all these selected states, C-beta distances between all possible pairwise transmembrane channel residue-residue combinations were calculated (C-alpha was used for glycines). C-beta atoms were selected because of their ability to approximate both backbone and sidechain movement.^74^ Using the scikit-learn 0.21.2 library, the number of states used to determine the putative MSM features was filtered by identifying which states occupied kmedoids cluster centers. The number of clusters was equivalent to the maximum number of metastable states seen in any individual atom-specific plot for the specified simulation dataset. The inverse of all pairwise channel residue-residue distances were then calculated from this filtered list of metastable states for increased sensitivity,^75^ and the distances whose standard z-score was greater than three standard deviations from the distance distribution mean were selected as initial molecular descriptors for generating a feature-agnostic MSM. With an increased standard z-score, it can be assumed that an increased dynamical range (extent of change in the measured distance) has been accurately captured across the extracted metastable states.

### Markov state model (MSM) construction

PyEMMA software was used for MSM construction.^76^ With exhaustive, agnostic features determined, MSMs were generated using an iterative grid search procedure where the VAMP score was optimized.^77^ The MSM protocol used herein can be described as follows:

1. Given the exhaustive set of agnostic features, construct pilot MSMs to determine the range of cluster counts and time-Independent Components (tICs) capable of returning converged implied timescale plots.^78^ From these pilot runs, obtain a working MSM lag time to use for a more exhaustive grid search based on visual inspection of resulting implied timescale plots.
2. Perform an exhaustive grid search where the accepted hyperparameter combination optimizes the VAMP score. With this initial MSM transition probability matrix determined, project tIC1 versus tIC2 from the tIC analysis (tICA) object output during clustering.
3. Confirm that the tICA decomposition landscape from Step 2 is well connected. If so, then determine which features are most correlated with tIC1 and tIC2 for the currently optimized MSM. If not, revisit adaptive sampling to connect this complex tICA landscape with more data.
4. Once the tICA landscape has been connected, plot the raw cluster counts versus the MSM-weighted cluster counts for the tICA-converged dataset. If the extent of reweighting between the raw and MSM-adjusted data deviates beyond a relationship of *x*=*y* where *y* is much greater than *x*, then the molecular descriptors used for MSM featurization are deriving an overtuned bias correction.^79,80^ In this case, molecular descriptors that are highly correlated to tIC1 and tIC2 of the exhaustive tICA decomposition can be used to simplify the MSM featurization step.
5. Repeat Steps 1-3 using molecular descriptor feature sets inspired from Step 4 until the raw cluster counts versus MSM-reweighting plots approximately satisfy a relationship of x=y.

The final feature sets for each system are listed in Supplementary Table 1. MSM hyperparameters used for each system are described as follows. *Apo* MSM hyperparameters: 950 clusters, 3 tICs, a tICA lag time of 8 ns, and an MSM lag time of 30 ns. GLC MSM hyperparameters: 750 clusters, 12 tICs, a tICA lag time of 8 ns, and an MSM lag time of 20 ns. SUC MSM hyperparameters: 900 clusters, 8 tICs, a tICA lag time of 8 ns, and an MSM lag time of 40 ns (see Supplementary Table 2).

### Adaptive Sampling Error Analysis

The clustering object obtained during the MSM generation protocol was used to obtain 200 data subsets, each containing a random total of 80% of all trajectories contained within each aggregate dataset. From each of these subsets an MSM was prepared. The dimensionality of each data subset was reproportioned to the corresponding number of bins used on the reported MSM-weight free energy landscapes contained within the Main Text of this manuscript. Binning dimensions were selected for each analysis as to maintain a consistent coarse-grained resolution and visual appearance across different analysis types. The standard deviation in reported free energy differences for each of the binned data subsets was used to generate a free energy landscape for sampling error. Bootstrapped free energy error landscapes are provided for all analyses contained within main text figures in Supplementary Figures 36-38.

### Protein structure visualization

Protein snapshots were visualized and rendered in Chimera 1.14.^81^ RMSD calculations shown in Figure 2 were calculated using CPPTRAJ and inserted into representative PDB structure b-factor columns for worms representation using MDAnalysis 2.0.0.^82,83^ Structures selected for Main Text and Supplementary figures were chosen from the centers of different metastable minima along relevant free energy landscapes. Selected PDB structures used to calculate transmembrane channel pore radius were analyzed using the HOLE program.^84^

## DATA AVAILABILITY

Simulation and simulation-related results and python scripts; jupyter notebooks; MSM objects and PDBs of select AtSWEET13 conformational states are available at the Shukla Group github at the following web address: https://github.com/ShuklaGroup/AtSWEET13_sugars

## ACKNOWLEDGEMENTS

This research was part of the Blue Waters sustained-petascale computing project, was supported by the National Science Foundation (awards OCI-0725070 and ACI-1238993), the State of Illinois, as of December 2019 the National Geospatial-Intelligence Agency. During its tenure, Blue Waters was a joint effort of the University of Illinois and its National Center for Supercomputing Allocations. D.S. acknowledges support from the National Institutes of Health (Award No. R35GM142745). We thank Soumajit Dutta for useful discussions concerning MSM validation.

## AUTHOR INFORMATION

**Austin T. Weigle**

*Department of Chemistry, University of Illinois at Urbana-Champaign, Urbana, Illinois 61801, United States*

**Diwakar Shukla**

*Department of Chemical & Biomolecular Engineering; Department of Plant Biology; Department of Bioengineering; Center for Biophysics and Computational Biology, University of Illinois at Urbana-Champaign, Urbana, Illinois 61801, United States*

## Author Contributions

D.S. conceived and supervised the project. A.T.W. performed modeling and simulations. A.T.W. and D.S. analyzed the simulation data. A.T.W. wrote the manuscript with inputs from D.S., where both authors have expressed their approval of the final version of this manuscript.

## ETHICS DECLARATION

There are no competing interests.

## SUPPLEMENTARY INFORMATION

This research article contains Supplementary Information.

