## Supplementary Materials for "AtSWEET13 transporter discriminates sugars by selective facial and positional substrate recognition"

### Table of Contents

#### Figures and Tables


Figure S1 tICA decomposition given simplified MSM features for AtSWEET13 transport processes S-6

Figure S2 Scatterplot representation of transport versus the difference in gating highlights that substrate transport proceeds through an hour-glass state S-7

Figure S3 MSM-weighted intracellular minus extracellular gating distance versus AtSWEET13 transmembrane channel Z position of the closest GLC molecule carbon atom to the W58-W180 binding pocket S-8

Figure S4 MSM-weighted intracellular minus extracellular gating distance versus AtSWEET13 transmembrane channel Z position of the closest GLC molecule oxygen atom to the W58-W180 binding pocket S-9

Figure S5 MSM-weighted intracellular minus extracellular gating distance versus AtSWEET13 transmembrane channel Z position of the closest SUC molecule glucosyl carbon atom to the W58-W180 binding pocket S-10

Figure S6 MSM-weighted intracellular minus extracellular gating distance versus AtSWEET13 transmembrane channel Z position of the closest SUC molecule glucosyl oxygen atom to the W58-W180 binding pocket S-11

Figure S7 MSM-weighted intracellular minus extracellular gating distance versus AtSWEET13 transmembrane channel Z position of the closest SUC molecule fructosyl carbon atom to the W58-W180 binding pocket S-12

Figure S8 MSM-weighted intracellular minus extracellular gating distance versus AtSWEET13 transmembrane channel Z position of the closest SUC molecule fructosyl oxygen atom to the W58-W180 binding pocket S-13

Figure S9 AtSWEET13 transport cycles for GLC and SUC translocation implicate minimal conformational change outside of commitment to alternate access S-14

Figure S10 Similar AtSWEET13 pore radius aperture is maintained regardless of substrate transported S-15

Figure S11 Raw counts versus MSM population in each clustered state S-16

Figure S12 Unweighted feature-diverse tICA decomposition landscapes S-17

Figure S13 Descriptor correlation to tIC1 from feature-diverse tICA decomposition of Apo transport S-18

Figure S14 Descriptor correlation to tIC1 from feature-diverse tICA decomposition of GLC transport S-18

Figure S15 Descriptor correlation to tIC1 from feature-diverse tICA decomposition of SUC transport S-19

Figure S16 Descriptor correlation to tIC2 from feature-diverse tICA decomposition of GLC transport S-20

Figure S17 Descriptor correlation to tIC2 from feature-diverse tICA decomposition of SUC transport S-21

Figure S18 MSM-weighted $\boldsymbol{\theta}_{xy}$ analysis versus AtSWEET13 transmembrane channel Z position of the closest GLC molecule carbon atom to the W58-W180 binding pocket S-22

Figure S19 MSM-weighted $\boldsymbol{\theta}_{xy}$ analysis versus AtSWEET13 transmembrane channel Z position of the closest GLC molecule oxygen atom to the W58-W180 binding pocket S-23

Figure S20 MSM-weighted $\boldsymbol{\theta}_{xy}$ analysis versus AtSWEET13 transmembrane channel Z position of the closest SUC molecule glucosyl carbon atom to the W58-W180 binding pocket S-24

Figure S21 MSM-weighted $\boldsymbol{\theta}_{xy}$ analysis versus AtSWEET13 transmembrane channel Z position of the closest SUC molecule glucosyl oxygen atom to the W58-W180 binding pocket S-25

Figure S22 MSM-weighted $\boldsymbol{\theta}_{xy}$ analysis versus AtSWEET13 transmembrane channel Z position of the closest SUC molecule fructosyl carbon atom to the W58-W180 binding pocket S-26

Figure S23 MSM-weighted $\boldsymbol{\theta}_{xy}$ analysis versus AtSWEET13 transmembrane channel Z position of the closest SUC molecule fructosyl oxygen atom to the W58-W180 binding pocket S-27

Figure S24 MSM-weighted $\boldsymbol{\theta}_{xz}$ analysis versus AtSWEET13 transmembrane channel Z position of the closest GLC molecule carbon atom to the W58-W180 binding pocket S-28

Figure S25 MSM-weighted $\boldsymbol{\theta}_{xz}$ analysis versus AtSWEET13 transmembrane channel Z position of the closest GLC molecule oxygen atom to the W58-W180 binding pocket S-29

Figure S26 MSM-weighted $\boldsymbol{\theta}_{xz}$ analysis versus AtSWEET13 transmembrane channel Z position of the closest SUC molecule glucosyl carbon atom to the W58-W180 binding pocket S-30

Figure S27 MSM-weighted $\theta_{xz}$ analysis versus AtSWEET13 transmembrane channel Z position of the closest SUC molecule glucosyl oxygen atom to the W58-W180 binding pocket S-31

Figure S28 MSM-weighted $\theta_{xz}$ analysis versus AtSWEET13 transmembrane channel Z position of the closest SUC molecule fructosyl carbon atom to the W58-W180 binding pocket S-32

Figure S29 MSM-weighted $\theta_{xz}$ analysis versus AtSWEET13 transmembrane channel Z position of the closest SUC molecule fructosyl oxygen atom to the W58-W180 binding pocket S-33

Figure S30 MSM-weighted $\theta_{yz}$ analysis versus AtSWEET13 transmembrane channel Z position of the closest GLC molecule carbon atom to the W58-W180 binding pocket S-34

Figure S31 MSM-weighted $\theta_{yz}$ analysis versus AtSWEET13 transmembrane channel Z position of the closest GLC molecule oxygen atom to the W58-W180 binding pocket S-35

Figure S32 MSM-weighted $\theta_{yz}$ analysis versus AtSWEET13 transmembrane channel Z position of the closest SUC molecule glucosyl carbon atom to the W58-W180 binding pocket S-36

Figure S33 MSM-weighted $\theta_{yz}$ analysis versus AtSWEET13 transmembrane channel Z position of the closest SUC molecule glucosyl oxygen atom to the W58-W180 binding pocket S-37

Figure S34 MSM-weighted $\theta_{yz}$ analysis versus AtSWEET13 transmembrane channel Z position of the closest SUC molecule fructosyl carbon atom to the W58-W180 binding pocket S-38

Figure S35 MSM-weighted $\theta_{yz}$ analysis versus AtSWEET13 transmembrane channel Z position of the closest SUC molecule fructosyl oxygen atom to the W58-W180 binding pocket S-39

Figure S36 GLC and SUC structures with highlighted functional considered critical for molecular recognition by AtSWEET13 S-40

Figure S37 MSM-weighted bootstrap error plots for adaptive sampling of gating landscapes S-41

Figure S38 MSM-weighted bootstrap error plots for adaptive sampling of intracellular minus extracellular gating distance versus AtSWEET13 transmembrane channel ligand Z position landscapes for sugar transport S-42

Figure S39 MSM-weighted bootstrap error plots for adaptive sampling of θ rotation analyses presented in Main Text Figure 3 S-43

Figure S40 Implied timescale plots calculated with Bayesian error for AtSWEET13 transport processes S-44

Table S1 Finalized features used for MSM discretization S-45

Table S2 Finalized MSM hyperparameters S-46

Table S3 Feature-diverse tICA decomposition features and correlations – Apo transport S-47

Table S4 Feature-diverse tICA features and correlations – SUC transport S-51

Table S5 Feature-diverse tICA features and correlations – GLC transport S-84

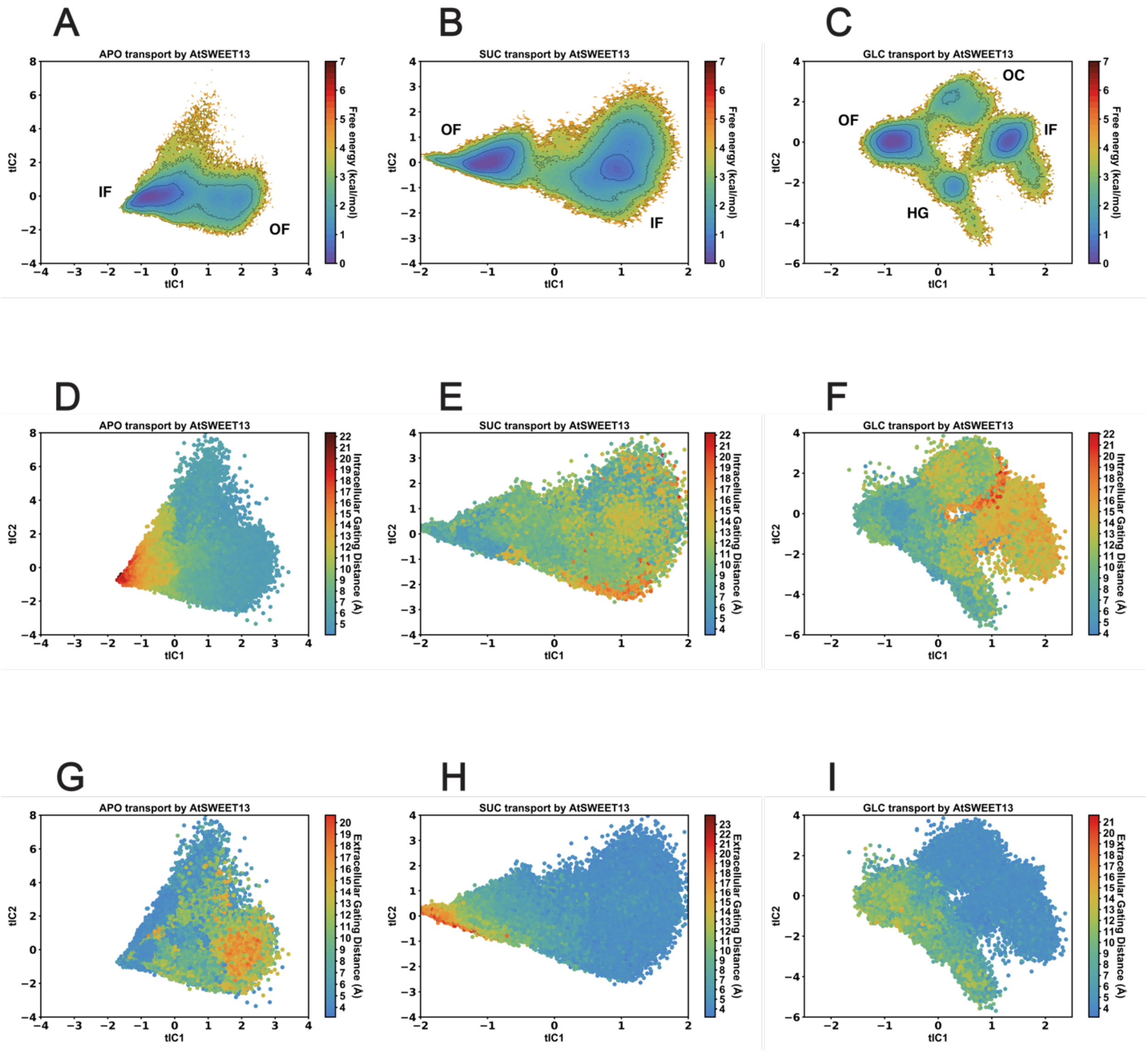


**Figure S1.** tICA decomposition given simplified MSM features for AtSWEET13 transport processes. (A) Apo, (B) SUC, and (C) GLC MSM-weighted tICA transport landscapes. tICA decomposition represented as scatterplots are colored by extent of intracellular gating distance for (D) Apo, (E) SUC, and (F) GLC transport processes. tICA decomposition represented as scatterplots are colored by extent of extracellular gating distance for (G) Apo, (H) SUC, and (I) GLC transport processes.

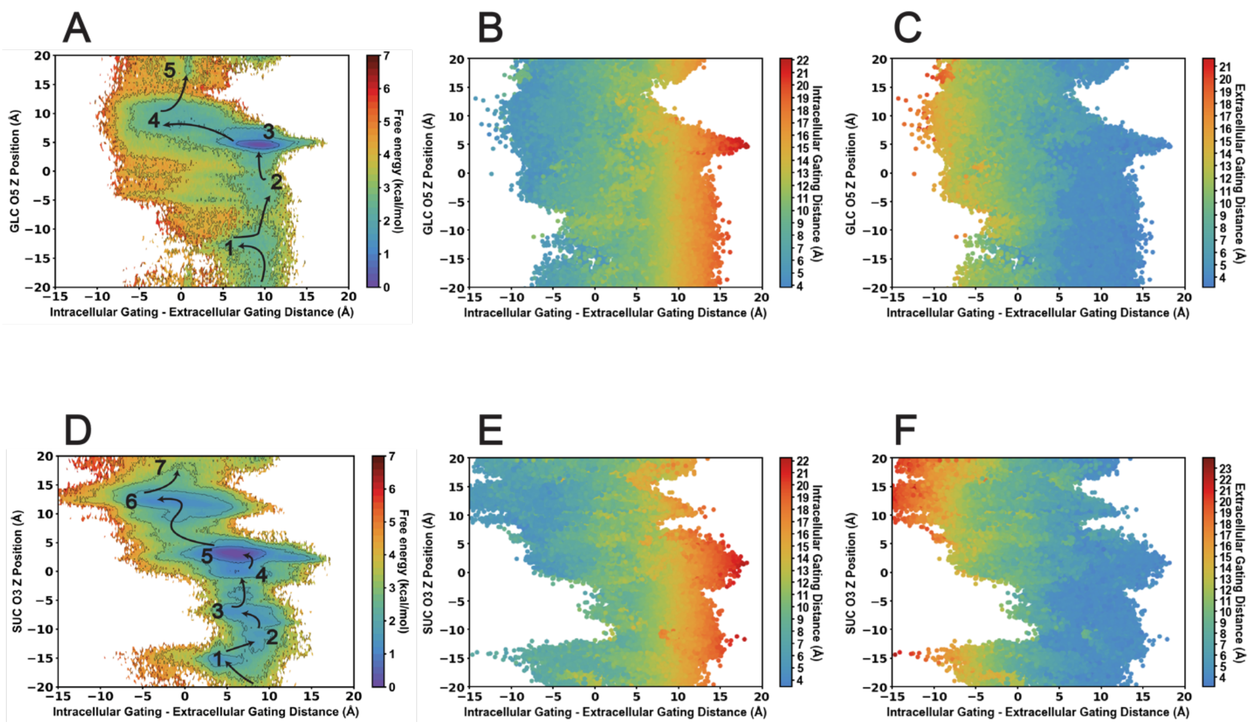


**Figure S2.** Scatterplot representation of transport versus the difference in gating highlights that substrate transport proceeds through an hour-glass state. GLC transport versus the difference in intracellular versus extracellular gating represented using (A) an MSM-weighted free energy landscape, (B) a scatterplot colored by extent of intracellular gating, and (C) a scatterplot colored by extent of extracellular gating. SUC transport versus the difference in intracellular versus extracellular gating represented using (D) an MSM-weighted free energy landscape, (E) a scatterplot colored by extent of intracellular gating, and (F) a scatterplot colored by extent of extracellular gating.

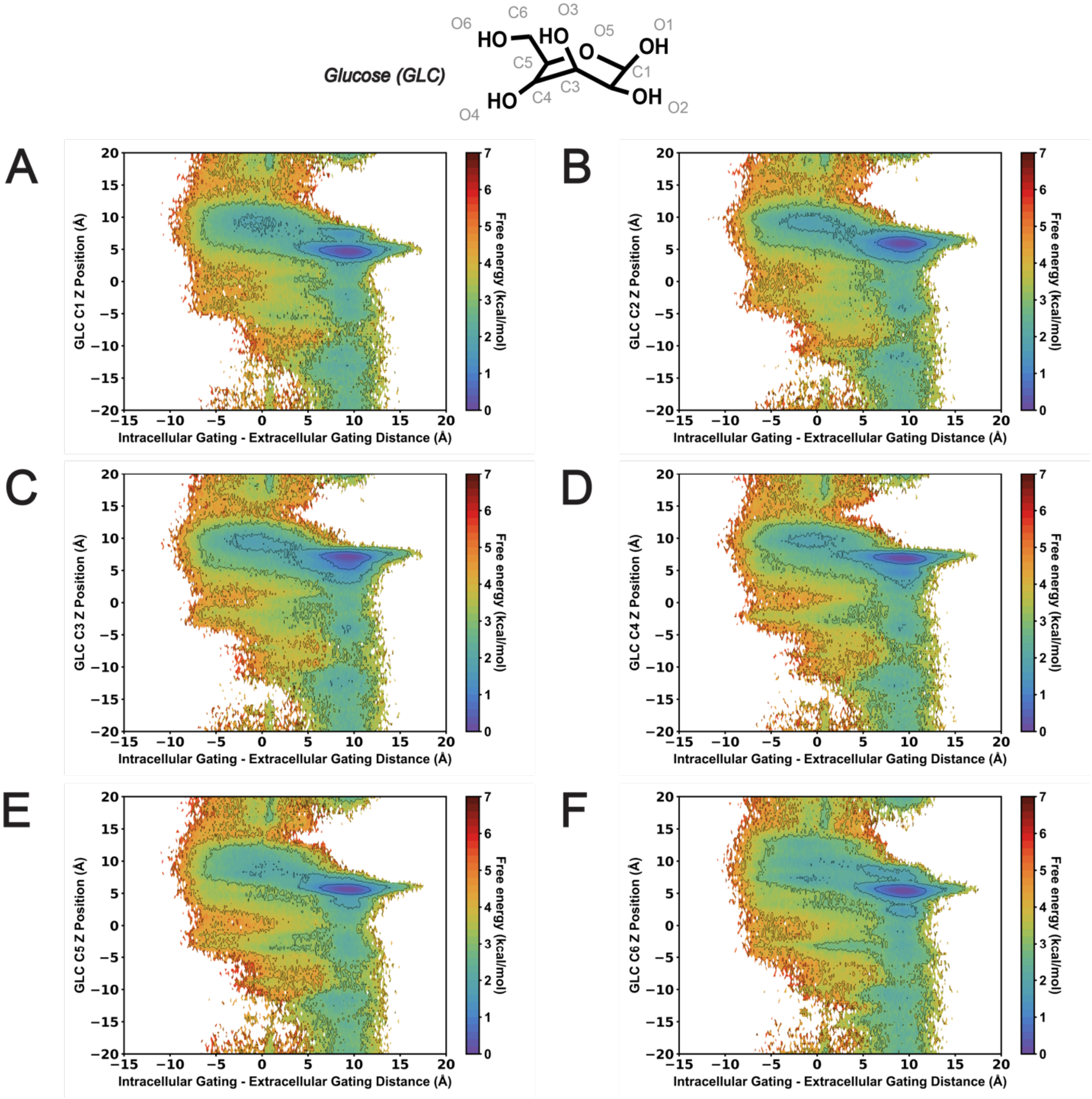


**Figure S3.** MSM-weighted intracellular minus extracellular gating distance versus AtSWEET13 transmembrane channel Z position of the closest GLC molecule carbon atom to the W58-W180 binding pocket. (A) GLC C1. (B) GLC C2. (C) GLC C3. (D) GLC C4. (E) GLC C5. (F) GLC C6.

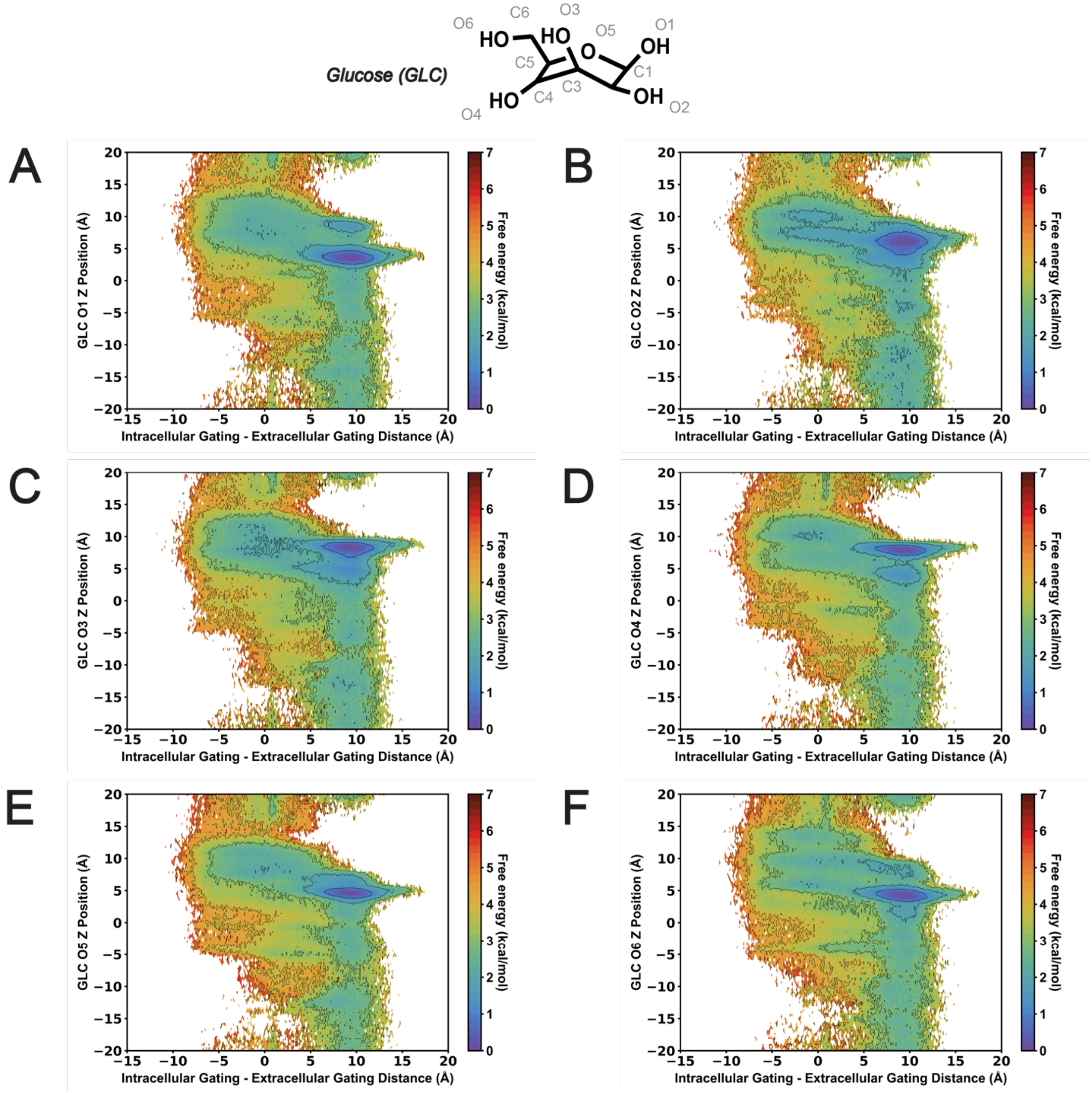


**Figure S4.** MSM-weighted intracellular minus extracellular gating distance versus AtSWEET13 transmembrane channel Z position of the closest GLC molecule oxygen atom to the W58-W180 binding pocket. (A) GLC O1. (B) GLC O2. (C) GLC O3. (D) GLC O4. (E) GLC O5. (F) GLC O6.

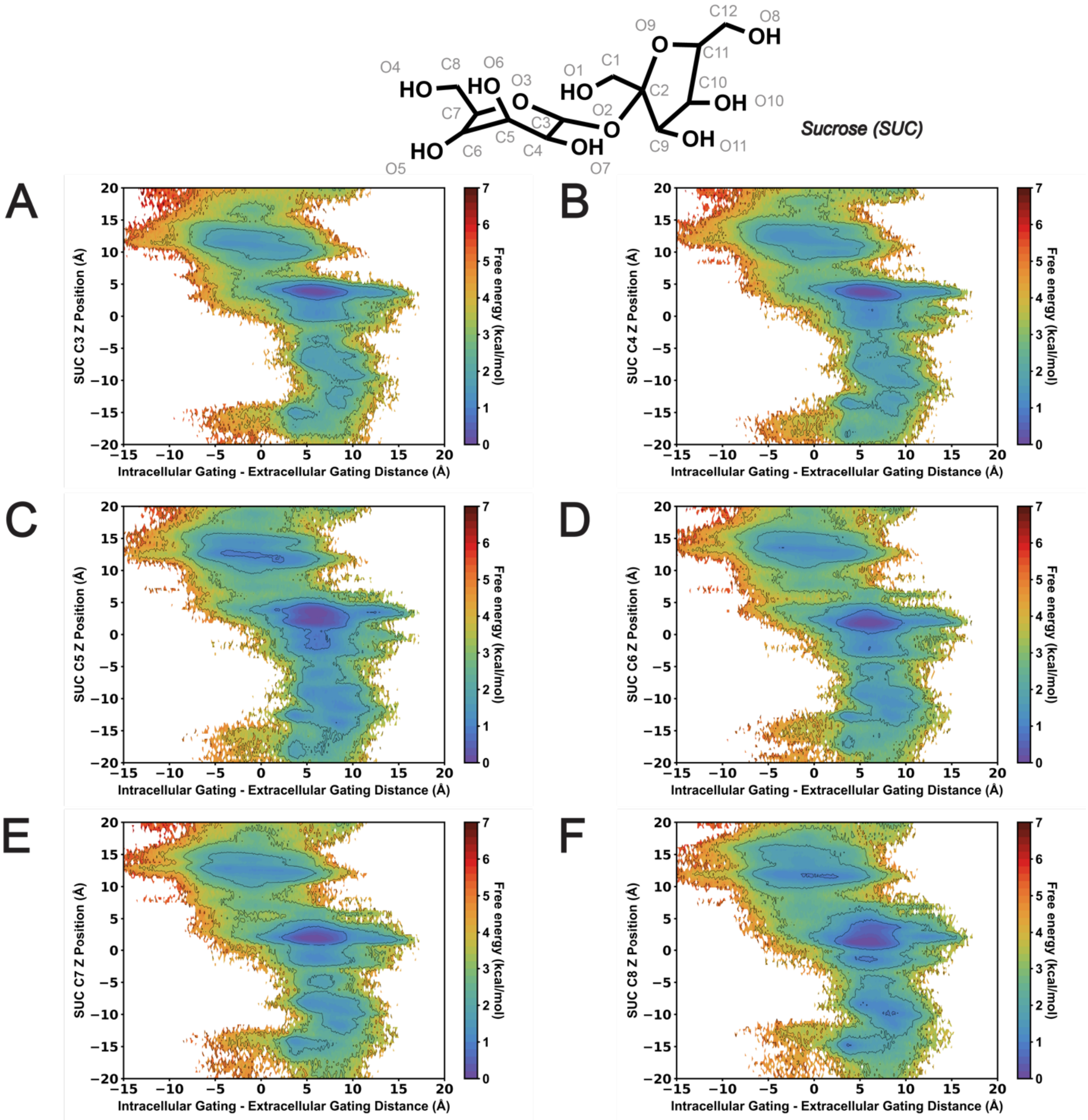


**Figure S5.** MSM-weighted intracellular minus extracellular gating distance versus AtSWEET13 transmembrane channel Z position of the closest SUC molecule glucosyl carbon atom to the W58-W180 binding pocket. (A) SUC C3. (B) SUC C4. (C) SUC C5. (D) SUC C6. (E) SUC C7. (F) SUC C8.

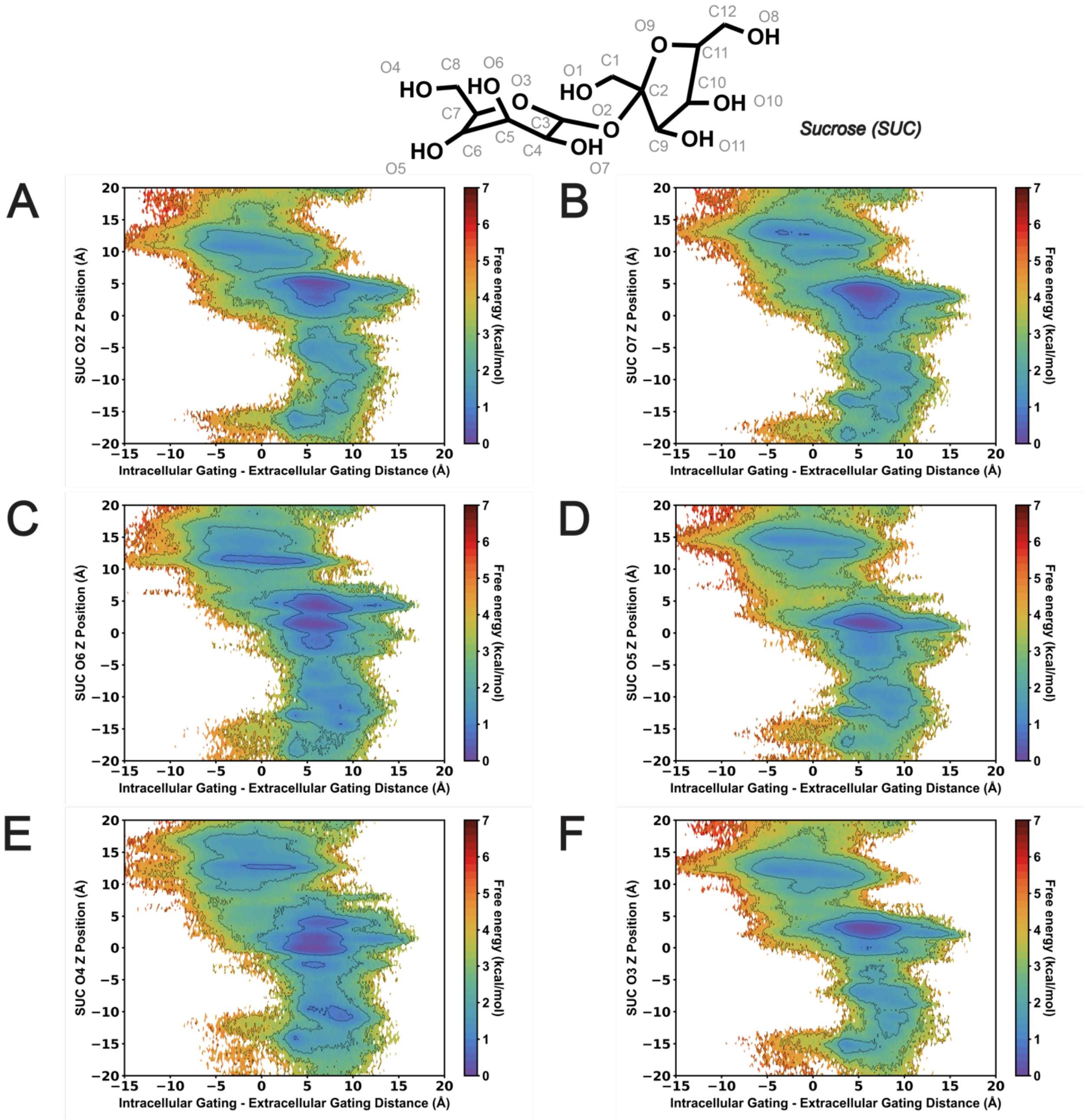


**Figure S6.** MSM-weighted intracellular minus extracellular gating distance versus AtSWEET13 transmembrane channel Z position of the closest SUC molecule glucosyl oxygen atom to the W58-W180 binding pocket. (A) SUC O2. (B) SUC O7. (C) SUC O6. (D) SUC O5. (E) SUC O4. (F) SUC O3.

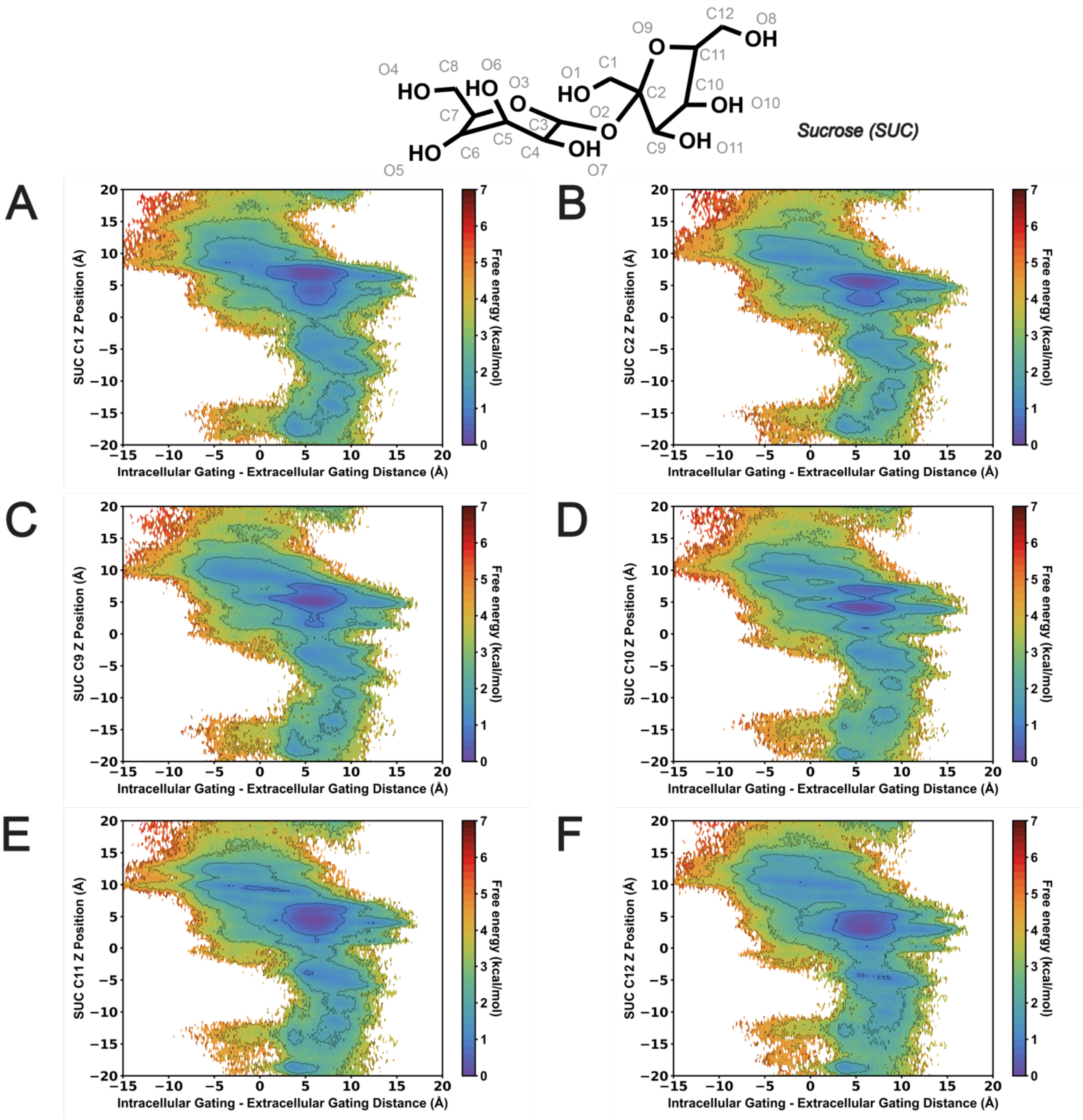


**Figure S7.** MSM-weighted intracellular minus extracellular gating distance versus AtSWEET13 transmembrane channel Z position of the closest SUC molecule fructosyl carbon atom to the W58-W180 binding pocket. (A) SUC C1. (B) SUC C2. (C) SUC C9. (D) SUC C10. (E) SUC C11. (F) SUC C12.

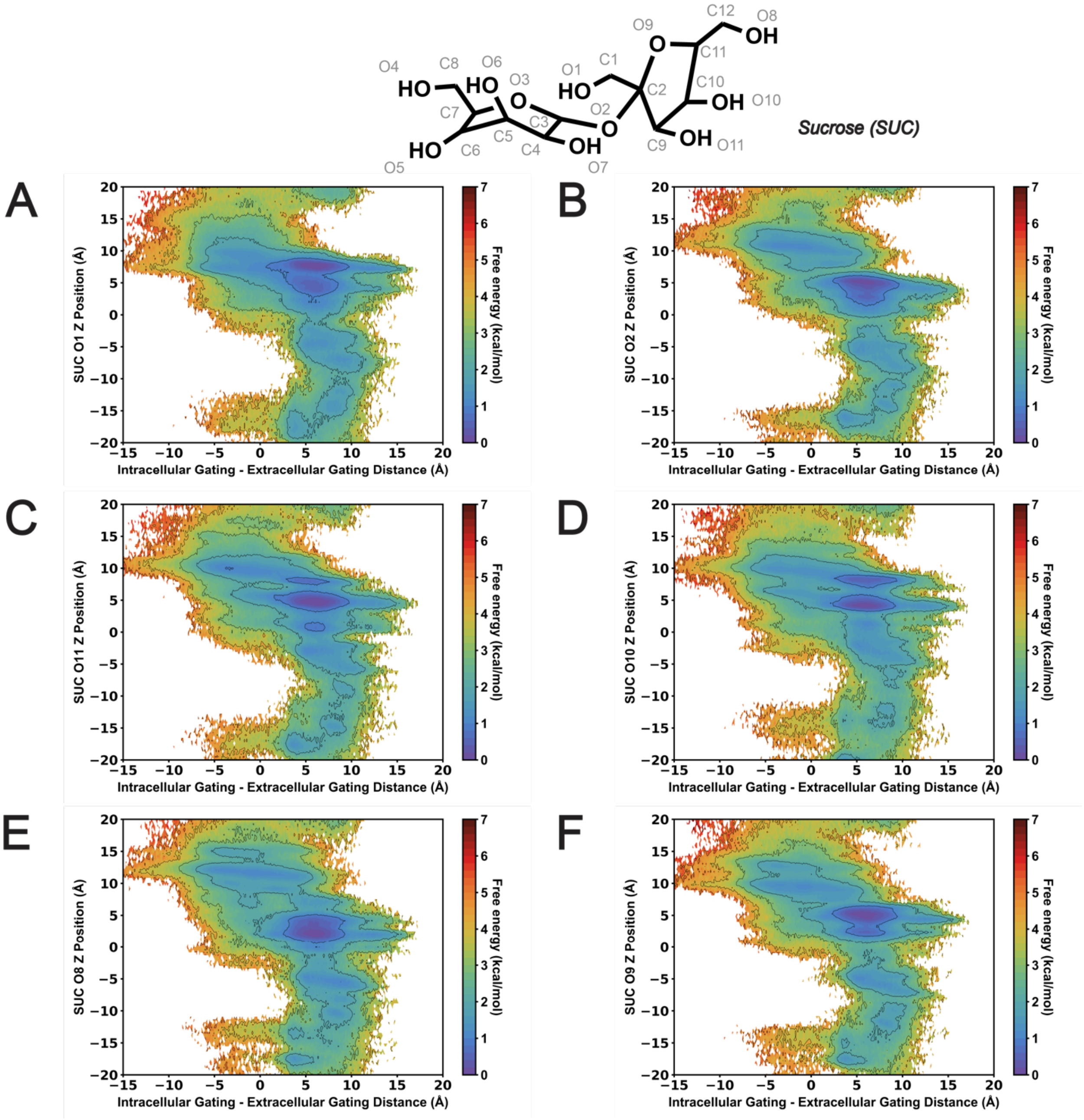


**Figure S8.** MSM-weighted intracellular minus extracellular gating distance versus AtSWEET13 transmembrane channel Z position of the closest SUC molecule fructosyl oxygen atom to the W58-W180 binding pocket. (A) SUC O1. (B) SUC O2. (C) SUC O11. (D) SUC O10. (E) SUC O8. (F) SUC O9.

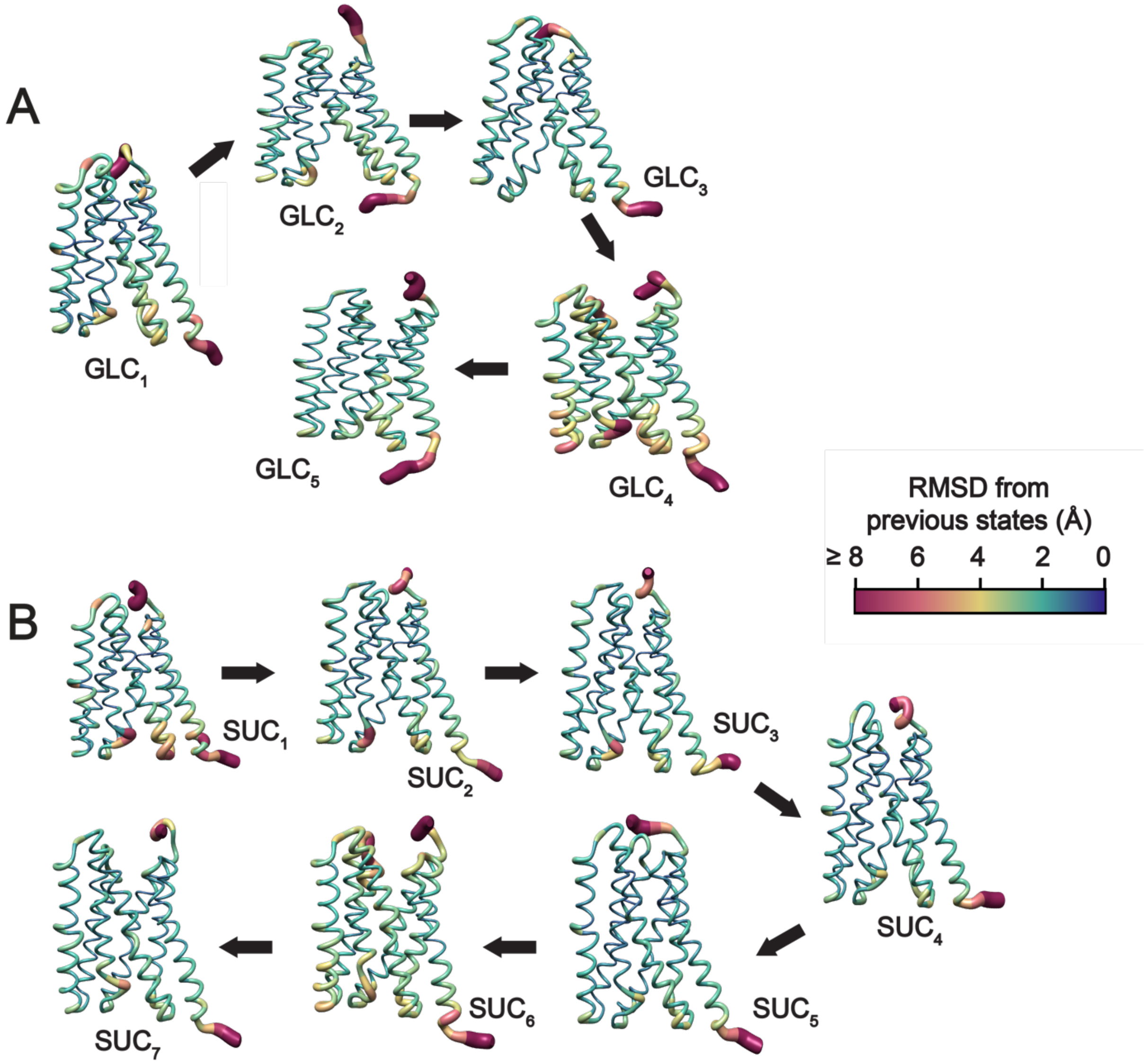


**Figure S9.** AtSWEET13 transport cycles for GLC and SUC translocation implicate minimal conformational change outside of commitment to alternate access. (A) Protein snapshots taken for different conformational states depicting GLC transport, where worms visualization represents protein residue RMSD values in Å units. (B) Protein snapshots taken for different conformational states depicting SUC transport using similar worms visualization as seen in (A). RMSDs are calculated as an average in comparison to the number of states found at the center of each energetic minima characterizing each enumerated state in Main Text Figure 2. RMSDs shown for states $GLC_1$ and $SUC_1$ are in comparison to the 5XPD crystal structure conformation.

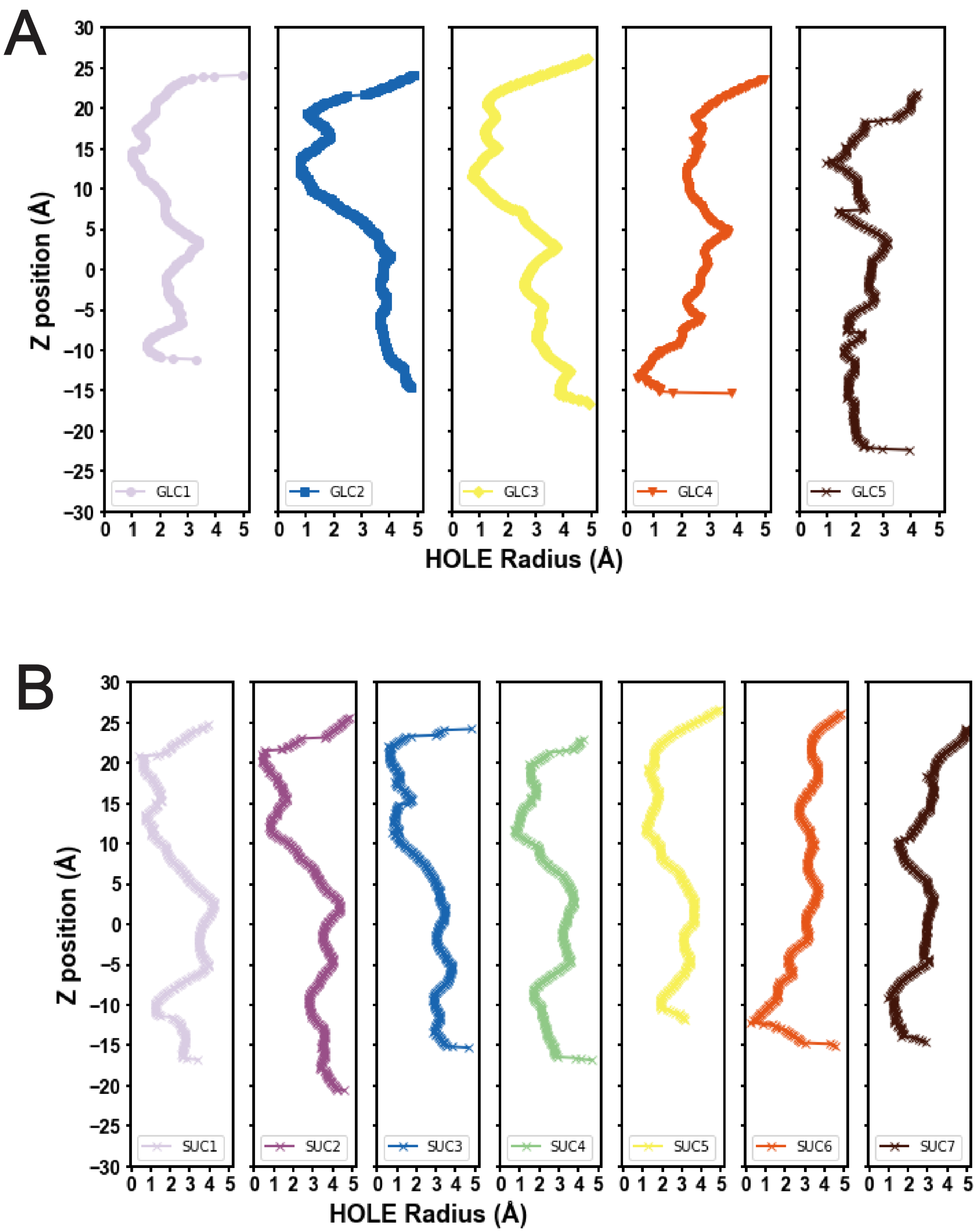


**Figure S10**. Similar AtSWEET13 pore radius aperture is maintained regardless of substrate transported. (A) HOLE calculations of metastable states observed for GLC transport. (B) HOLE calculations of metastable states observed for SUC transport. State numbering is identical to states shown in Main Text Figure 2. GLC HOLE calculations shown here in Panel A correspond to the state numbering shown in Main Text Figure 2A and Figure S9A. SUC HOLE calculations shown here in Panel B correspond to the state numbering shown in Main Text Figure 2B and Figure S9B.

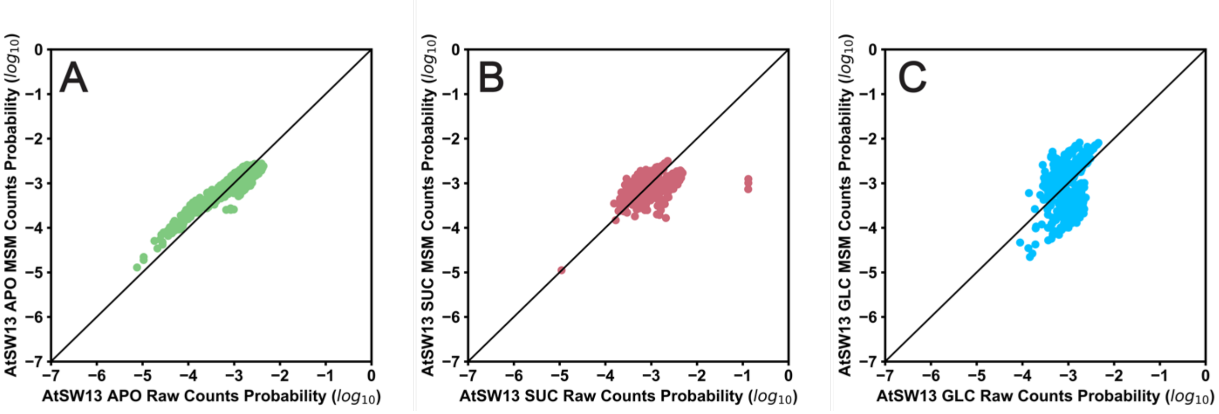


**Figure S11.** Raw counts versus MSM population in each clustered state for (A) Apo, (B) SUC, and (C) GLC transport by AtSWEET13.

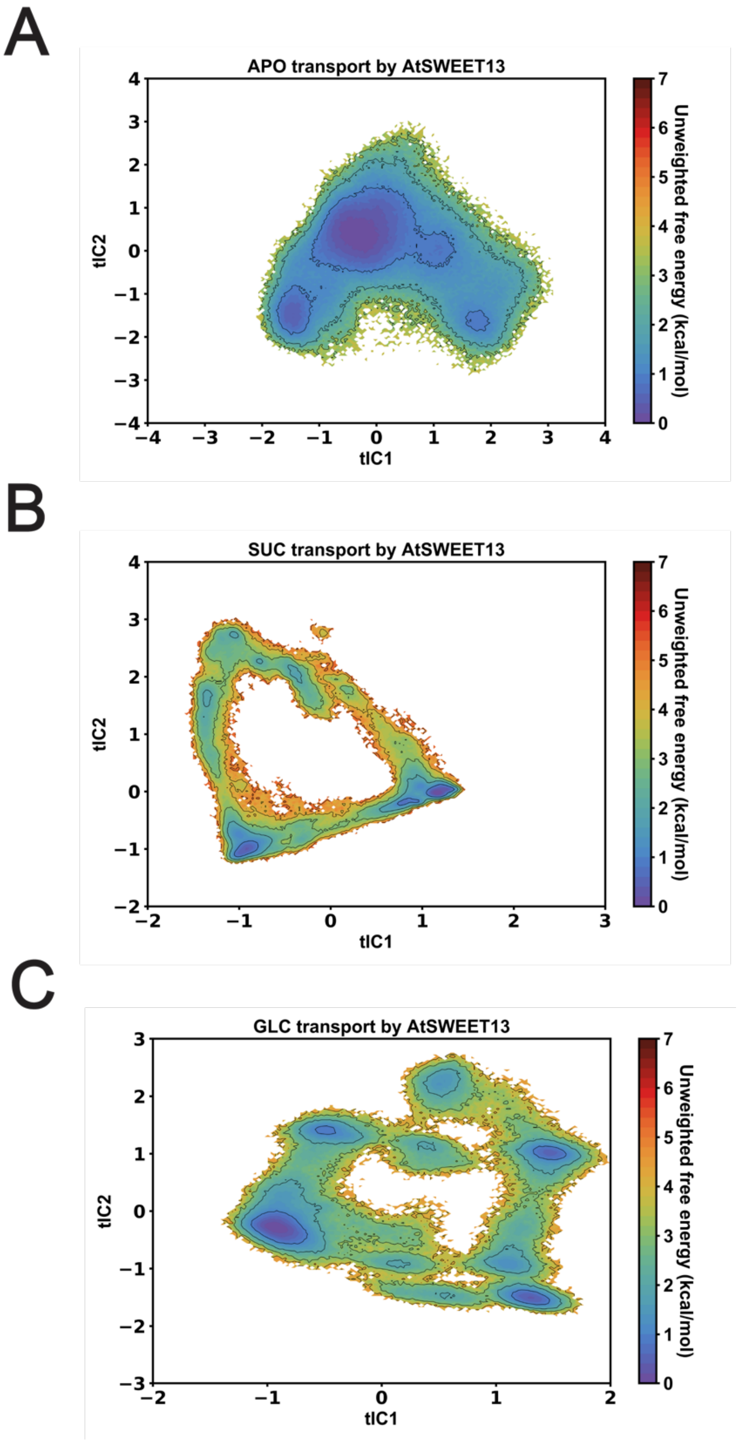


**Figure S12.** Unweighted feature-diverse tICA decomposition landscapes for (A) Apo, (B) SUC, and (C) GLC transport by AtSWEET13.

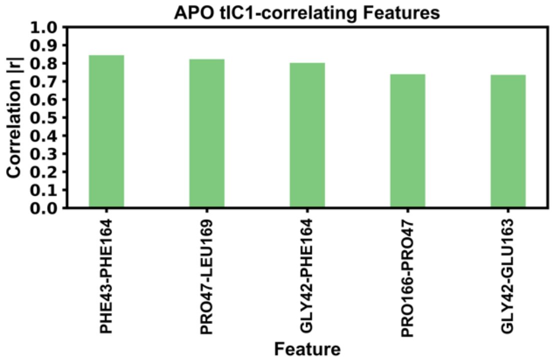


**Figure S13.** Descriptor correlation to tIC1 from feature-diverse tICA decomposition of Apo transport.

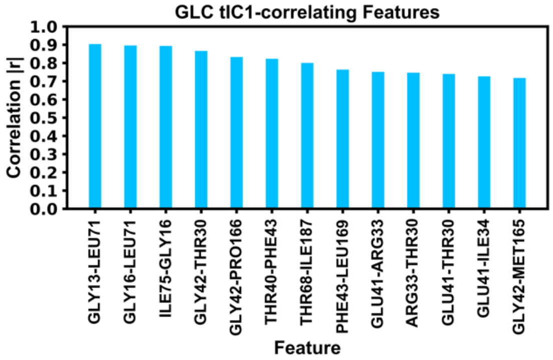


**Figure S14.** Descriptor correlation to tIC1 from feature-diverse tICA decomposition of GLC transport.

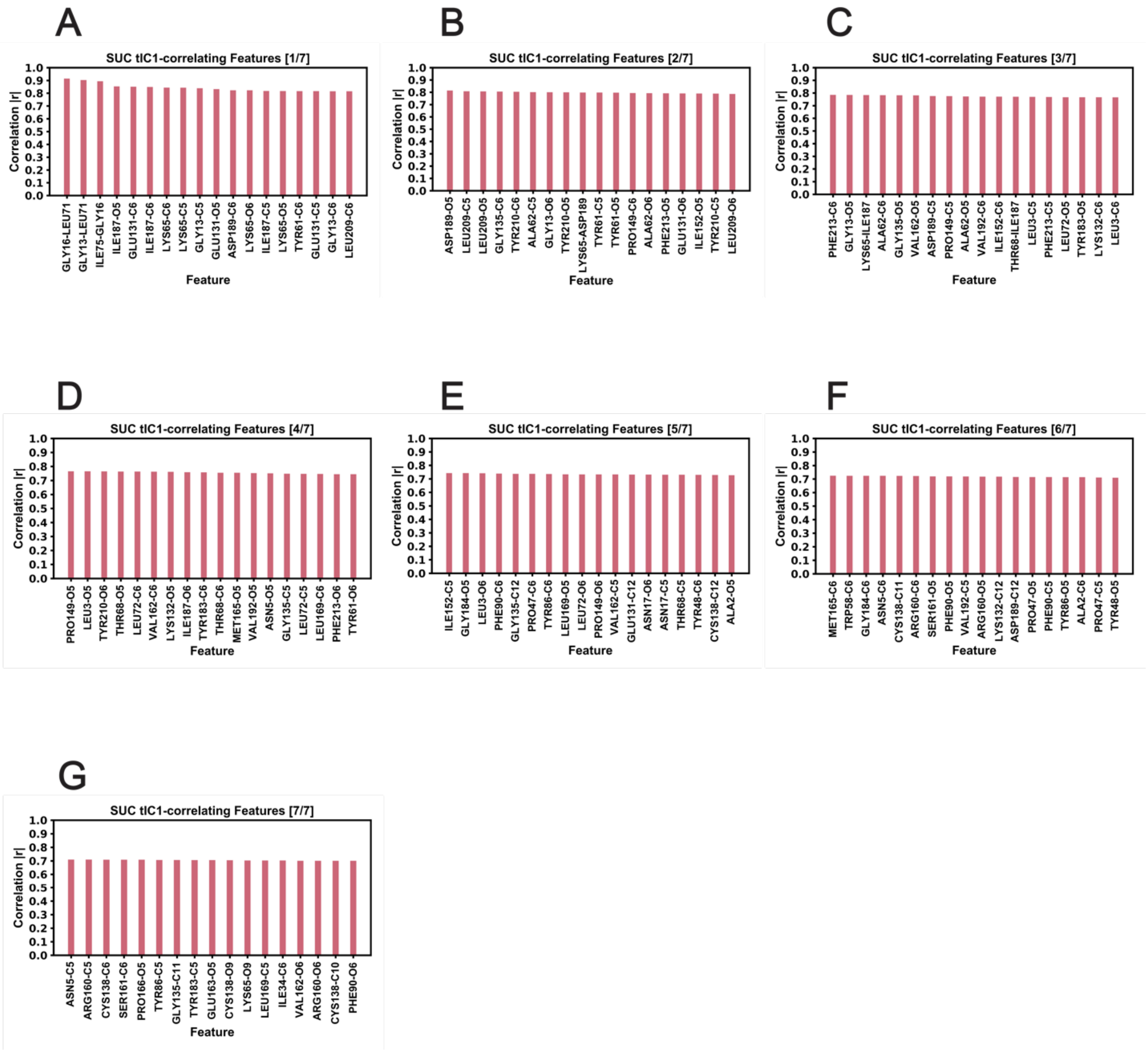


**Figure S15.** Descriptor correlation to tIC1 from feature-diverse tICA decomposition of SUC transport.

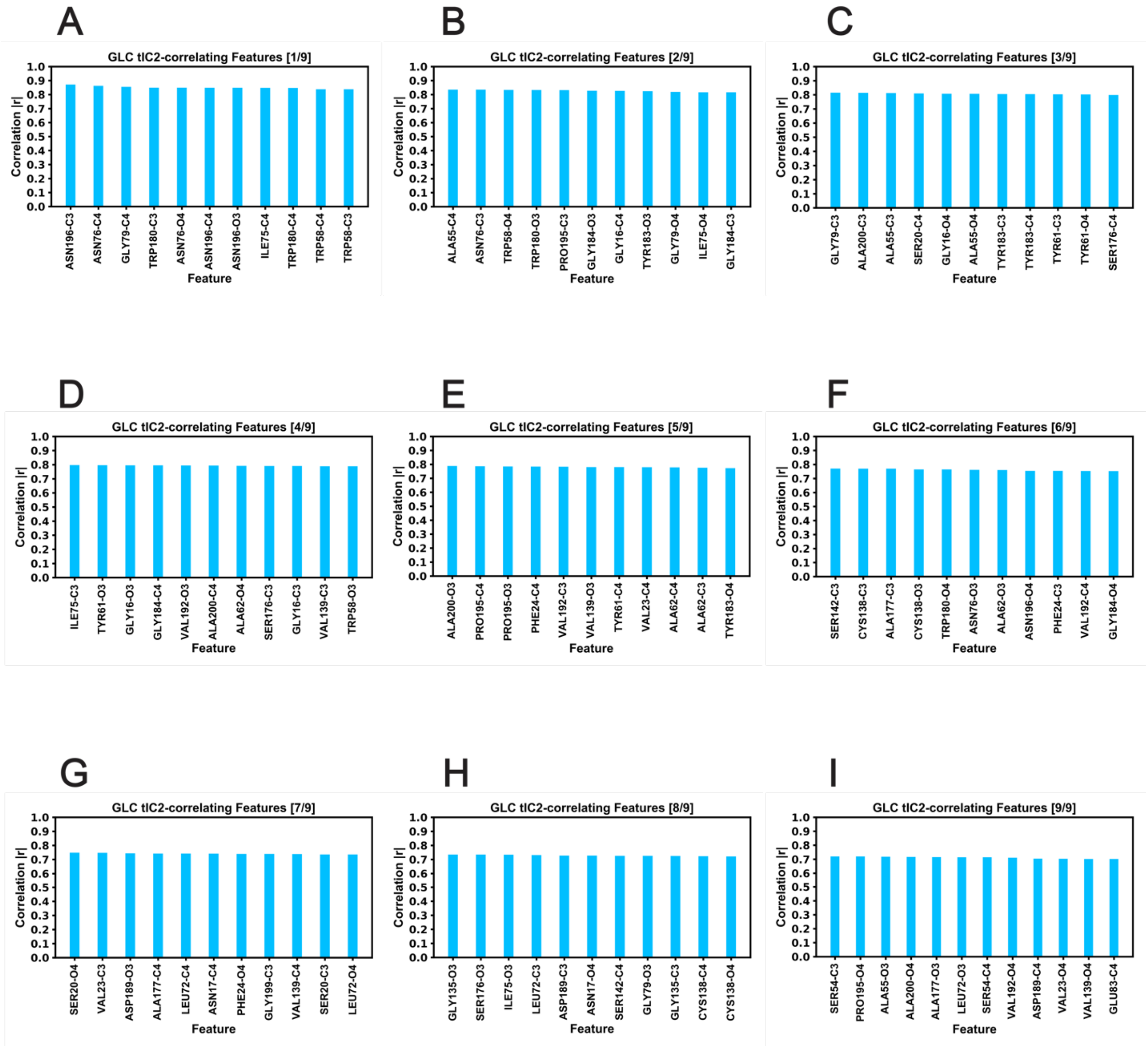


**Figure S16.** Descriptor correlation to tIC2 from feature-diverse tICA decomposition of GLC transport.

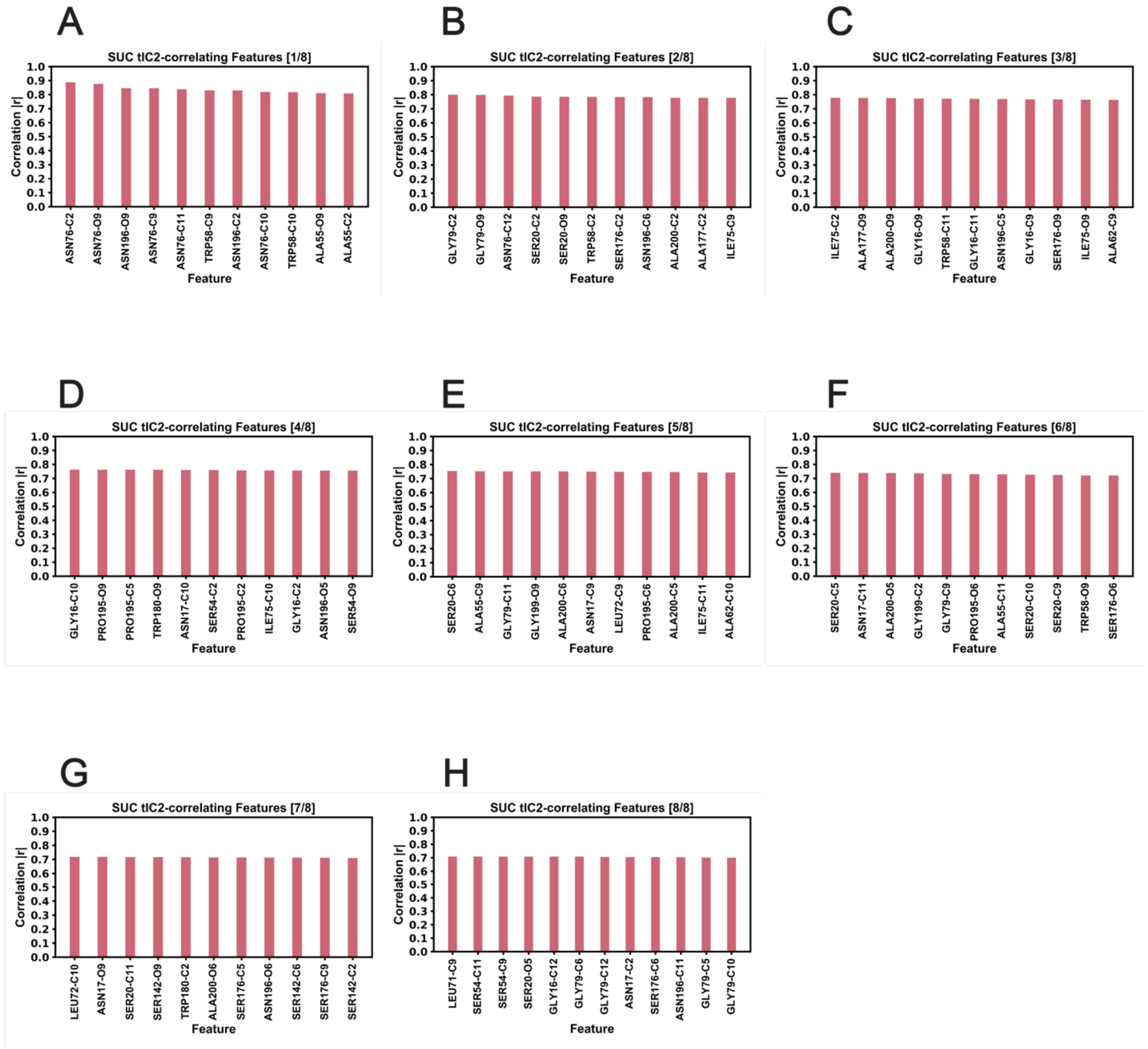


**Figure S17.** Descriptor correlation to tIC2 from feature-diverse tICA decomposition of SUC transport.

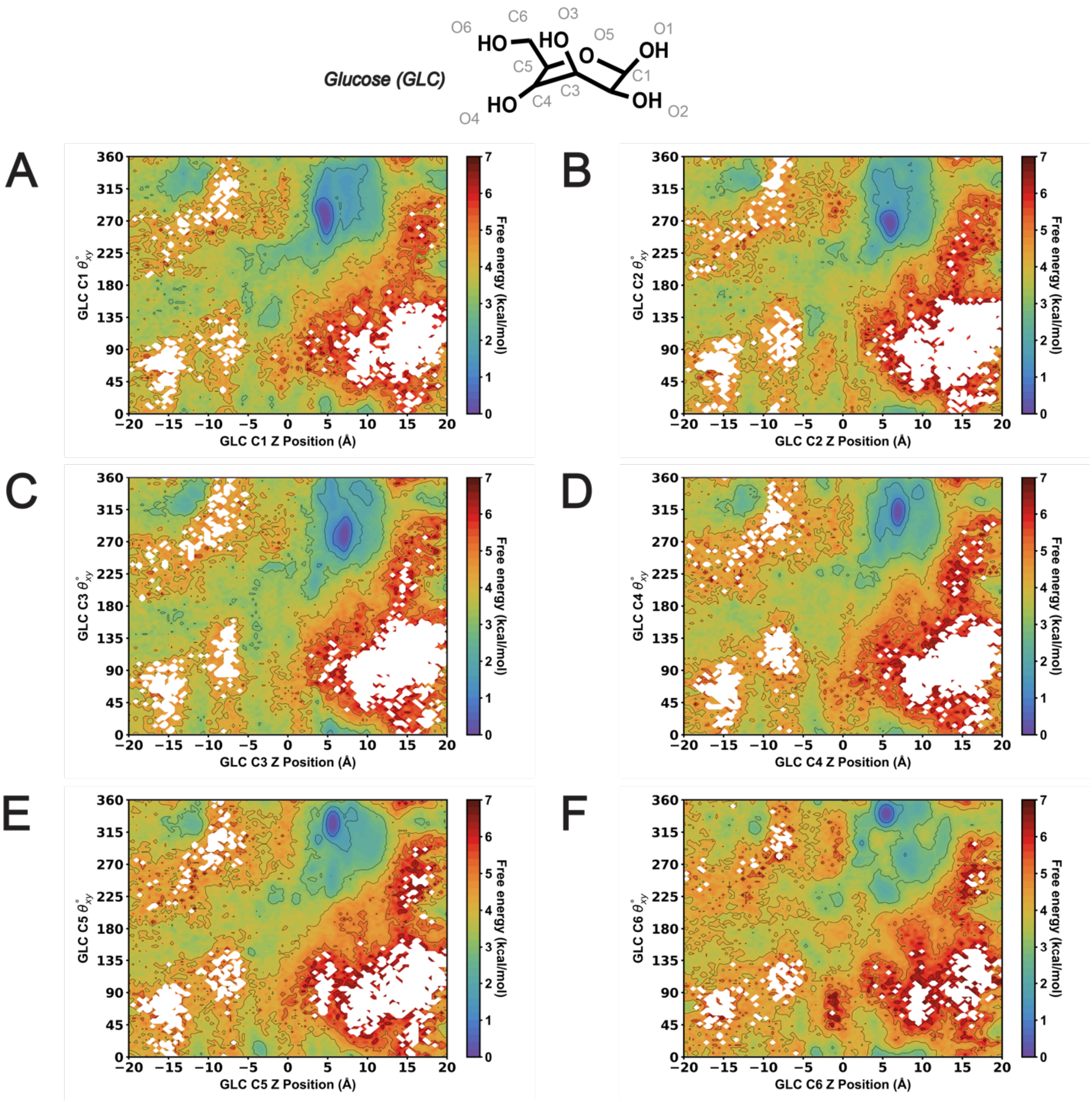


**Figure S18.** MSM-weighted $\theta_{xy}$ analysis versus AtSWEET13 transmembrane channel Z position of the closest GLC molecule carbon atom to the W58-W180 binding pocket. (A) GLC C1. (B) GLC C2 (C) GLC C3. (D) GLC C4. (E) GLC C5. (F) GLC C6.

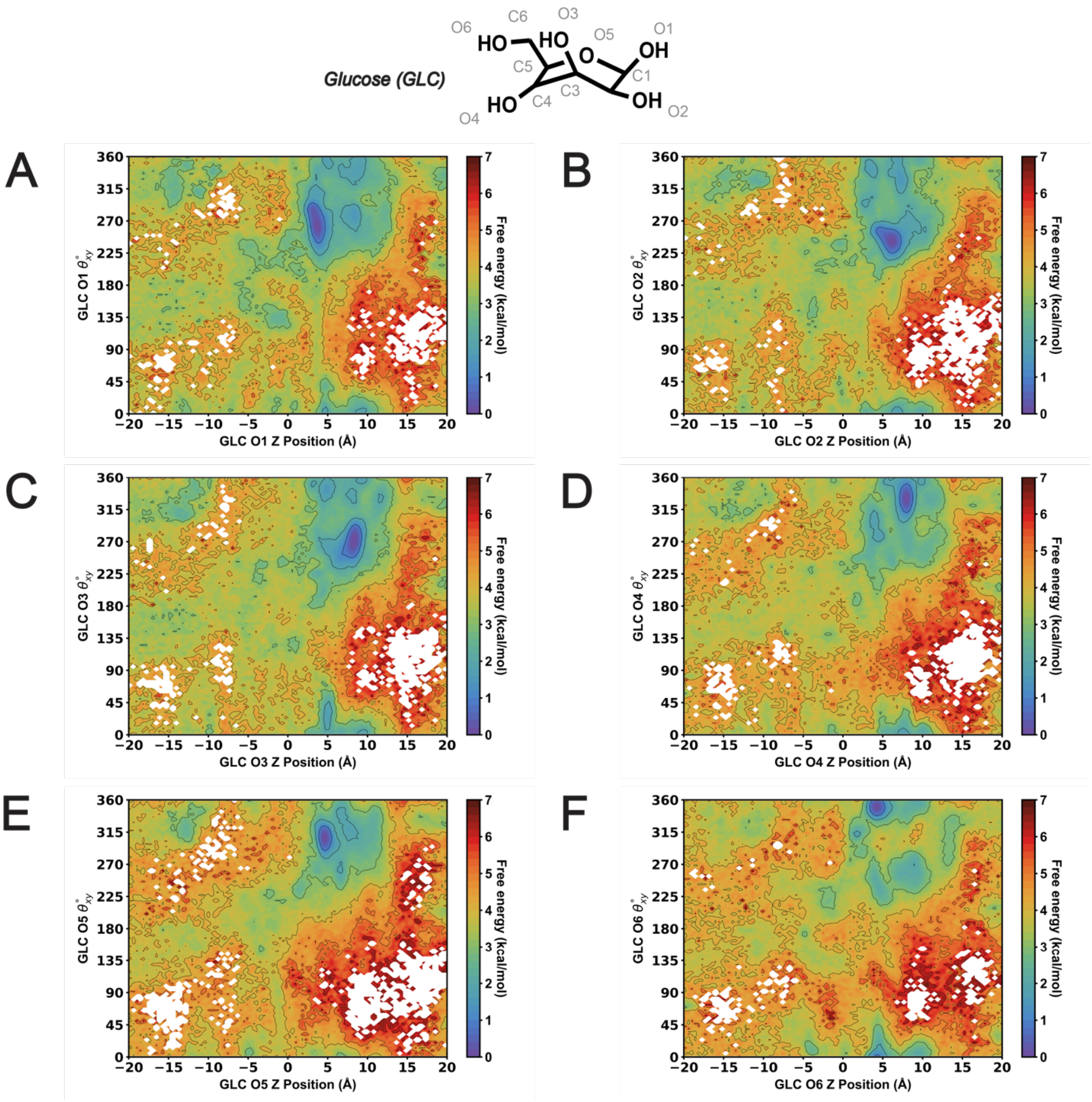


**Figure S19.** MSM-weighted $\theta_{xy}$ analysis versus AtSWEET13 transmembrane channel Z position of the closest GLC molecule oxygen atom to the W58-W180 binding pocket. (A) GLC O1. (B) GLC O2 (C) GLC O3. (D) GLC O4. (E) GLC O5. (F) GLC O6.

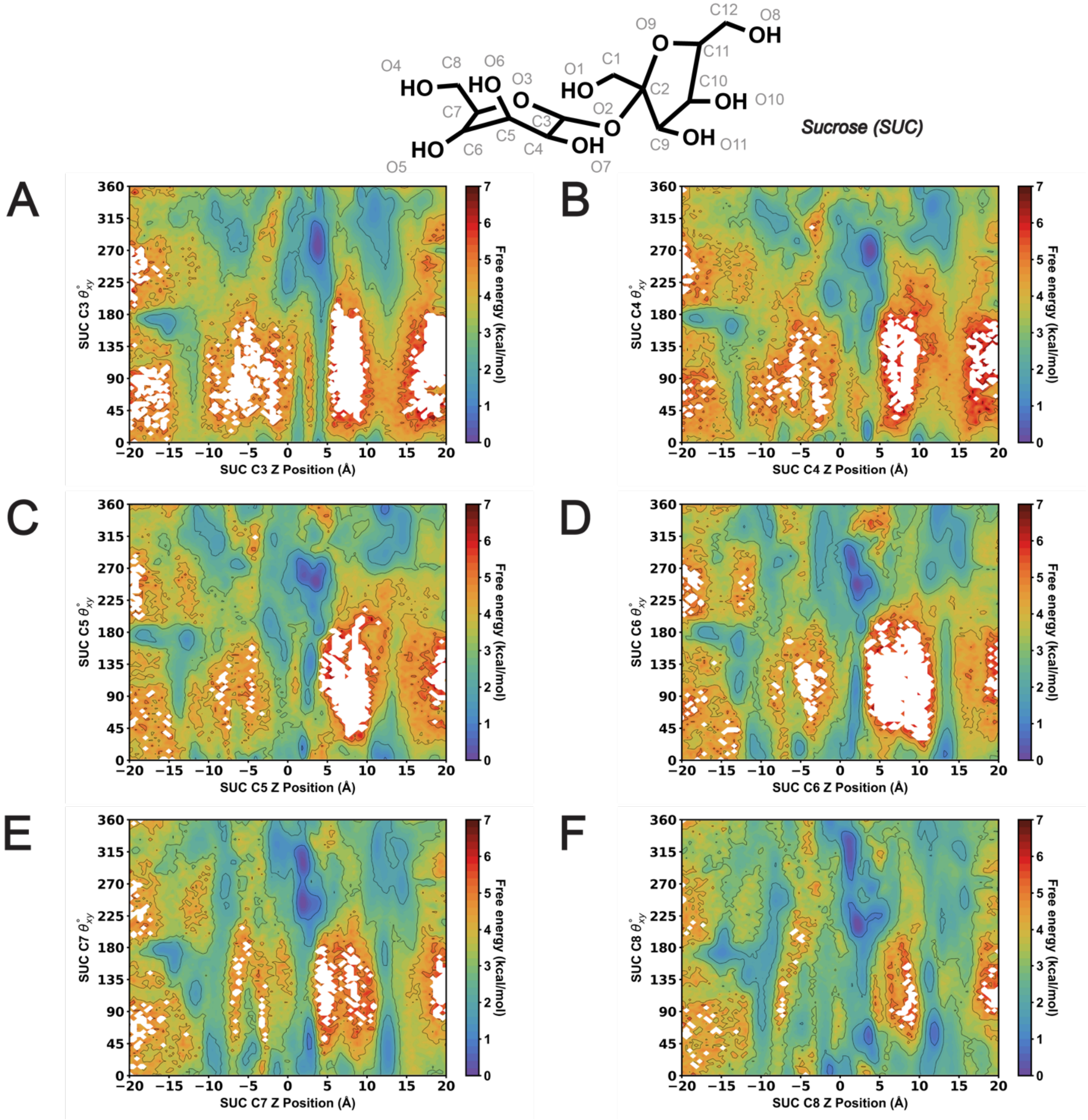


**Figure S20.** MSM-weighted $\theta_{xy}$ analysis versus AtSWEET13 transmembrane channel Z position of the closest SUC molecule glucosyl carbon atom to the W58-W180 binding pocket. (A) SUC C3. (B) SUC C4 (C) SUC C5. (D) SUC C6. (E) SUC C7. (F) SUC C8.

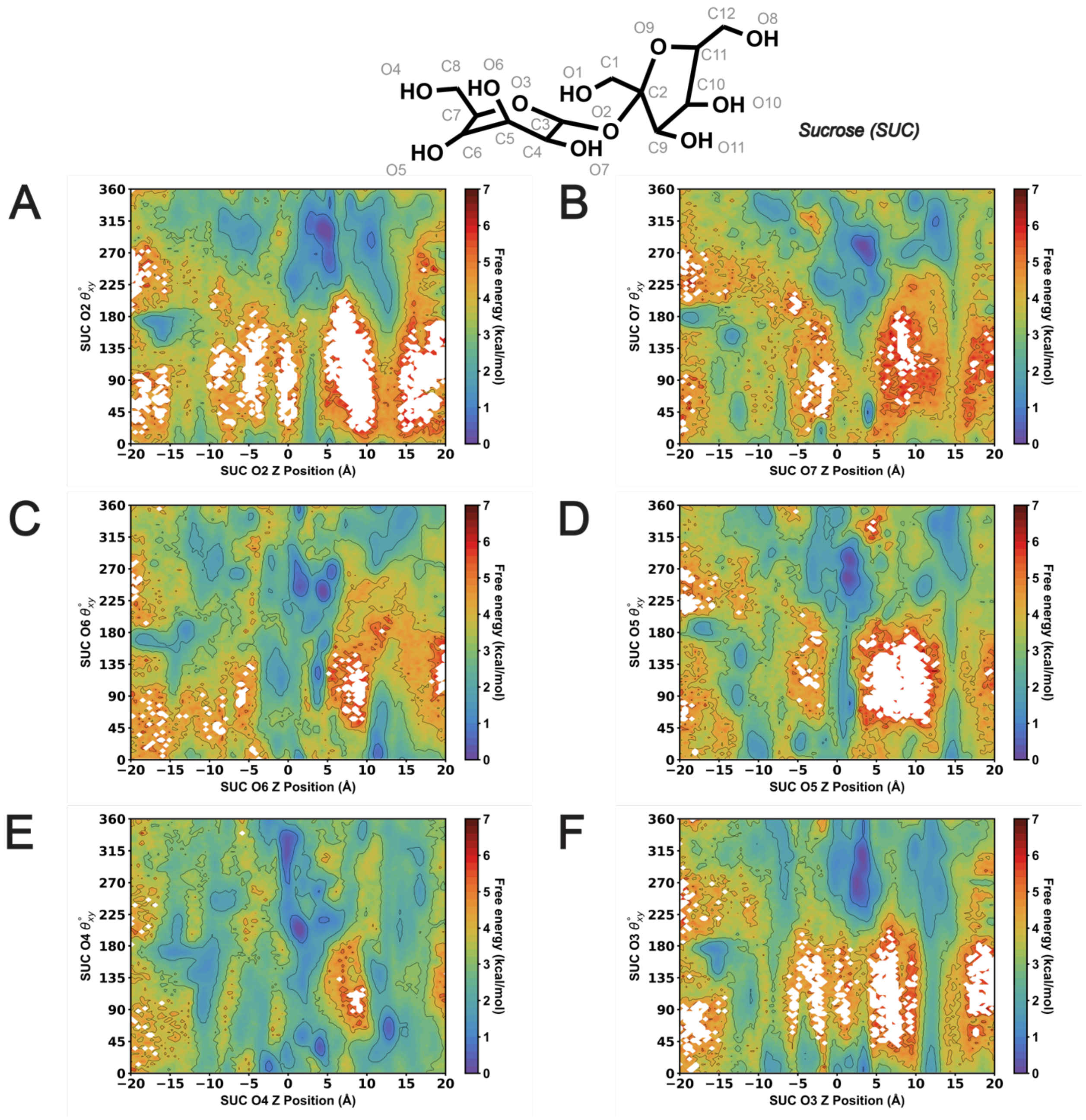


**Figure S21.** MSM-weighted $\theta_{xy}$ analysis versus AtSWEET13 transmembrane channel Z position of the closest SUC molecule glucosyl oxygen atom to the W58-W180 binding pocket. (A) SUC O2. (B) SUC O7 (C) SUC O6. (D) SUC O5. (E) SUC O4. (F) SUC O3.

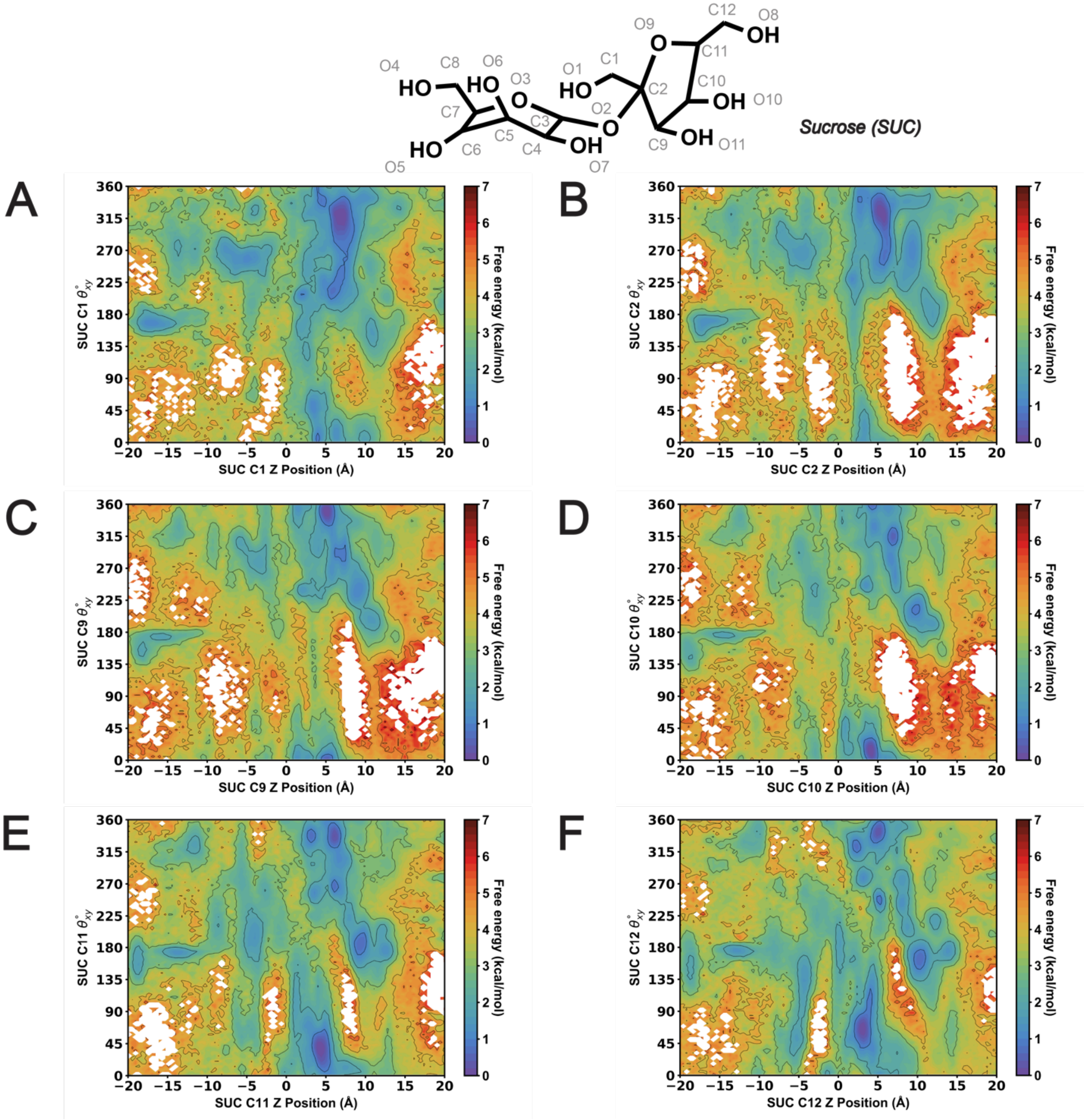


**Figure S22.** MSM-weighted $\theta_{xy}$ analysis versus AtSWEET13 transmembrane channel Z position of the closest SUC molecule fructosyl carbon atom to the W58-W180 binding pocket. (A) SUC C1. (B) SUC C2. (C) SUC C9. (D) SUC C10. (E) SUC C11. (F) SUC C12.

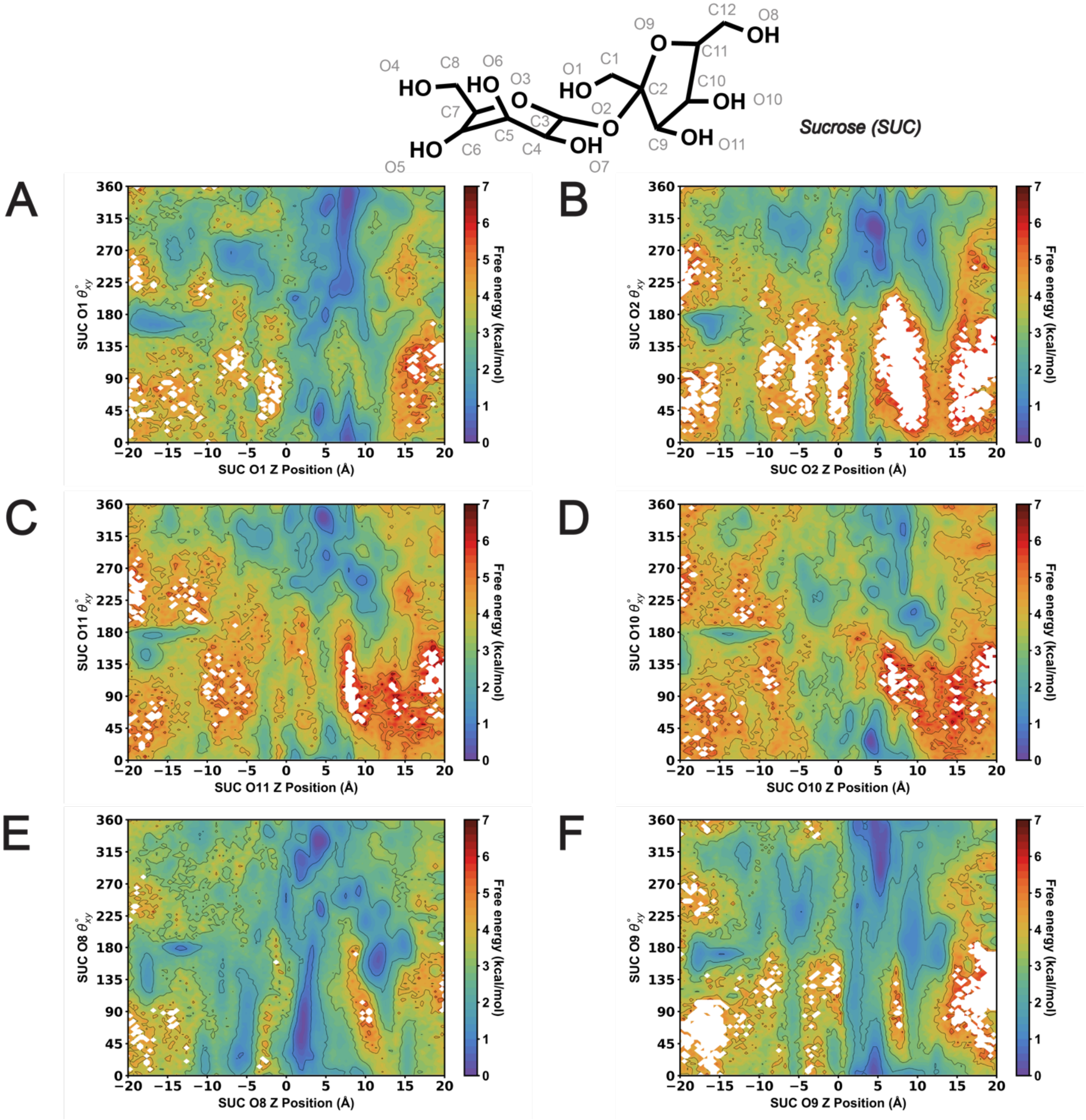


**Figure S23.** MSM-weighted $\theta_{xy}$ analysis versus AtSWEET13 transmembrane channel Z position of the closest SUC molecule fructosyl oxygen atom to the W58-W180 binding pocket. (A) SUC O1. (B) SUC O2 (C) SUC O11. (D) SUC O10. (E) SUC O8. (F) SUC O9.

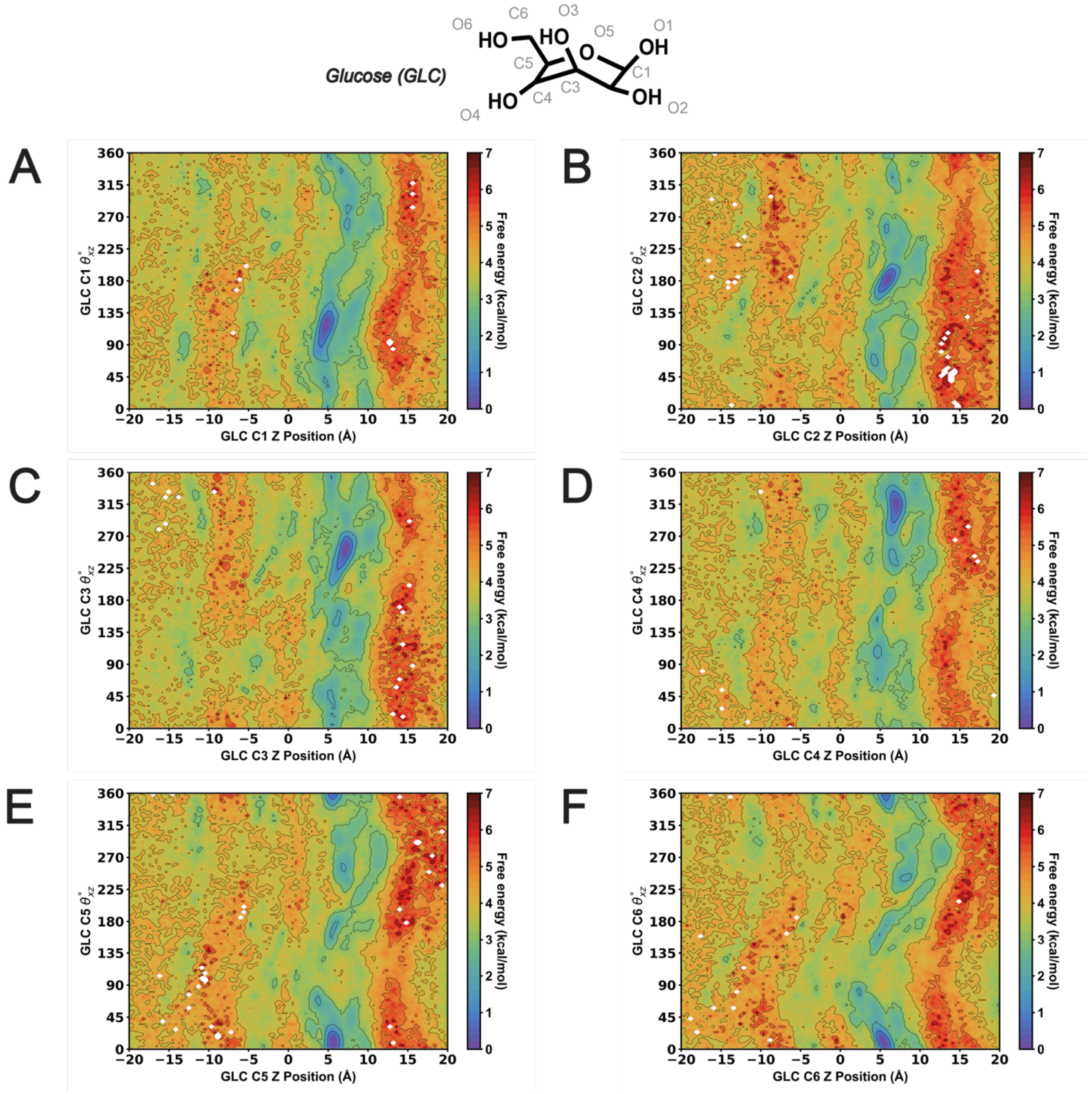


**Figure S24.** MSM-weighted $\theta_{xz}$ analysis versus AtSWEET13 transmembrane channel Z position of the closest GLC molecule carbon atom to the W58-W180 binding pocket. (A) GLC C1. (B) GLC C2 (C) GLC C3. (D) GLC C4. (E) GLC C5. (F) GLC C6.

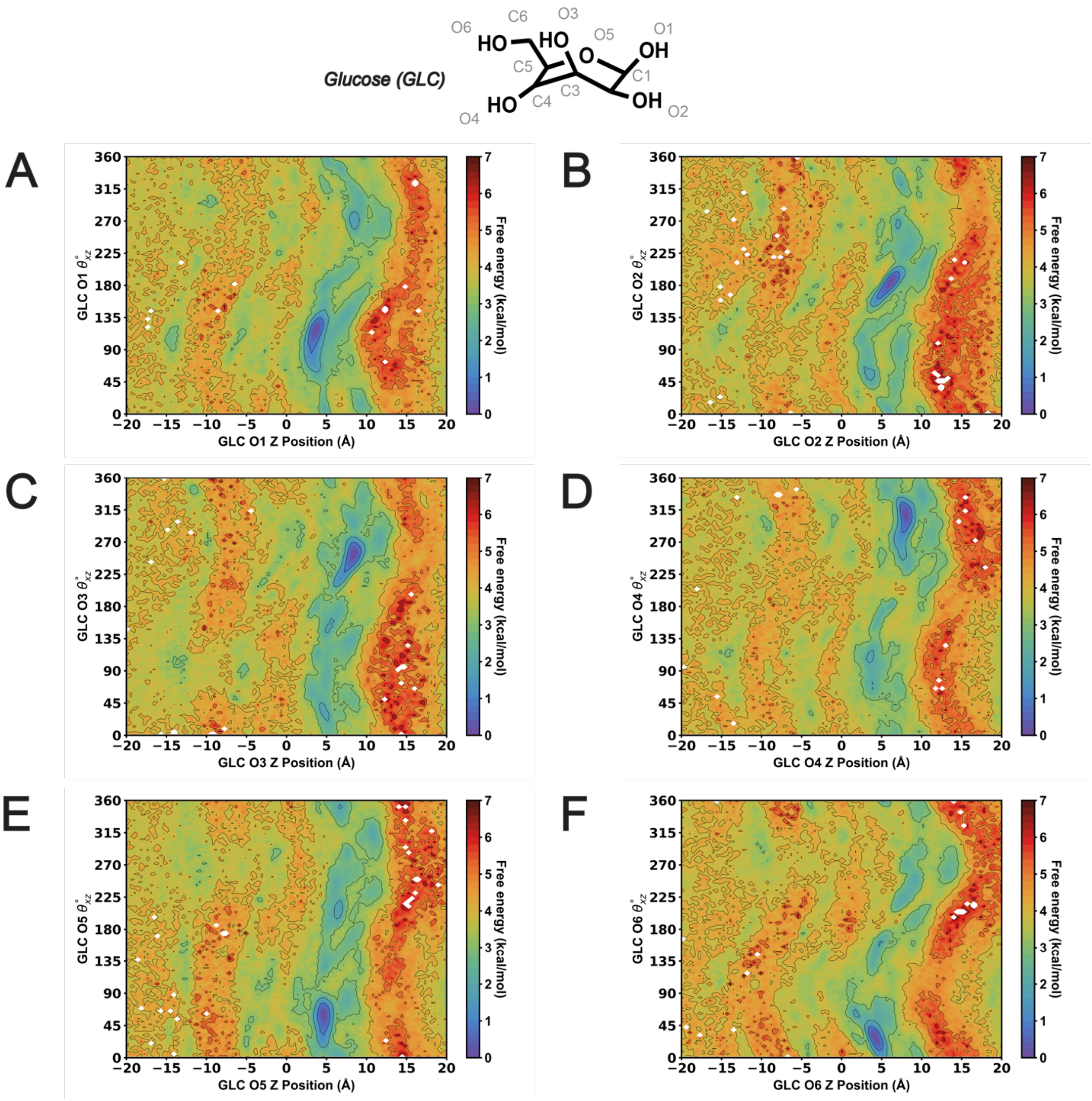


**Figure S25.** MSM-weighted $\theta_{xz}$ analysis versus AtSWEET13 transmembrane channel Z position of the closest GLC molecule oxygen atom to the W58-W180 binding pocket. (A) GLC O1. (B) GLC O2 (C) GLC O3. (D) GLC O4. (E) GLC O5. (F) GLC O6.

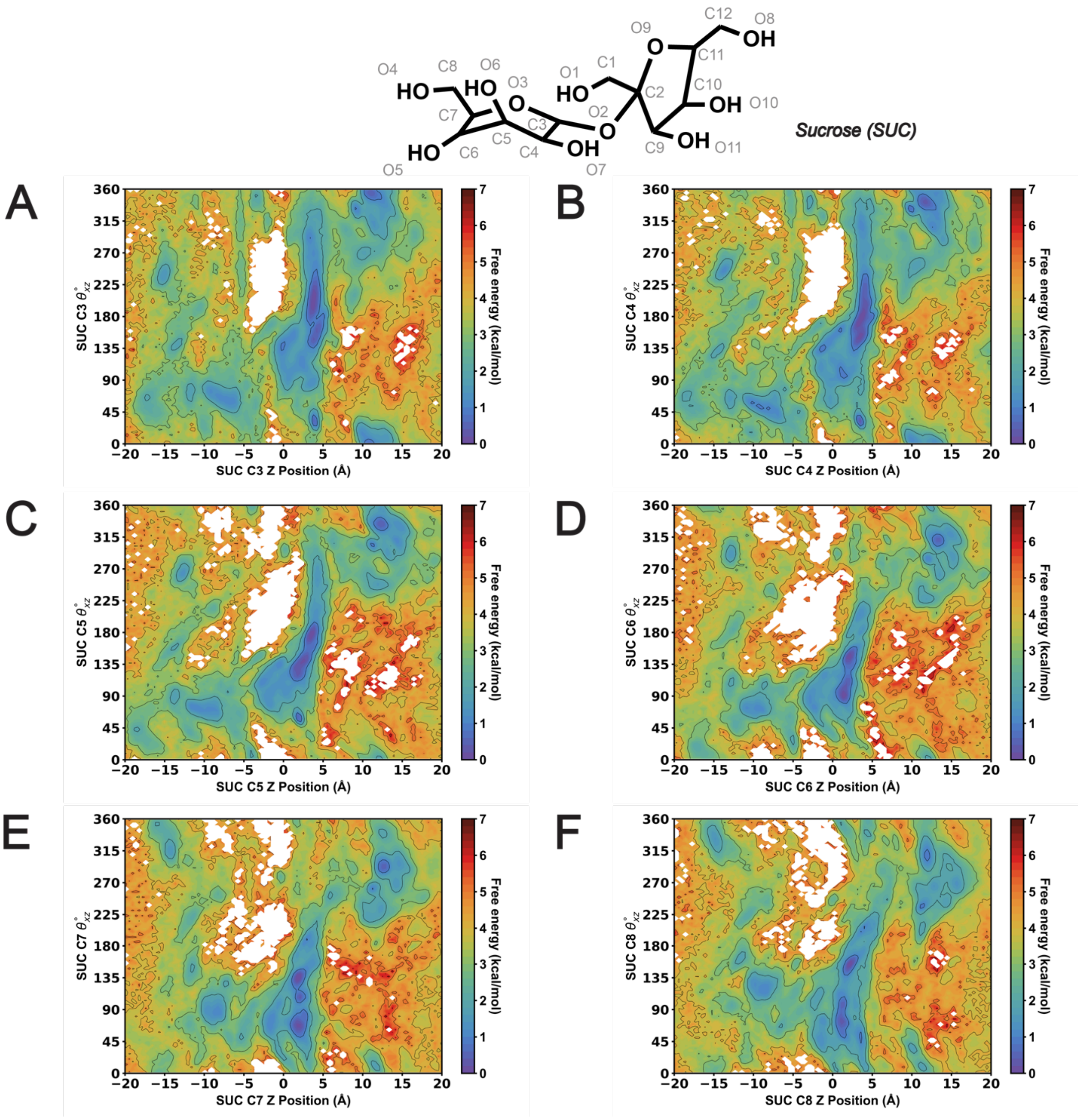


**Figure S26.** MSM-weighted $\theta_{xz}$ analysis versus AtSWEET13 transmembrane channel Z position of the closest SUC molecule glucosyl carbon atom to the W58-W180 binding pocket. (A) SUC C3. (B) SUC C4 (C) SUC C5. (D) SUC C6. (E) SUC C7. (F) SUC C8.

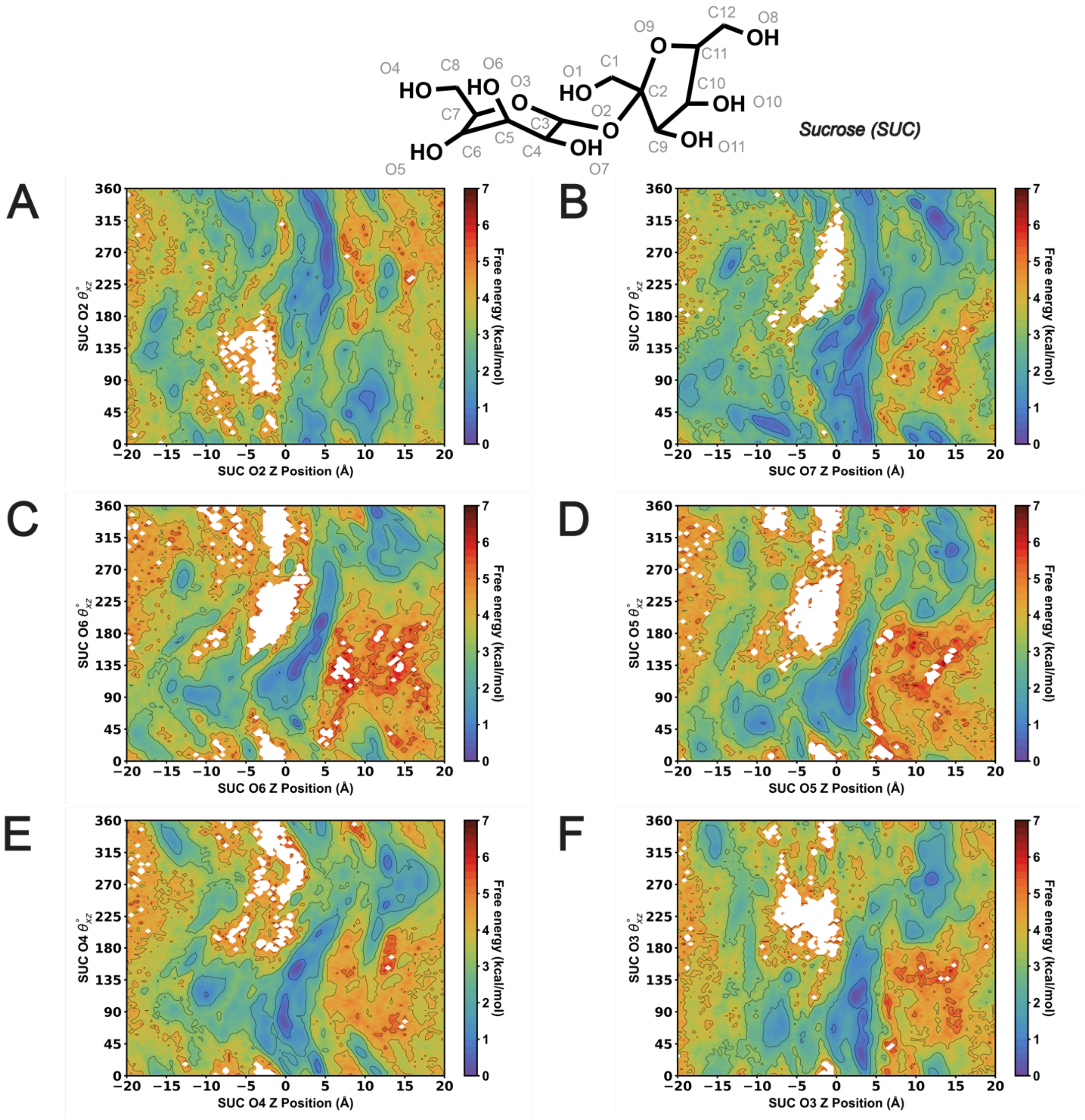


**Figure S27.** MSM-weighted $\theta_{xz}$ analysis versus AtSWEET13 transmembrane channel Z position of the closest SUC molecule glucosyl oxygen atom to the W58-W180 binding pocket. (A) SUC O2. (B) SUC O7 (C) SUC O6. (D) SUC O5. (E) SUC O4. (F) SUC O3.

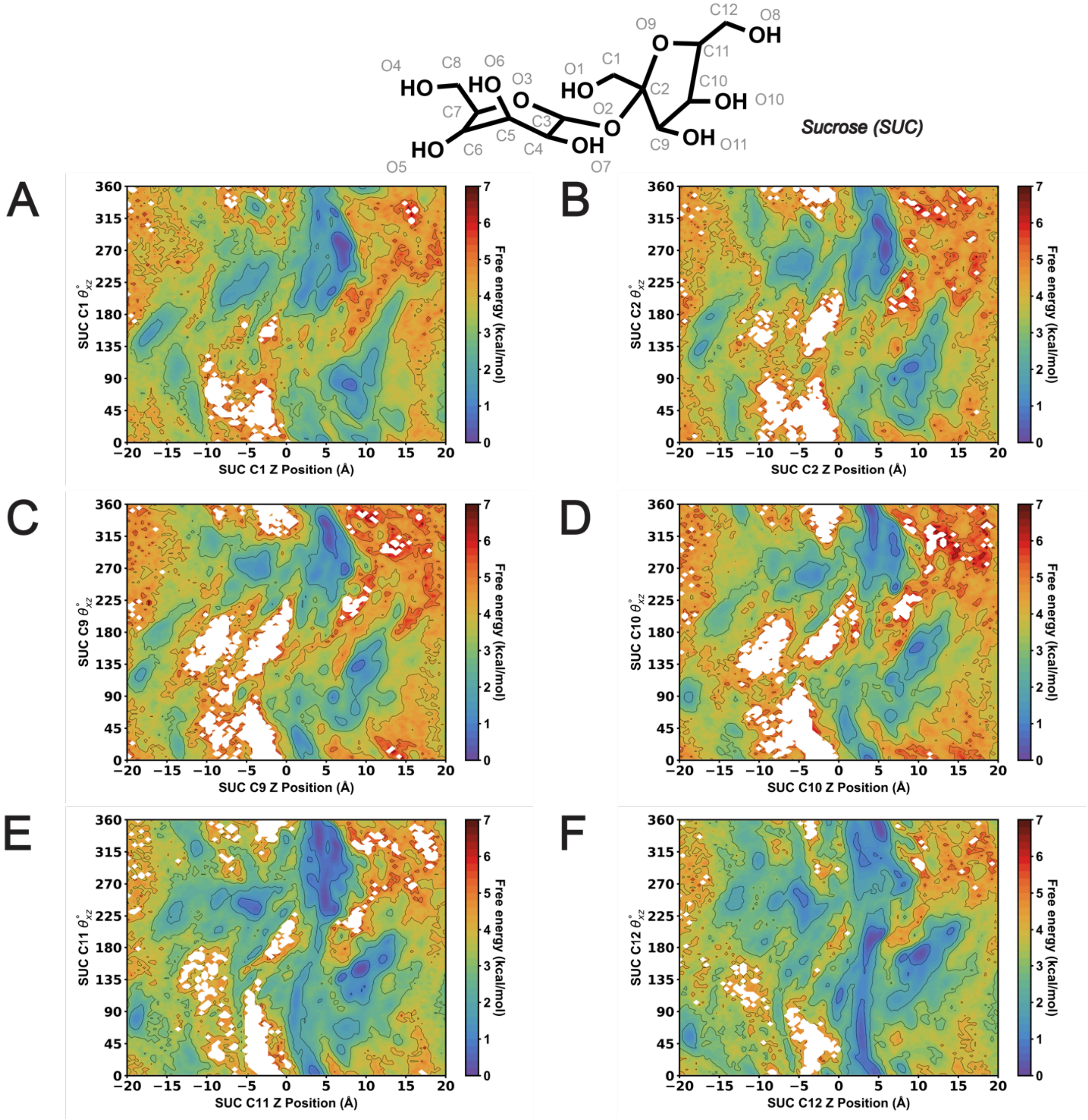


**Figure S28.** MSM-weighted $\theta_{xz}$ analysis versus AtSWEET13 transmembrane channel Z position of the closest SUC molecule fructosyl carbon atom to the W58-W180 binding pocket. (A) SUC C1. (B) SUC C2 (C) SUC C9. (D) SUC C10. (E) SUC C11. (F) SUC C12.

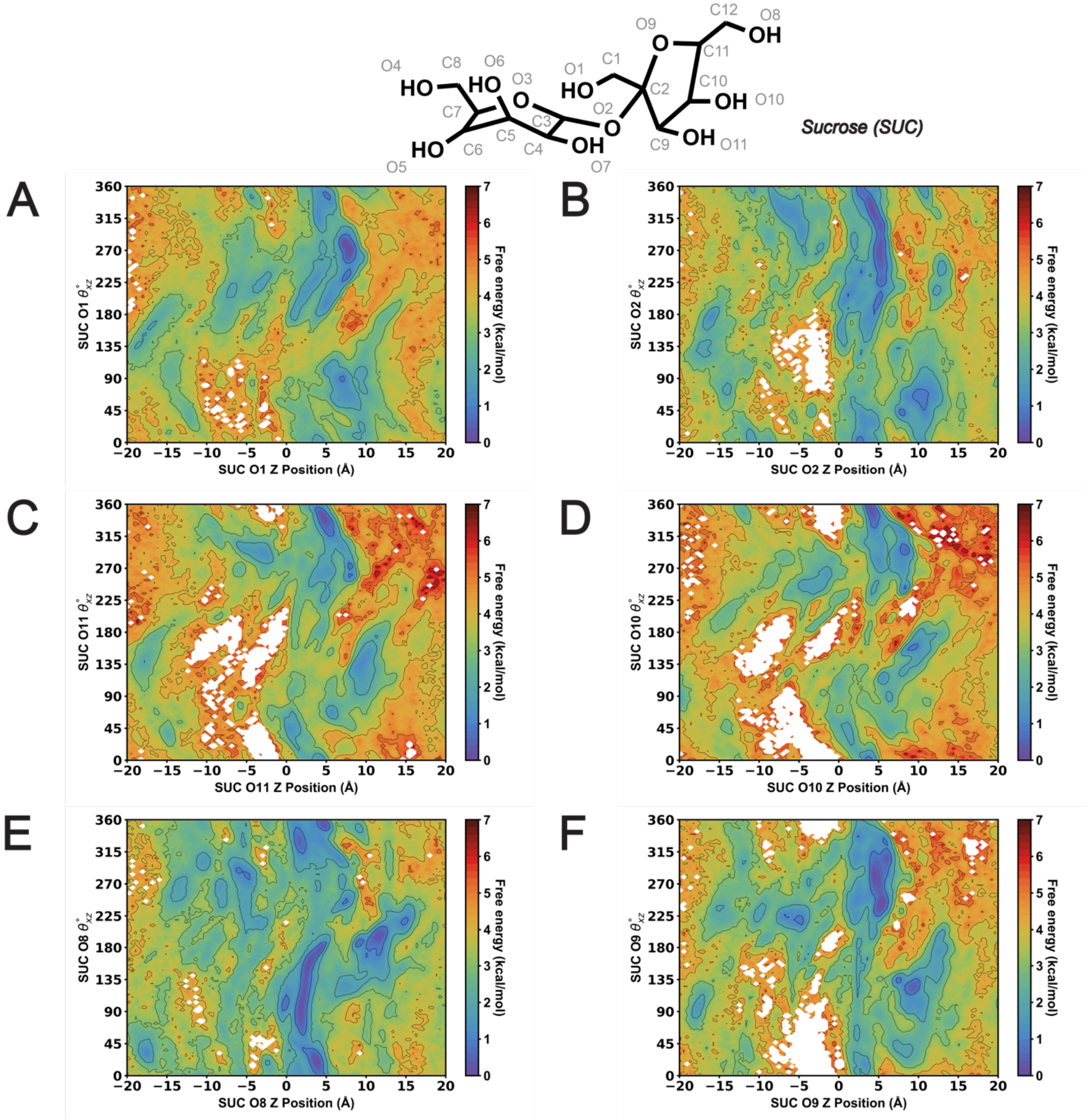


**Figure S29.** MSM-weighted $\theta_{xz}$ analysis versus AtSWEET13 transmembrane channel Z position of the closest SUC molecule fructosyl oxygen atom to the W58-W180 binding pocket. (A) SUC O1. (B) SUC O2 (C) SUC O11. (D) SUC O10. (E) SUC O8. (F) SUC O9.

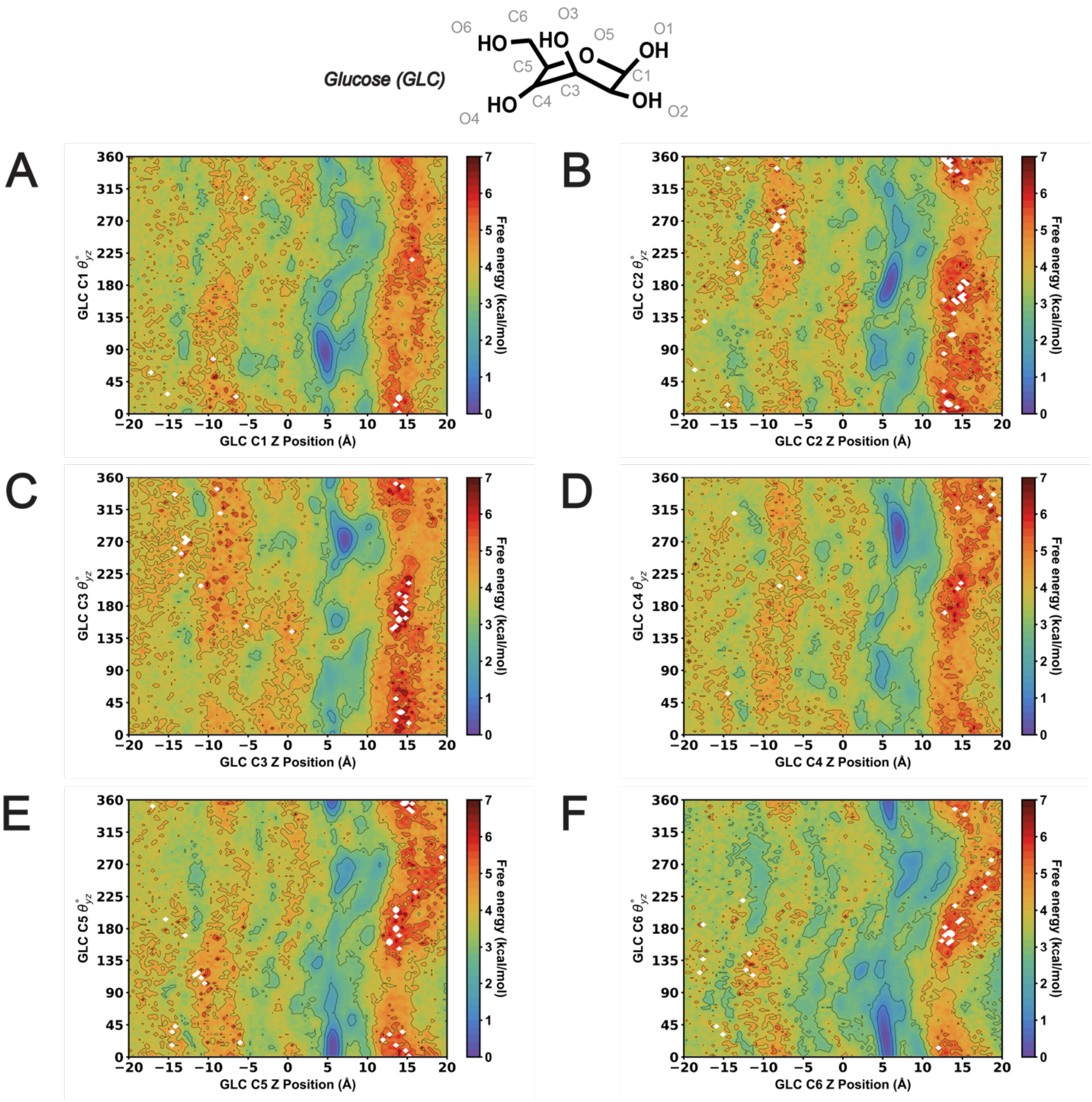


**Figure S30.** MSM-weighted $\theta_{yz}$ analysis versus AtSWEET13 transmembrane channel Z position of the closest GLC molecule carbon atom to the W58-W180 binding pocket. (A) GLC C1. (B) GLC C2 (C) GLC C3. (D) GLC C4. (E) GLC C5. (F) GLC C6.

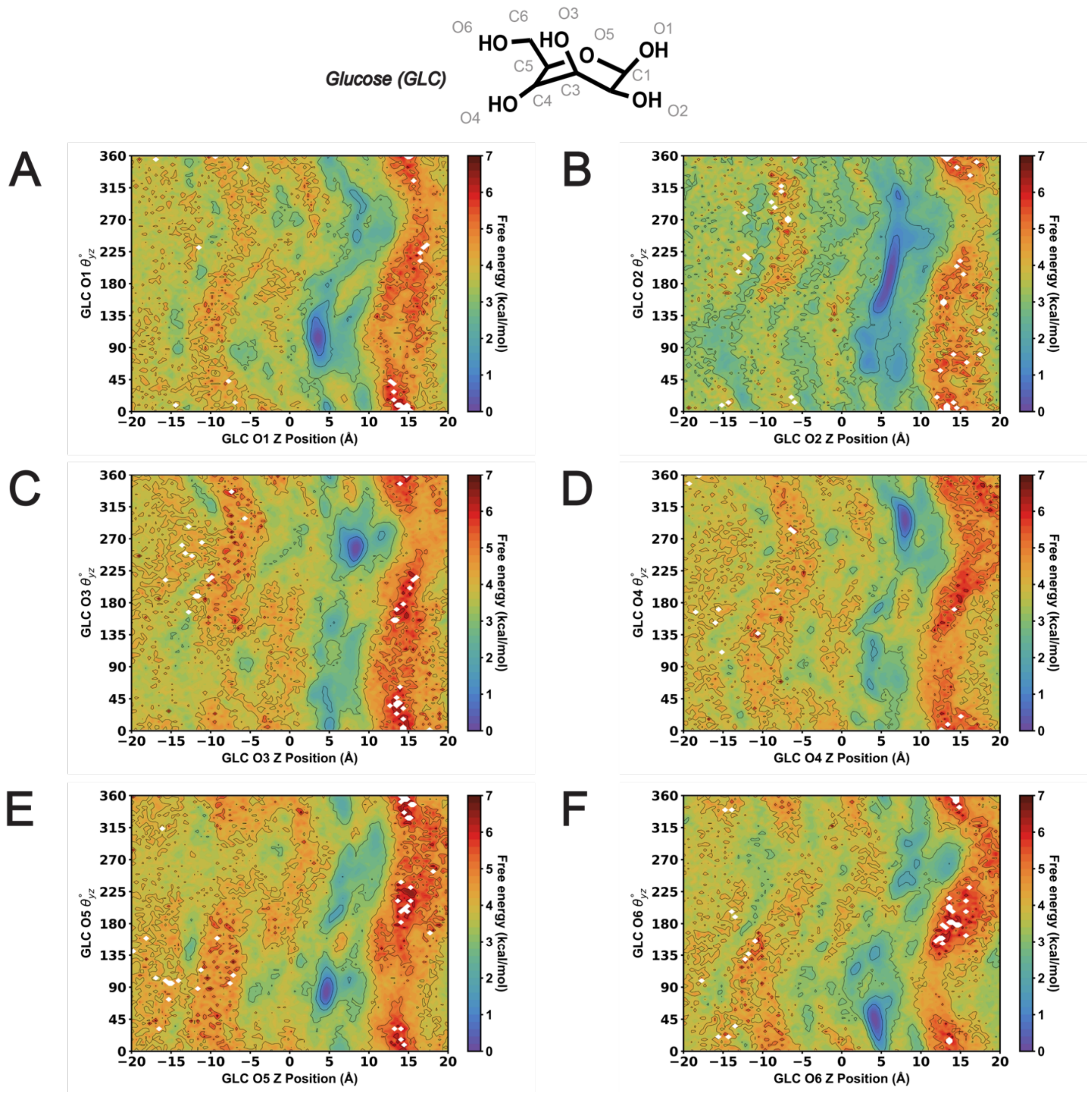


**Figure S31.** MSM-weighted $\theta_{yz}$ analysis versus AtSWEET13 transmembrane channel Z position of the closest GLC molecule oxygen atom to the W58-W180 binding pocket. (A) GLC O1. (B) GLC O2 (C) GLC O3. (D) GLC O4. (E) GLC O5. (F) GLC O6.

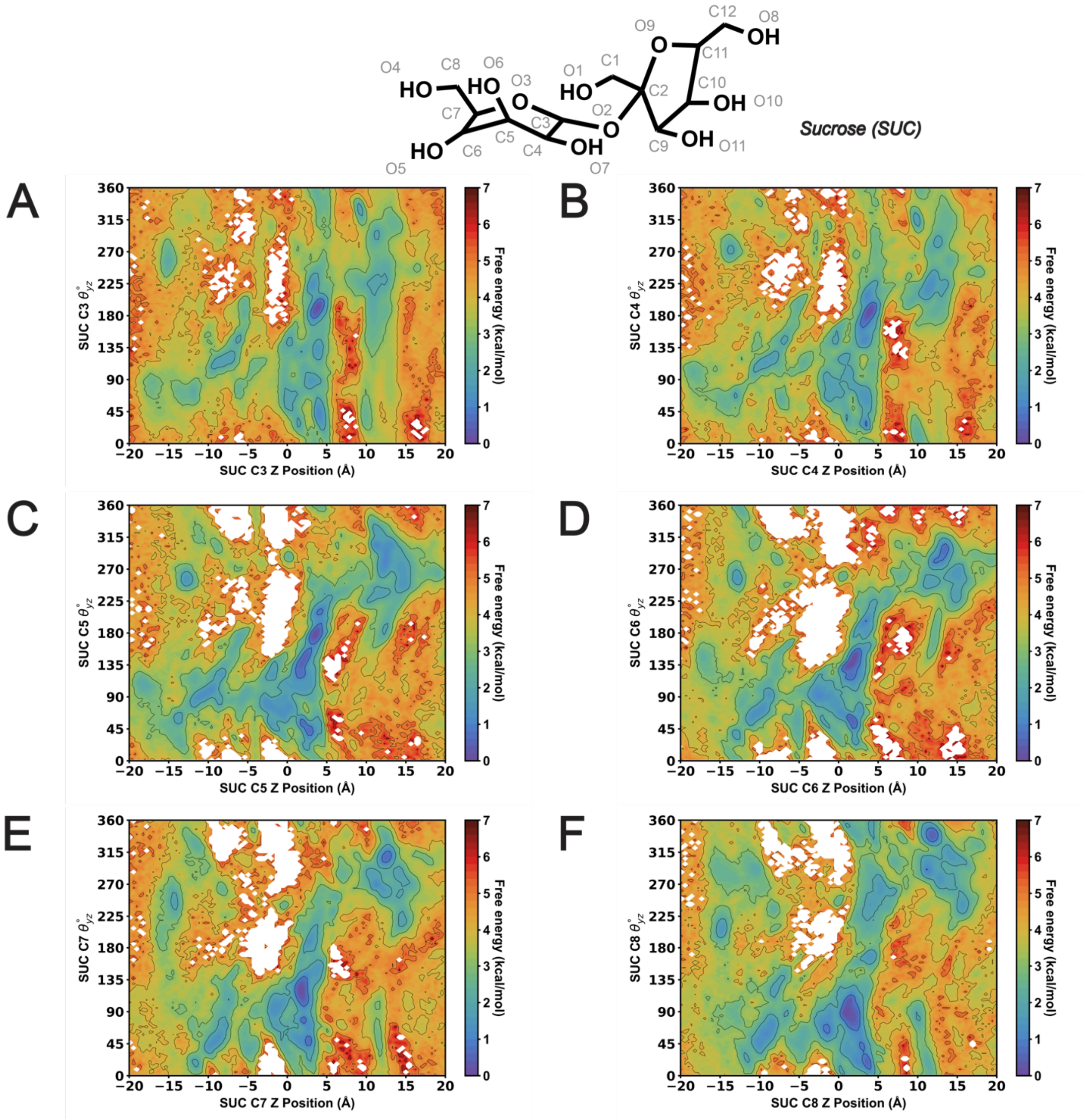


**Figure S32.** MSM-weighted $\theta_{yz}$ analysis versus AtSWEET13 transmembrane channel Z position of the closest SUC molecule glucosyl carbon atom to the W58-W180 binding pocket. (A) SUC C3. (B) SUC C4 (C) SUC C5. (D) SUC C6. (E) SUC C7. (F) SUC C8.

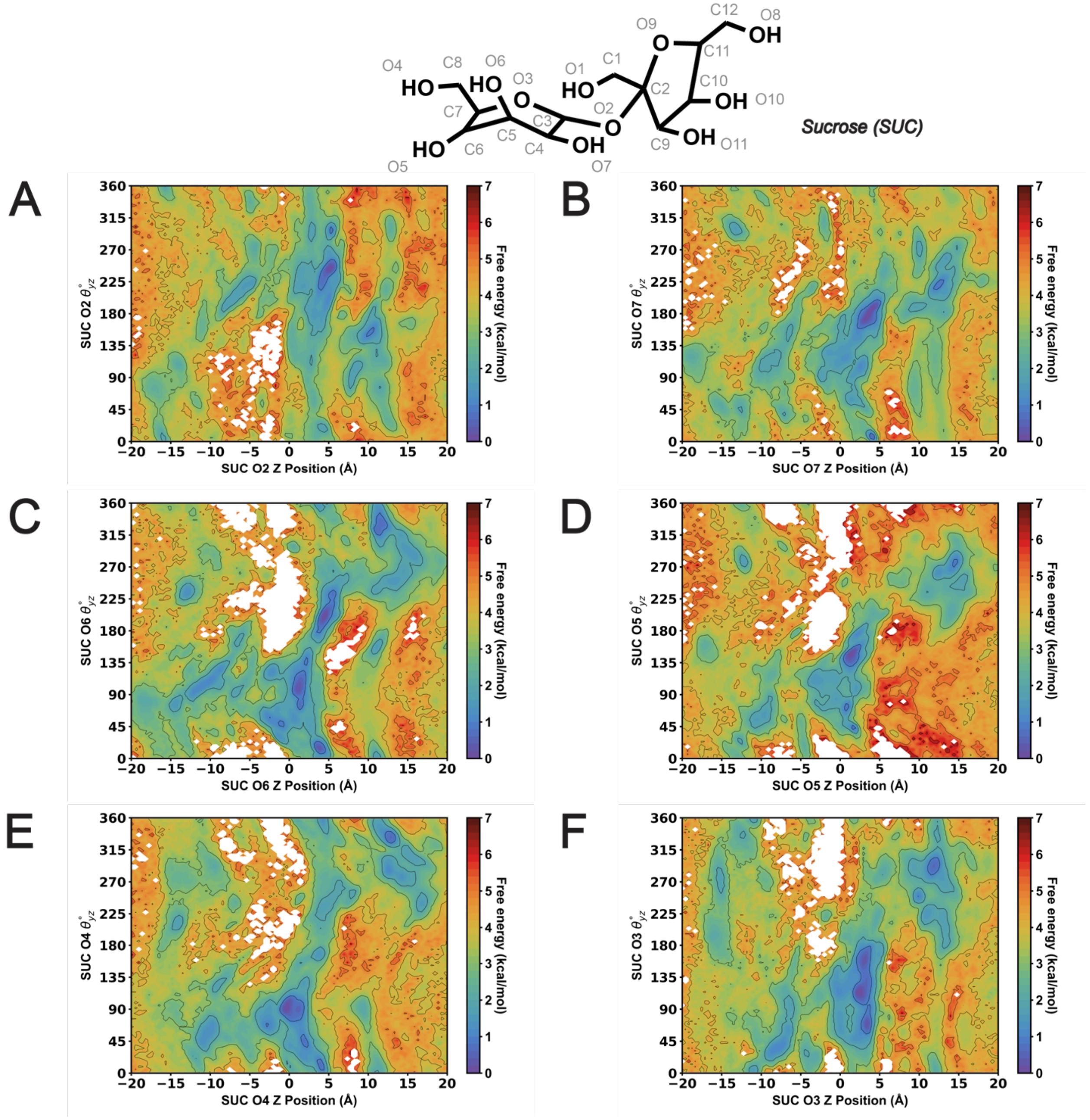


**Figure S33.** MSM-weighted $\theta_{yz}$ analysis versus AtSWEET13 transmembrane channel Z position of the closest SUC molecule glucosyl oxygen atom to the W58-W180 binding pocket. (A) SUC O2. (B) SUC O7 (C) SUC O6. (D) SUC O5. (E) SUC O4. (F) SUC O3.

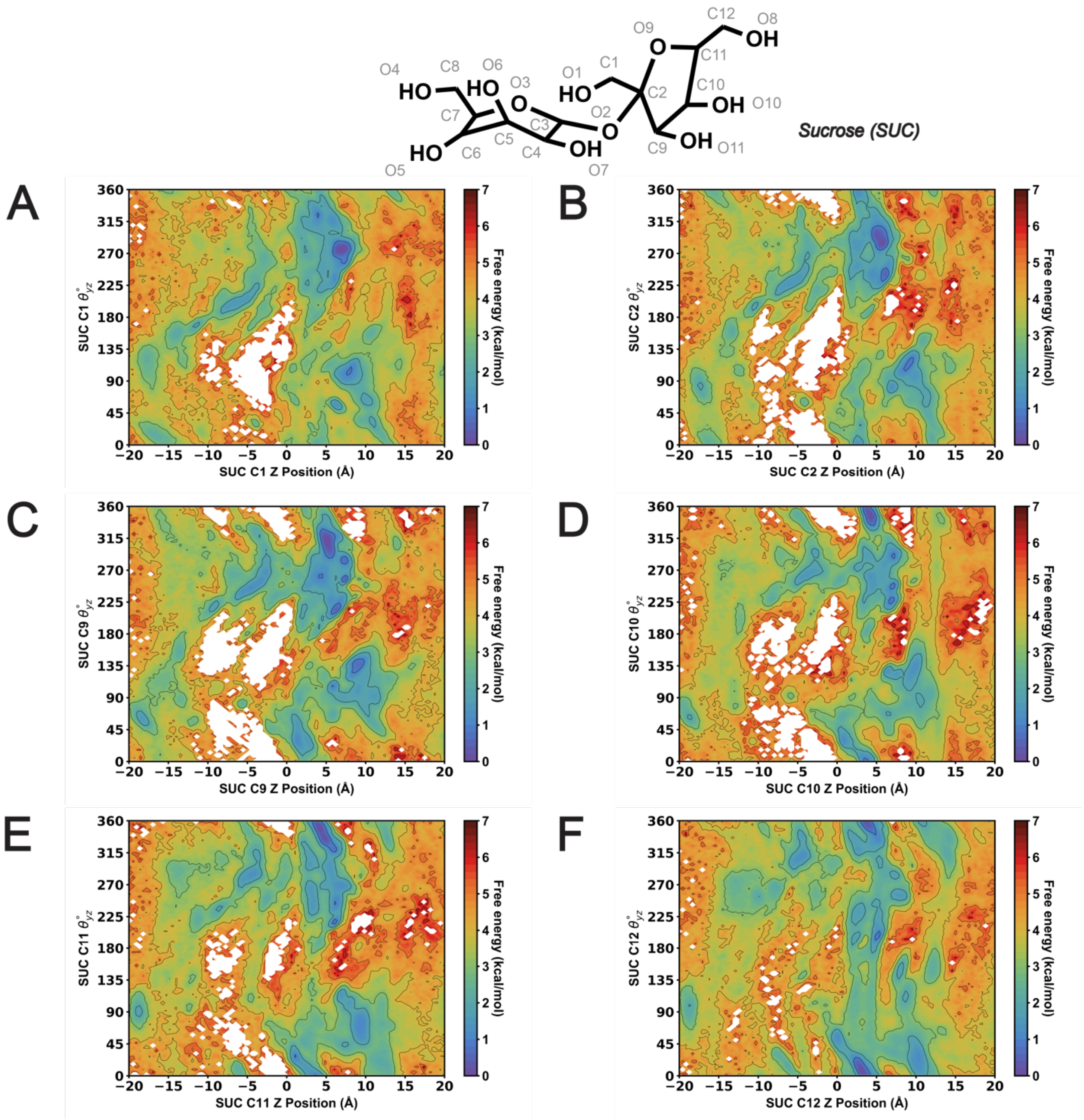


**Figure S34.** MSM-weighted $\theta_{yz}$ analysis versus AtSWEET13 transmembrane channel Z position of the closest SUC molecule fructosyl carbon atom to the W58-W180 binding pocket. (A) SUC C1. (B) SUC C2 (C) SUC C9. (D) SUC C10. (E) SUC C11. (F) SUC C12.

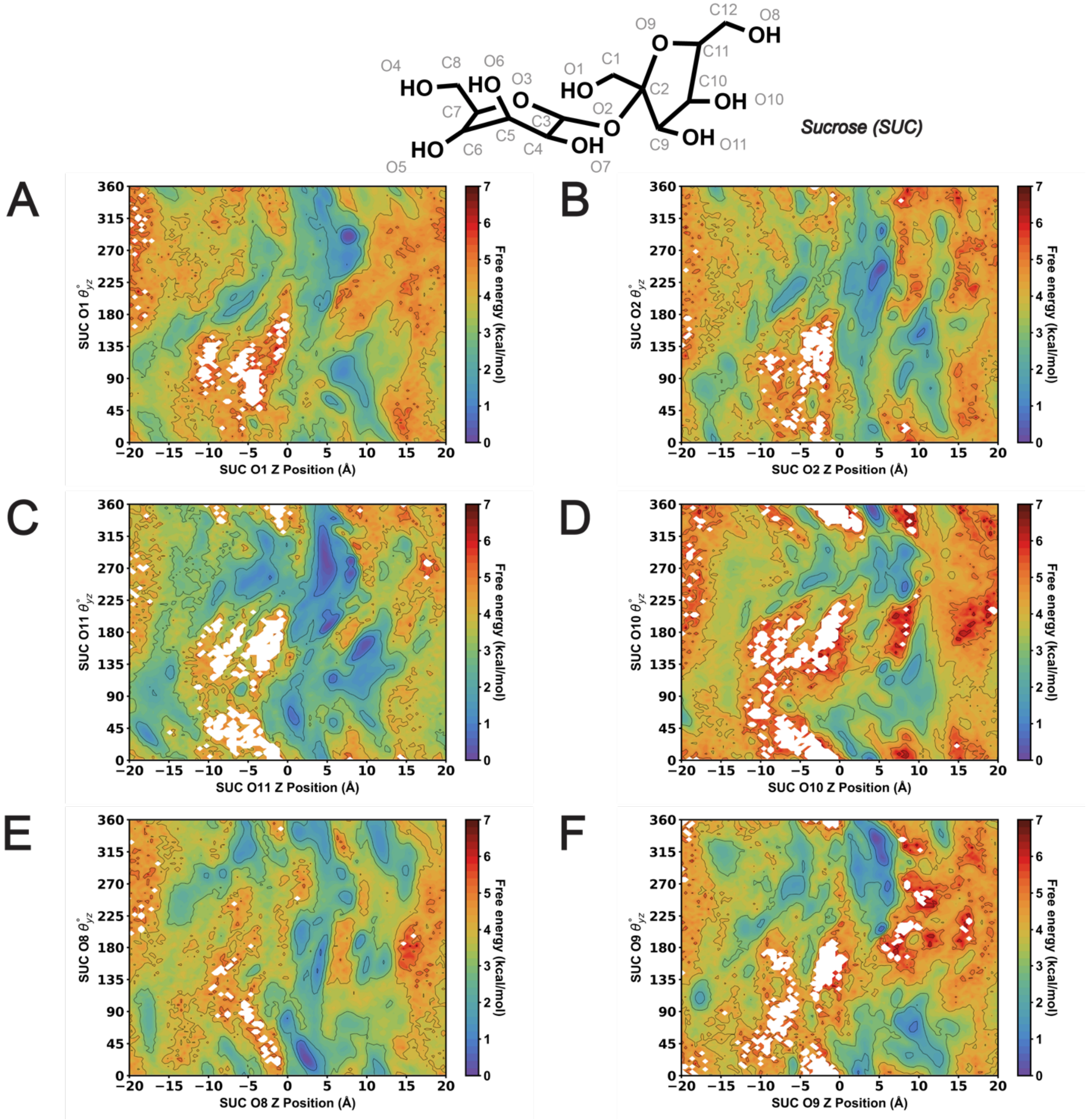


**Figure S35.** MSM-weighted $\theta_{yz}$ analysis versus AtSWEET13 transmembrane channel Z position of the closest SUC molecule fructosyl oxygen atom to the W58-W180 binding pocket. (A) SUC O1. (B) SUC O2 (C) SUC O11. (D) SUC O10. (E) SUC O8. (F) SUC O9.

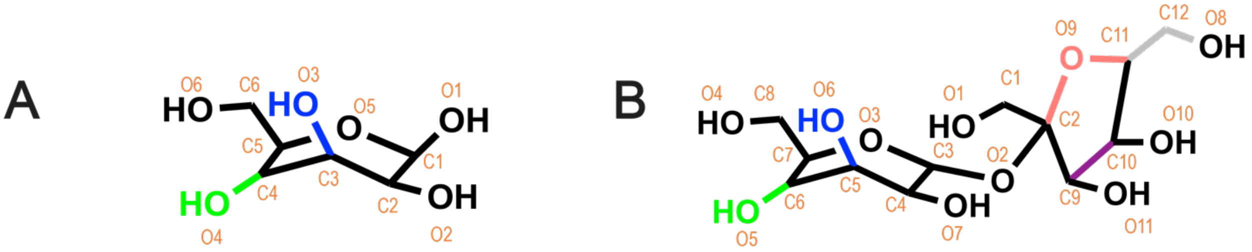


**Figure S36.** GLC and SUC structures with highlighted functional considered critical for molecular recognition by AtSWEET13. Atoms colored blue and green represent moieties which AtSWEET13 recognizes as "GLC-like" between GLC and SUC. Atoms colored salmon, grey, and magenta represent moieties which AtSWEET13 recognizes as "SUC-like". The advent of the fructosyl monomeric unit of SUC is best represented by the salmon, grey, and magenta moieties on SUC. Classification of these functional units was based off rational interpretation of atom-specific $\theta$ rotation analyses.

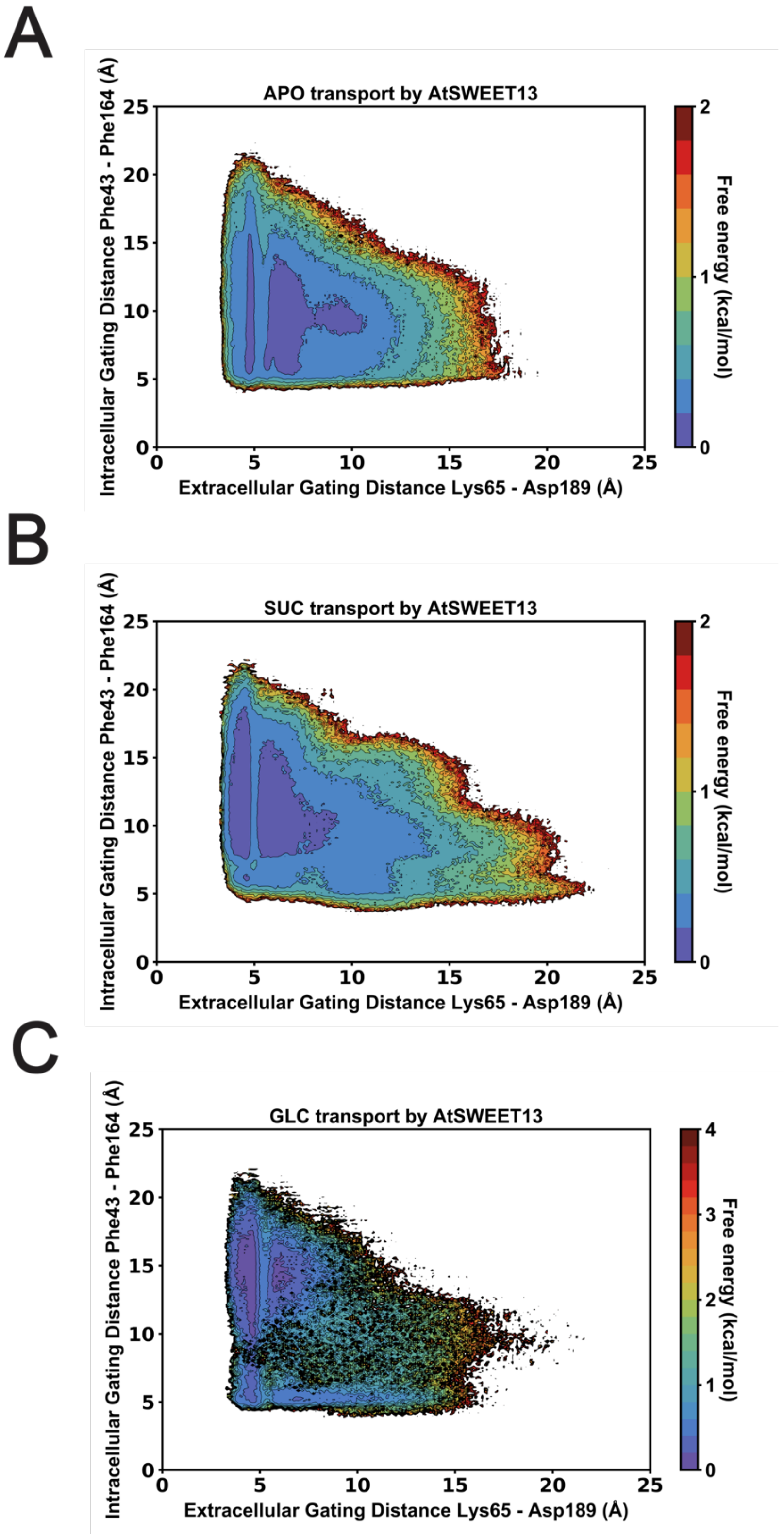


**Figure S37.** MSM-weighted bootstrap error plots for adaptive sampling of (A) *Apo*, (B) SUC, and (C) GLC gating landscapes. Colorbar scales vary between plots to improve landscape resolution and aid in visualization.

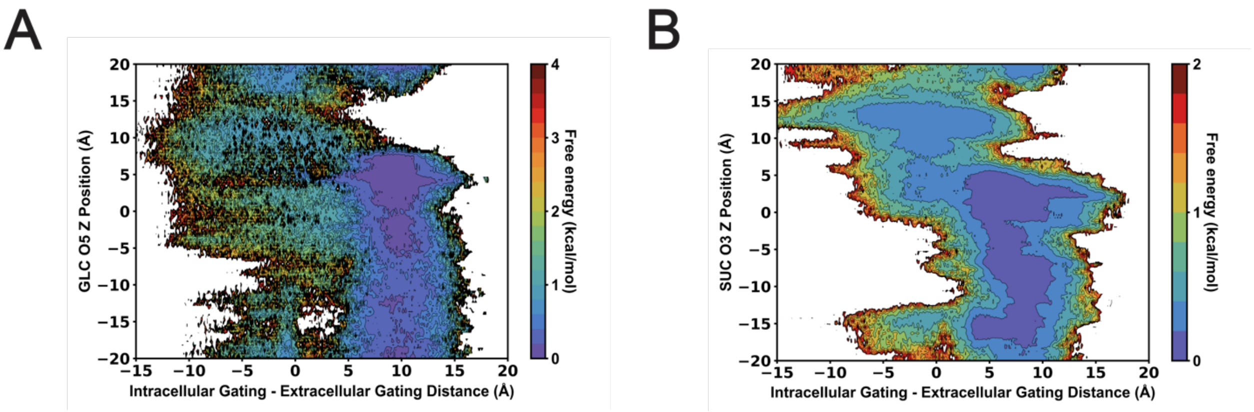


**Figure S38.** MSM-weighted bootstrap error plots for adaptive sampling of (A) SUC and (B) GLC intracellular minus extracellular gating distance versus AtSWEET13 transmembrane channel ligand Z position landscapes. Colorbar scales vary between plots to improve landscape resolution and aid in visualization.

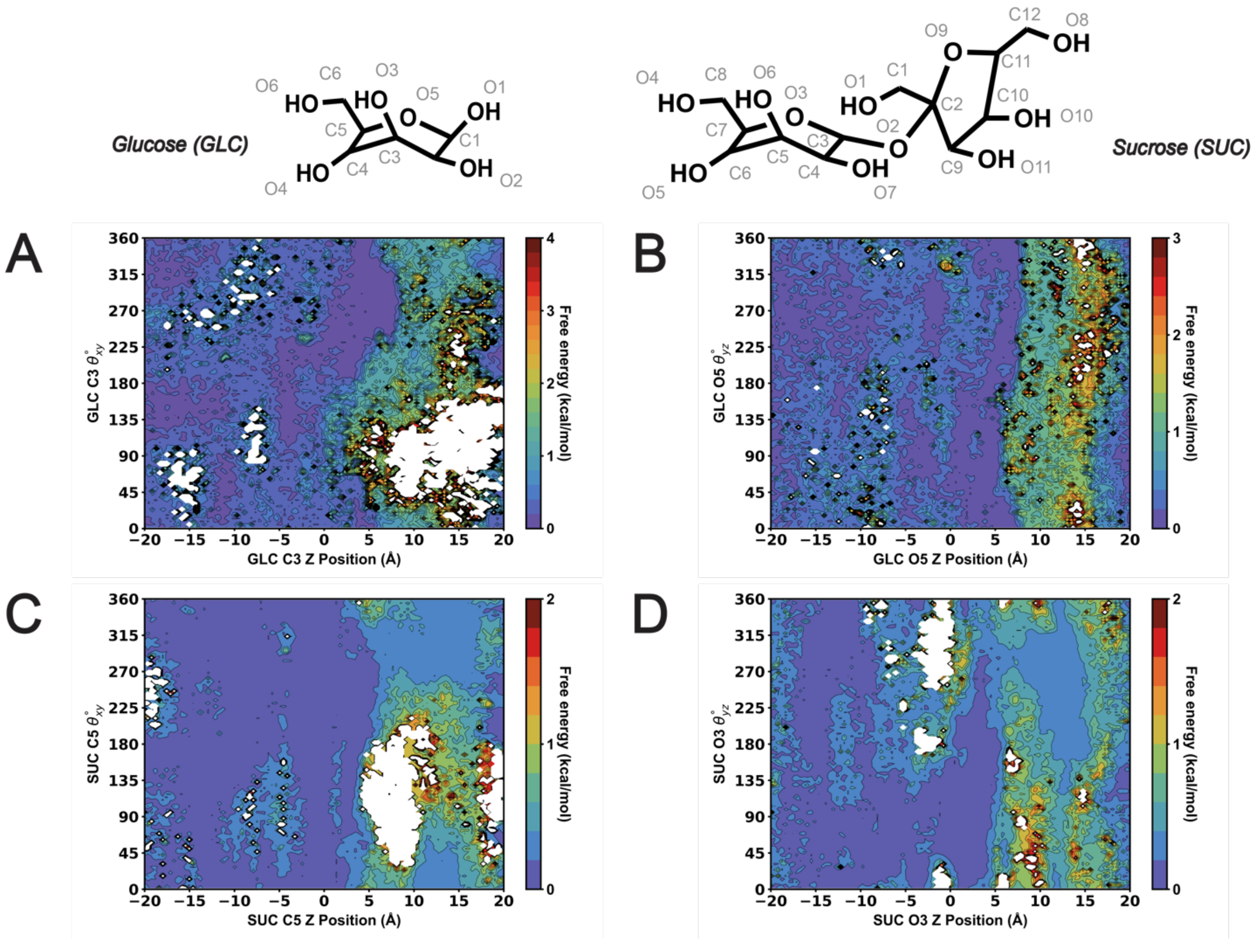


**Figure S39.** MSM-weighted bootstrap error plots for adaptive sampling of $\boldsymbol{\theta}$ rotation analyses presented in Main Text Figure 3. (A) GLC C3 $\boldsymbol{\theta}_{yz}$. (B) GLC O5 $\boldsymbol{\theta}_{yz}$. (C) SUC C5 $\boldsymbol{\theta}_{xy}$. (D) SUC O3 $\boldsymbol{\theta}_{yz}$. Colorbar scales vary between plots to improve landscape resolution and aid in visualization.

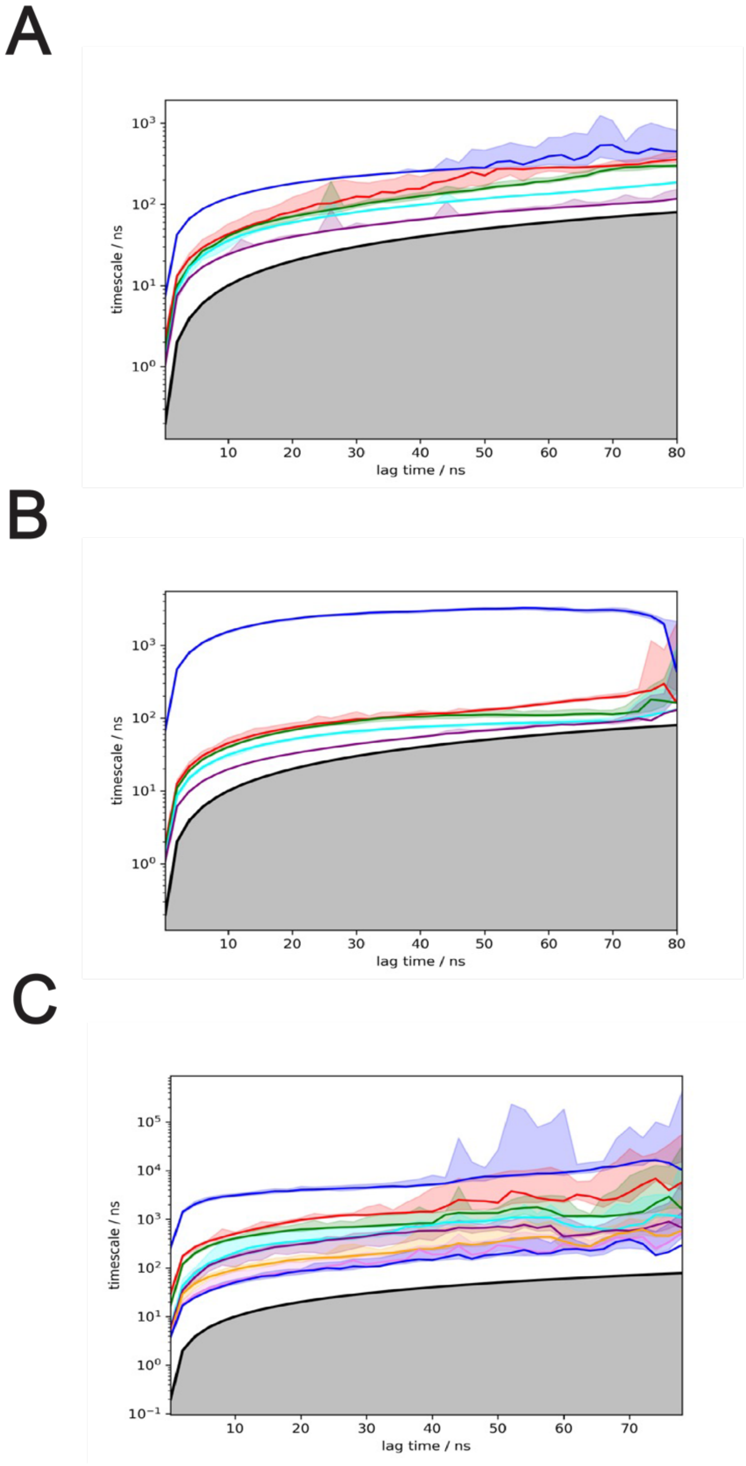


**Figure S40.** Implied timescale plots calculated with Bayesian error for (A) Apo, (B) SUC and (C) GLC transport.

**Table S1.** Finalized features used for MSM discretization. Residue-residue distances are signified as "RES#1_ATOM_RES#2_ATOM". Ligand center of mass (CoM) and atom are listed for either Z position or a specific theta ($\theta$) vector.

| | APO | SUC | GLC |
|---|---|---|---|
| Final MSM features for trajectory discretization | GLY42_CA_GLU163_CB | GLY16_CA_LEU71_CB | GLY13_CA_LEU71 |
| | PRO47_CB_LEU169_CB | GLY13_CA_LEU71_CB | GLY16_CA_LEU71_CB |
| | GLY42_CA_PHE164_CB | ILE75_CB_GLY16_CA | ILE75_CB_GLY16_CA |
| | PHE43_CB_PHE164_CB | LYS65_CB_ASP189_CB | GLY42_CA_THR30_CB |
| | PRO166_CB_PRO47_CB | LYS65_CB_ILE187_CB | GLY42_CA_PRO166_CB |
| | | THR68_CB_ILE187_CB | THR40_CB_PHE43_CB |
| | | SUC CoM Z position | THR68_CB_ILE187_CB |
| | | SUC_C2_theta_xy | PHE43_CB_LEU169_CB |
| | | SUC_C2_theta_xz | GLU41_CB_ARG33_CB |
| | | SUC_C2_theta_yz | ARG33_CB_THR30_CB |
| | | SUC_C5_theta_xy | GLU41_CB_THR30_CB |
| | | SUC_C5_theta_xz | GLU41_CB_ILE34_CB |
| | | SUC_C5_theta_yz | GLY42_CA_MET165_CB |
| | | SUC_C6_theta_xy | GLC CoM Z position |
| | | SUC_C6_theta_xz | |
| | | SUC_C6_theta_yz | |
| | | SUC_C9_theta_xy | |
| | | SUC_C9_theta_xz | |
| | | SUC_C9_theta_yz | |
| | | SUC_C10_theta_xy | |
| | | SUC_C10_theta_xz | |
| | | SUC_C10_theta_yz | |
| | | SUC_C11_theta_xy | |
| | | SUC_C11_theta_xz | |
| | | SUC_C11_theta_yz | |
| | | SUC_C12_theta_xy | |
| | | SUC_C12_theta_xz | |
| | | SUC_C12_theta_yz | |
| | | SUC_O5_theta_xy | |
| | | SUC_O5_theta_xz | |
| | | SUC_O5_theta_yz | |
| | | SUC_O6_theta_xy | |
| | | SUC_O6_theta_xz | |
| | | SUC_O6_theta_yz | |
| | | SUC_O9_theta_xy | |
| | | SUC_O9_theta_xz | |
| | | SUC_O9_theta_yz | |

**Table S2.** Finalized MSM hyperparameters

| System | # Clusters | tICA dimensions | tiCA lag time | MSM lag time |
|---|---|---|---|---|
| **APO** | 950 | 3 | 8 ns | 30 ns |
| **SUC** | 900 | 8 | 8 ns | 40 ns |
| **GLC** | 750 | 12 | 8 ns | 20 ns |

The following pages include Supplementary Tables containing each of the features used for the feature-diverse tICA decompositions for Apo, SUC, and GLC transport by AtSWEET13. Residue-residue distances are signified as “RES#1_ATOM_RES#2_ATOM”, where beta carbons were used for all residues with the exception of alpha carbon for glycines. The Z position of the ligand is approximated using the distance between an AtSWEET13 residue versus a specific ligand atom provided the atomic substrate numbering shown in Supplemental Figure XX. Protein residue-ligand atom distances are signified as “RES##_RES-ATOM_LIGAND-ATOM”, where the “LIGAND-ATOM”. Specific ligand atom theta ($\theta$) rotational vectors are enumerated as “theta_LIGAND-ATOM_VECTOR”.

**Table S3.** Feature-diverse tICA decomposition features and correlations – APO transport

| System | APO | |
|---|---|---|
| # of features | 113 | |
| Feature | \| tIC1 correlation (*r*) \| | \| tIC2 correlation (*r*) \| |
| GLY203_CA_ALA177_CB | 0.064460833 | 0.150293639 |
| LEU173_CB_ALA177_CB | 0.005209448 | 0.001928949 |
| SER176_CB_ALA177_CB | 0.000325243 | 0.002499811 |
| MET1_CB_ALA2_CB | 0.026115698 | 0.000597437 |
| ALA177_CB_ALA200_CB | 0.066300157 | 0.051185722 |
| GLY199_CA_ALA200_CB | 0.008626179 | 0.003866548 |
| GLY203_CA_ALA200_CB | 0.039433079 | 0.109160419 |
| PRO47_CB_ALA51_CB | 0.001360774 | 0.006165371 |
| SER50_CB_ALA51_CB | 0.085784049 | 0.058114437 |
| TYR48_CB_ALA51_CB | 0.018641856 | 0.011838054 |
| GLY79_CA_ALA55_CB | 0.064283787 | 0.020557796 |
| SER54_CB_ALA55_CB | 0.021222477 | 0.002871329 |
| LEU72_CB_ALA62_CB | 0.250428519 | 0.048609968 |
| TRP58_CB_ALA62_CB | 0.055796707 | 0.036414729 |
| TYR61_CB_ALA62_CB | 0.057345646 | 0.00718485 |
| ILE75_CB_ASN17_CB | 0.045712681 | 0.036624931 |
| SER20_CB_ASN17_CB | 0.179467098 | 0.114037893 |
| ALA177_CB_ASN196_CB | 0.035604665 | 0.004052984 |
| ALA200_CB_ASN196_CB | 0.049786031 | 0.022545655 |
| GLY199_CA_ASN196_CB | 0.082495759 | 0.186440036 |
| TRP180_CB_ASN196_CB | 0.085721694 | 0.117302715 |
| ALA55_CB_ASN76_CB | 0.001282193 | 0.000157732 |
| GLY79_CA_ASN76_CB | 0.063124379 | 0.014353045 |
| GLY184_CA_ASP189_CB | 0.112907427 | 0.010375813 |
| ILE187_CB_ASP189_CB | 0.01726444 | 0.12000357 |
| SER142_CB_CYS138_CB | 0.067249785 | 0.170817157 |
| TRP58_CB_CYS138_CB | 0.020713462 | 0.066127972 |
| GLY135_CA_GLU131_CB | 0.081217656 | 0.05563119 |
| GLY42_CA_GLU163_CB | 0.735515645 | 0.067425631 |
| SER161_CB_GLU163_CB | 0.053415735 | 0.33693962 |
| SER39_CB_GLU41_CB | 0.012032227 | 0.006235093 |

**Table S3.** Feature-diverse tICA features and correlations – APO transport (cont.)

| System | APO | |
|---|---|---|
| # of features | 113 | |
| **Feature** | **\| tIC1 correlation (*r*) \|** | **\| tIC2 correlation (*r*) \|** |
| **ALA51_CB_GLU83_CB** | 0.005101782 | 0.032917297 |
| **TYR48_CB_GLU83_CB** | 0.022201456 | 0.001956743 |
| **TYR86_CB_GLU83_CB** | 0.021986978 | 0.033722716 |
| **GLY16_CA_GLY13_CA** | 0.060467994 | 0.049752374 |
| **CYS138_CB_GLY135_CA** | 0.026006159 | 0.077873316 |
| **TRP180_CB_GLY184_CA** | 0.034540545 | 0.01706313 |
| **TYR183_CB_GLY184_CA** | 0.006899806 | 0.00031804 |
| **ALA177_CB_GLY199_CA** | 0.067492236 | 0.117613163 |
| **PHE146_CB_GLY199_CA** | 0.454933861 | 0.401047147 |
| **LEU173_CB_GLY203_CA** | 0.285508713 | 0.045457826 |
| **GLU41_CB_GLY42_CA** | 0.277036454 | 0.248234924 |
| **PHE24_CB_GLY79_CA** | 0.358621257 | 0.179616333 |
| **SER151_CB_ILE152_CB** | 0.12620958 | 0.124922071 |
| **VAL155_CB_ILE152_CB** | 0.18085867 | 0.038369749 |
| **LEU3_CB_ILE187_CB** | 0.160003438 | 0.313185921 |
| **THR4_CB_ILE187_CB** | 0.22292298 | 0.19711629 |
| **THR68_CB_ILE187_CB** | 0.44787437 | 0.201997805 |
| **ARG33_CB_ILE34_CB** | 0.092810043 | 0.143257653 |
| **SER20_CB_ILE75_CB** | 0.14280683 | 0.001793725 |
| **PRO149_CB_LEU169_CB** | 0.35857467 | 0.359675308 |
| **PRO166_CB_LEU169_CB** | 0.078799411 | 0.101652178 |
| **PRO47_CB_LEU169_CB** | 0.821513287 | 0.148678108 |
| **LEU169_CB_LEU173_CB** | 0.0125623 | 0.039361481 |
| **PHE146_CB_LEU173_CB** | 0.465176549 | 0.186405301 |
| **PHE213_CB_LEU209_CB** | 0.356936091 | 0.47516041 |
| **ASN17_CB_LEU71_CB** | 0.326937327 | 0.196381578 |
| **GLY13_CA_LEU71_CB** | 0.627582054 | 0.322271582 |
| **ILE75_CB_LEU71_CB** | 0.083740927 | 0.020558263 |
| **TRP58_CB_LEU72_CB** | 0.069774474 | 0.046436075 |
| **GLY135_CA_LYS132_CB** | 0.074077945 | 0.041010322 |
| **ARG33_CB_LYS37_CB** | 0.072699973 | 0.109079388 |

**Table S3.** Feature-diverse tICA features and correlations – APO transport (cont.)

| System | APO | |
|---|---|---|
| # of features | 113 | |
| **Feature** | **\| tIC1 correlation (*r*) \|** | **\| tIC2 correlation (*r*) \|** |
| SER39_CB_LYS37_CB | 0.007851825 | 0.007064312 |
| MET1_CB_LYS65_CB | 0.242637629 | 0.257193026 |
| PHE164_CB_MET165_CB | 0.234751767 | 0.104128171 |
| VAL162_CB_MET165_CB | 0.054154012 | 0.503362302 |
| GLU163_CB_PHE164_CB | 0.168823715 | 0.005187457 |
| GLY42_CA_PHE164_CB | 0.801658424 | 0.243121215 |
| PHE43_CB_PHE164_CB | 0.844202892 | 0.188729683 |
| SER161_CB_PHE164_CB | 0.193549353 | 0.310257316 |
| VAL162_CB_PHE213_CB | 0.415664029 | 0.486682517 |
| THR40_CB_PHE43_CB | 0.101580028 | 0.162793055 |
| ILE152_CB_PRO149_CB | 0.014511258 | 0.018231714 |
| PRO166_CB_PRO149_CB | 0.258017293 | 0.408594695 |
| PRO47_CB_PRO149_CB | 0.544849013 | 0.135382046 |
| SER142_CB_PRO195_CB | 0.015222509 | 0.149996912 |
| VAL139_CB_PRO195_CB | 0.271191059 | 0.23488928 |
| THR30_CB_PRO27_CB | 0.148546153 | 0.091809021 |
| PRO166_CB_PRO47_CB | 0.738749881 | 0.003660056 |
| VAL145_CB_SER142_CB | 0.0287935 | 0.001135269 |
| LEU173_CB_SER176_CB | 0.134607704 | 0.010397381 |
| VAL23_CB_SER176_CB | 0.084040537 | 0.058601657 |
| VAL23_CB_SER20_CB | 0.095537355 | 0.108487306 |
| ALA51_CB_SER54_CB | 0.003426548 | 0.038068069 |
| SER50_CB_SER54_CB | 0.013867151 | 0.028632169 |
| VAL145_CB_SER54_CB | 0.25547349 | 0.267772313 |
| SER161_CB_THR159_CB | 0.102009173 | 0.317053685 |
| ARG33_CB_THR30_CB | 0.111676558 | 0.149423021 |
| ILE34_CB_THR30_CB | 0.009613371 | 0.005862642 |
| ALA2_CB_THR4_CB | 0.02194781 | 0.031920973 |
| LEU3_CB_THR4_CB | 0.011900607 | 0.102386595 |
| ALA2_CB_THR68_CB | 0.162540396 | 0.299442548 |
| LEU3_CB_THR68_CB | 0.093666937 | 0.355925607 |

**Table S3.** Feature-diverse tICA features and correlations – APO transport (cont.)

| System | APO | |
|---|---|---|
| # of features | 113 | |
| Feature | \| tIC1 correlation (*r*) \| | \| tIC2 correlation (*r*) \| |
| LYS65_CB_THR68_CB | 0.066612068 | 0.107116391 |
| MET1_CB_THR68_CB | 0.080605784 | 0.268049331 |
| ASN76_CB_TRP58_CB | 0.004697473 | 0.044513855 |
| GLY13_CA_TYR183_CB | 0.055201818 | 0.029636509 |
| GLY16_CA_TYR183_CB | 0.131529625 | 0.048149791 |
| TRP180_CB_TYR183_CB | 0.0336046 | 0.128075837 |
| PHE213_CB_TYR210_CB | 0.463148955 | 0.559576133 |
| PHE90_CB_TYR48_CB | 0.10322662 | 0.036750219 |
| CYS138_CB_TYR61_CB | 0.111431654 | 0.035868376 |
| TRP58_CB_TYR61_CB | 0.036068481 | 0.023942738 |
| PHE90_CB_TYR86_CB | 0.012633681 | 0.013574101 |
| PRO27_CB_TYR86_CB | 0.215811336 | 0.124877539 |
| TYR48_CB_TYR86_CB | 0.110001381 | 0.110780415 |
| CYS138_CB_VAL139_CB | 0.004099522 | 0.021729298 |
| SER142_CB_VAL139_CB | 0.043849454 | 0.12236406 |
| ALA51_CB_VAL145_CB | 0.070223802 | 0.082133505 |
| SER50_CB_VAL145_CB | 0.11992962 | 0.086169632 |
| PHE164_CB_VAL155_CB | 0.313404658 | 0.139621621 |
| ASP189_CB_VAL192_CB | 0.108040286 | 0.097034136 |
| GLY184_CA_VAL192_CB | 0.035664913 | 0.027689624 |

**Table S4.** Feature-diverse tICA features and correlations – SUC transport

| System | SUC | |
|---|---|---|
| # of features | 1002 | |
| Feature | \| tIC1 correlation (*r*) \| | \| tIC2 correlation (*r*) \| |
| theta_C2_xy | 0.000507919 | 0.002644896 |
| theta_C2_xz | 0.00291462 | 0.012894096 |
| theta_C2_yz | 0.00440776 | 0.017881753 |
| theta_C5_xy | 0.001377873 | 0.000551382 |
| theta_C5_xz | 0.001352212 | 0.000116881 |
| theta_C5_yz | 0.003088558 | 0.001535354 |
| theta_C6_xy | 0.005093511 | 0.000119118 |
| theta_C6_xz | 0.001237996 | 0.000809024 |
| theta_C6_yz | 0.000345341 | 0.000849222 |
| theta_C9_xy | 0.006610436 | 0.006561418 |
| theta_C9_xz | 0.004742509 | 0.005432844 |
| theta_C9_yz | 0.003056355 | 0.003393752 |
| theta_C10_xy | 0.000427391 | 0.001640712 |
| theta_C10_xz | 0.004870096 | 0.003488025 |
| theta_C10_yz | 0.008033599 | 0.004835636 |
| theta_C11_xy | 0.002115493 | 0.004907802 |
| theta_C11_xz | 0.002709805 | 0.000281608 |
| theta_C11_yz | 0.003080475 | 6.90438E-05 |
| theta_C12_xy | 0.000134135 | 0.000700641 |
| theta_C12_xz | 0.00051783 | 0.000724687 |
| theta_C12_yz | 0.003007091 | 0.000332507 |
| theta_O5_xy | 0.001699923 | 0.001449819 |
| theta_O5_xz | 0.000302155 | 0.002224304 |
| theta_O5_yz | 0.000912033 | 0.003497133 |
| theta_O6_xy | 0.006203179 | 0.000253431 |
| theta_O6_xz | 0.001870487 | 0.000151261 |
| theta_O6_yz | 0.003486647 | 0.001991186 |
| theta_O9_xy | 0.000895247 | 0.000207144 |
| theta_O9_xz | 0.001101415 | 0.000382445 |
| theta_O9_yz | 0.000671666 | 0.001621664 |
| GLY203_CA_ALA177_CB | 0.044513131 | 0.009097411 |

**Table S4.** Feature-diverse tICA features and correlations – SUC transport (cont.)

| System | SUC | |
|---|---|---|
| # of features | 1002 | |
| Feature | I tIC1 correlation (*r*) I | I tIC2 correlation (*r*) I |
| LEU173_CB_ALA177_CB | 0.06242024 | 0.013409084 |
| SER176_CB_ALA177_CB | 0.025813336 | 0.018290445 |
| MET1_CB_ALA2_CB | 0.022851916 | 0.008695539 |
| ALA177_CB_ALA200_CB | 0.12071087 | 0.001401036 |
| GLY199_CA_ALA200_CB | 0.011537573 | 0.002176687 |
| GLY203_CA_ALA200_CB | 0.084034806 | 0.006745332 |
| PRO47_CB_ALA51_CB | 0.032842342 | 0.005678437 |
| SER50_CB_ALA51_CB | 0.143112657 | 0.009697914 |
| TYR48_CB_ALA51_CB | 0.016215349 | 0.003433758 |
| ALA51_CB_ALA55_CB | 0.011748511 | 0.012227936 |
| GLU83_CB_ALA55_CB | 0.03977742 | 0.007103814 |
| GLY79_CA_ALA55_CB | 0.008717022 | 0.039221475 |
| SER54_CB_ALA55_CB | 0.239186752 | 0.026901369 |
| LEU72_CB_ALA62_CB | 0.333067767 | 0.001560938 |
| TRP58_CB_ALA62_CB | 0.187074934 | 0.004009901 |
| TYR61_CB_ALA62_CB | 0.079930472 | 0.000634697 |
| GLU41_CB_ARG33_CB | 0.050503599 | 0.342025252 |
| ILE75_CB_ASN17_CB | 0.190363124 | 0.089835222 |
| SER20_CB_ASN17_CB | 0.338057581 | 0.001513129 |
| ALA177_CB_ASN196_CB | 0.07763582 | 0.033797943 |
| ALA200_CB_ASN196_CB | 0.055220923 | 0.005603244 |
| GLY199_CA_ASN196_CB | 0.023567961 | 0.085286766 |
| TRP180_CB_ASN196_CB | 0.085372131 | 0.009993466 |
| ALA2_CB_ASN5_CB | 0.232032118 | 0.16028043 |
| MET1_CB_ASN5_CB | 0.13434621 | 0.147244783 |
| THR4_CB_ASN5_CB | 0.080995968 | 0.011766275 |
| ALA55_CB_ASN76_CB | 0.025821351 | 0.003235421 |
| GLY79_CA_ASN76_CB | 0.062588638 | 0.032699352 |
| GLY184_CA_ASP189_CB | 0.048320981 | 0.018043818 |
| ILE187_CB_ASP189_CB | 0.055240415 | 0.027931345 |
| LYS65_CB_ASP189_CB | 0.797909801 | 0.019807188 |

**Table S4.** Feature-diverse tICA features and correlations – SUC transport (cont.)

| System | SUC | |
|---|---|---|
| # of features | 1002 | |
| **Feature** | **\| tIC1 correlation (*r*) \|** | **\| tIC2 correlation (*r*) \|** |
| SER142_CB_CYS138_CB | 0.140175152 | 0.050029632 |
| TRP58_CB_CYS138_CB | 0.222487198 | 0.027971643 |
| GLY42_CA_GLN44_CB | 0.034758513 | 0.044287818 |
| PHE43_CB_GLN44_CB | 0.055865157 | 0.125343878 |
| GLY135_CA_GLU131_CB | 0.111069276 | 0.012938435 |
| LYS65_CB_GLU131_CB | 0.20807438 | 0.064365273 |
| GLU41_CB_GLU163_CB | 0.526004232 | 0.078883929 |
| SER161_CB_GLU163_CB | 0.038498986 | 0.008651042 |
| VAL162_CB_GLU163_CB | 0.010520401 | 0.011588959 |
| SER39_CB_GLU41_CB | 0.267130866 | 0.233035607 |
| THR40_CB_GLU41_CB | 0.164067042 | 0.252040498 |
| ALA51_CB_GLU83_CB | 0.021809176 | 0.02623884 |
| TYR48_CB_GLU83_CB | 0.046221356 | 0.010230596 |
| TYR86_CB_GLU83_CB | 0.023709995 | 0.001117892 |
| GLY16_CA_GLY13_CA | 0.211955026 | 0.02415925 |
| CYS138_CB_GLY135_CA | 0.432438357 | 0.072945749 |
| VAL139_CB_GLY135_CA | 0.185821249 | 0.073233119 |
| ILE75_CB_GLY16_CA | 0.89373789 | 0.023575092 |
| SER20_CB_GLY16_CA | 0.202247513 | 0.022068111 |
| TRP180_CB_GLY184_CA | 0.047168285 | 0.010133215 |
| TYR183_CB_GLY184_CA | 0.074075239 | 0.037496993 |
| ALA177_CB_GLY199_CA | 0.100419556 | 0.003168393 |
| LEU173_CB_GLY199_CA | 0.039827563 | 0.012987373 |
| PHE146_CB_GLY199_CA | 0.259358378 | 0.022192728 |
| LEU173_CB_GLY203_CA | 0.527873807 | 0.007919872 |
| PHE146_CB_GLY203_CA | 0.23779852 | 0.003892765 |
| GLU41_CB_GLY42_CA | 0.12362205 | 0.004158185 |
| THR40_CB_GLY42_CA | 0.097543639 | 0.095855793 |
| ALA51_CB_GLY79_CA | 0.08884127 | 0.005768293 |
| PHE24_CB_GLY79_CA | 0.543254432 | 0.031584804 |
| MET165_CB_ILE152_CB | 0.031132536 | 0.045155529 |

**Table S4.** Feature-diverse tICA features and correlations – SUC transport (cont.)

| System | SUC | |
|---|---|---|
| # of features | 1002 | |
| Feature | \| tIC1 correlation (*r*) \| | \| tIC2 correlation (*r*) \| |
| PHE164_CB_ILE152_CB | 0.234135535 | 0.029253892 |
| PHE43_CB_ILE152_CB | 0.311487551 | 0.22582445 |
| SER151_CB_ILE152_CB | 0.026688255 | 0.061814361 |
| VAL155_CB_ILE152_CB | 0.07497249 | 0.004680508 |
| ALA2_CB_ILE187_CB | 0.402157214 | 0.029660903 |
| ASN5_CB_ILE187_CB | 0.270833447 | 0.048019525 |
| LEU3_CB_ILE187_CB | 0.116432765 | 0.240747177 |
| LYS65_CB_ILE187_CB | 0.783740806 | 0.023722147 |
| MET1_CB_ILE187_CB | 0.467433089 | 0.03805153 |
| THR4_CB_ILE187_CB | 0.138955964 | 0.1254702 |
| THR68_CB_ILE187_CB | 0.770071689 | 0.048535597 |
| ARG33_CB_ILE34_CB | 0.174095368 | 0.098405618 |
| GLU41_CB_ILE34_CB | 0.015232098 | 0.331770576 |
| THR40_CB_ILE34_CB | 0.058864433 | 0.041740444 |
| SER20_CB_ILE75_CB | 0.503477657 | 0.09091451 |
| PRO166_CB_LEU169_CB | 0.076687935 | 0.041558728 |
| LEU169_CB_LEU173_CB | 0.241965294 | 0.04351417 |
| ALA2_CB_LEU3_CB | 0.130722205 | 0.01012673 |
| GLY13_CA_LEU3_CB | 0.548484987 | 0.00222202 |
| MET1_CB_LEU3_CB | 0.015093287 | 0.001299378 |
| ASN17_CB_LEU71_CB | 0.255236078 | 0.007305336 |
| GLY13_CA_LEU71_CB | 0.902584068 | 0.011503472 |
| GLY16_CA_LEU71_CB | 0.914822429 | 0.003269903 |
| ILE75_CB_LEU71_CB | 0.433249171 | 0.09780171 |
| LEU72_CB_LEU71_CB | 0.697791919 | 0.005021256 |
| ASN76_CB_LEU72_CB | 0.102816641 | 0.038209831 |
| ILE75_CB_LEU72_CB | 0.461598386 | 0.019256588 |
| TRP58_CB_LEU72_CB | 0.34168922 | 0.031008695 |
| GLY135_CA_LYS132_CB | 0.163117071 | 0.016420298 |
| ARG33_CB_LYS37_CB | 0.058103507 | 0.112627991 |
| SER39_CB_LYS37_CB | 0.034302187 | 0.117731054 |

**Table S4.** Feature-diverse tICA features and correlations – SUC transport (cont.)

| System | SUC | |
|---|---|---|
| # of features | 1002 | |
| Feature | \| tIC1 correlation (*r*) \| | \| tIC2 correlation (*r*) \| |
| SER39_CB_LYS38_CB | 0.00430046 | 0.019124897 |
| ALA62_CB_LYS65_CB | 0.309479154 | 0.089794539 |
| MET1_CB_LYS65_CB | 0.101966366 | 0.011932714 |
| TYR61_CB_LYS65_CB | 0.449942341 | 0.038745638 |
| GLY13_CA_MET1_CB | 0.360744706 | 0.041968061 |
| LEU71_CB_MET1_CB | 0.157698001 | 0.084999225 |
| PHE164_CB_MET165_CB | 0.060840439 | 0.01639184 |
| VAL162_CB_MET165_CB | 0.165653233 | 0.012773927 |
| PRO149_CB_PHE146_CB | 0.046006769 | 0.022326019 |
| GLU163_CB_PHE164_CB | 0.07272902 | 0.015477832 |
| GLY42_CA_PHE164_CB | 0.419275872 | 0.058172763 |
| PHE43_CB_PHE164_CB | 0.458514425 | 0.051335762 |
| SER161_CB_PHE164_CB | 0.10909017 | 0.031087727 |
| ARG160_CB_PHE213_CB | 0.105771362 | 0.001874181 |
| SER161_CB_PHE213_CB | 0.249080224 | 0.002975936 |
| VAL162_CB_PHE213_CB | 0.021499908 | 0.038478918 |
| VAL23_CB_PHE24_CB | 0.302368699 | 0.023808431 |
| GLU41_CB_PHE43_CB | 0.350977656 | 0.176482401 |
| GLY42_CA_PHE43_CB | 0.193718324 | 0.142156793 |
| THR40_CB_PHE43_CB | 0.207500952 | 0.530408808 |
| ILE152_CB_PRO149_CB | 0.126310979 | 0.023274713 |
| PHE43_CB_PRO149_CB | 0.161288402 | 0.245394203 |
| PRO47_CB_PRO149_CB | 0.308933921 | 0.018619188 |
| GLY42_CA_PRO166_CB | 0.478264279 | 0.214155605 |
| PHE43_CB_PRO166_CB | 0.580840831 | 0.09152994 |
| SER142_CB_PRO195_CB | 0.010868757 | 0.006405824 |
| VAL139_CB_PRO195_CB | 0.16201219 | 0.037156301 |
| THR30_CB_PRO27_CB | 0.14074592 | 0.021634238 |
| GLN44_CB_PRO47_CB | 0.24603655 | 0.116621451 |
| PHE43_CB_PRO47_CB | 0.198077157 | 0.269896784 |
| SER54_CB_SER142_CB | 0.324931144 | 0.069748124 |

**Table S4.** Feature-diverse tICA features and correlations – SUC transport (cont.)

| System | SUC | |
|---|---|---|
| # of features | 1002 | |
| Feature | ǀ tIC1 correlation (*r*) ǀ | ǀ tIC2 correlation (*r*) ǀ |
| VAL145_CB_SER142_CB | 0.305412688 | 0.047092404 |
| GLN44_CB_SER151_CB | 0.443332073 | 0.059402582 |
| ARG160_CB_SER161_CB | 0.139360503 | 0.016459262 |
| LEU173_CB_SER176_CB | 0.310470711 | 0.031281059 |
| SER20_CB_SER176_CB | 0.501796579 | 0.06757347 |
| VAL23_CB_SER176_CB | 0.363673889 | 0.04339195 |
| VAL23_CB_SER20_CB | 0.330529392 | 0.001901287 |
| PRO47_CB_SER50_CB | 0.189762269 | 0.039384536 |
| ALA51_CB_SER54_CB | 0.419342545 | 0.040178133 |
| SER50_CB_SER54_CB | 0.260869444 | 0.069318947 |
| VAL145_CB_SER54_CB | 0.324037081 | 0.00124315 |
| GLU163_CB_THR159_CB | 0.097173785 | 0.005656187 |
| SER161_CB_THR159_CB | 0.105656594 | 0.006314615 |
| ARG33_CB_THR30_CB | 0.230185433 | 0.086410677 |
| GLU41_CB_THR30_CB | 0.083403717 | 0.365977418 |
| GLY42_CA_THR30_CB | 0.206898446 | 0.35640617 |
| ILE34_CB_THR30_CB | 0.020691969 | 0.012516384 |
| ALA2_CB_THR4_CB | 0.164955358 | 0.128102615 |
| LEU3_CB_THR4_CB | 0.064945137 | 0.118946328 |
| MET1_CB_THR4_CB | 0.234519 | 0.006447078 |
| ALA2_CB_THR68_CB | 0.103768188 | 0.003421895 |
| LEU3_CB_THR68_CB | 0.204724403 | 0.156235591 |
| LEU71_CB_THR68_CB | 0.63806983 | 0.024118941 |
| LYS65_CB_THR68_CB | 0.652503109 | 0.005126568 |
| THR4_CB_THR68_CB | 0.394551641 | 0.029456213 |
| ASN76_CB_TRP58_CB | 0.254720792 | 0.015897374 |
| GLY13_CA_TYR183_CB | 0.235177407 | 0.003992889 |
| GLY16_CA_TYR183_CB | 0.236142636 | 0.039414741 |
| TRP180_CB_TYR183_CB | 0.089797275 | 0.032879966 |
| LEU209_CB_TYR210_CB | 0.110012265 | 0.006882411 |
| MET165_CB_TYR210_CB | 0.170848251 | 0.033304907 |

**Table S4.** Feature-diverse tICA features and correlations – SUC transport (cont.)

| System | SUC | |
|---|---|---|
| # of features | 1002 | |
| Feature | l tIC1 correlation (*r*) l | l tIC2 correlation (*r*) l |
| PHE213_CB_TYR210_CB | 0.139497089 | 0.005871226 |
| PHE90_CB_TYR48_CB | 0.087968389 | 0.132696692 |
| CYS138_CB_TYR61_CB | 0.615613584 | 0.079349907 |
| TRP58_CB_TYR61_CB | 0.295609252 | 0.044635406 |
| PHE90_CB_TYR86_CB | 0.094791212 | 0.075191266 |
| PRO27_CB_TYR86_CB | 0.22573854 | 0.023550817 |
| TYR48_CB_TYR86_CB | 0.100324671 | 0.028953383 |
| CYS138_CB_VAL139_CB | 0.306983518 | 0.079688833 |
| SER142_CB_VAL139_CB | 0.020044986 | 0.002218842 |
| ALA51_CB_VAL145_CB | 0.257031756 | 0.019532142 |
| SER50_CB_VAL145_CB | 0.288710926 | 0.05082581 |
| PHE164_CB_VAL155_CB | 0.08371074 | 0.022380705 |
| ASP189_CB_VAL192_CB | 0.205042148 | 0.002996213 |
| GLY184_CA_VAL192_CB | 0.166063686 | 0.002586362 |
| TRP180_CB_VAL192_CB | 0.162648833 | 0.015767534 |
| TYR183_CB_VAL192_CB | 0.226501278 | 0.015928664 |
| ALA177_CB_C2 | 0.121837771 | 0.77764877 |
| ALA2_CB_C2 | 0.5940792 | 0.191759249 |
| ALA200_CB_C2 | 0.029206228 | 0.777901037 |
| ALA51_CB_C2 | 0.477168466 | 0.434366306 |
| ALA55_CB_C2 | 0.214705025 | 0.808148655 |
| ALA62_CB_C2 | 0.473656517 | 0.690278212 |
| ARG160_CB_C2 | 0.502544473 | 0.441781125 |
| ARG33_CB_C2 | 0.491108254 | 0.395774244 |
| ASN17_CB_C2 | 0.377966793 | 0.705174201 |
| ASN196_CB_C2 | 0.203667183 | 0.829207838 |
| ASN5_CB_C2 | 0.460550097 | 0.265834285 |
| ASN76_CB_C2 | 0.011171066 | 0.888573941 |
| ASP189_CB_C2 | 0.639038284 | 0.521286263 |
| CYS138_CB_C2 | 0.655769267 | 0.413281836 |
| GLN44_CB_C2 | 0.497538383 | 0.407934383 |

**Table S4.** Feature-diverse tICA features and correlations – SUC transport (cont.)

| System | SUC | |
|---|---|---|
| # of features | 1002 | |
| Feature | \| tIC1 correlation (*r*) \| | \| tIC2 correlation (*r*) \| |
| GLU131_CB_C2 | 0.645404358 | 0.381588373 |
| GLU163_CB_C2 | 0.473731918 | 0.469283744 |
| GLU41_CB_C2 | 0.474254457 | 0.358836576 |
| GLU83_CB_C2 | 0.485828807 | 0.51181523 |
| GLY13_CA_C2 | 0.542864854 | 0.605167564 |
| GLY135_CA_C2 | 0.647572357 | 0.420376696 |
| GLY16_CA_C2 | 0.40340206 | 0.756138302 |
| GLY184_CA_C2 | 0.563581391 | 0.66015131 |
| GLY199_CA_C2 | 0.21806704 | 0.736352146 |
| GLY203_CA_C2 | 0.420702888 | 0.464452812 |
| GLY42_CA_C2 | 0.528644434 | 0.326745764 |
| GLY79_CA_C2 | 0.305405676 | 0.800294528 |
| ILE152_CB_C2 | 0.546493284 | 0.367251839 |
| ILE187_CB_C2 | 0.600418583 | 0.460752161 |
| ILE34_CB_C2 | 0.557897465 | 0.292365432 |
| ILE75_CB_C2 | 0.030911636 | 0.777318504 |
| LEU169_CB_C2 | 0.533393377 | 0.186239788 |
| LEU173_CB_C2 | 0.385022782 | 0.393991331 |
| LEU209_CB_C2 | 0.574509063 | 0.146051946 |
| LEU3_CB_C2 | 0.631522116 | 0.229103165 |
| LEU71_CB_C2 | 0.266748303 | 0.658777461 |
| LEU72_CB_C2 | 0.399631426 | 0.679129373 |
| LYS132_CB_C2 | 0.587456346 | 0.423640252 |
| LYS37_CB_C2 | 0.500386753 | 0.394613877 |
| LYS38_CB_C2 | 0.529705431 | 0.303577677 |
| LYS65_CB_C2 | 0.672877475 | 0.37710321 |
| MET1_CB_C2 | 0.607293944 | 0.189326001 |
| MET165_CB_C2 | 0.542608374 | 0.362760194 |
| PHE146_CB_C2 | 0.385853193 | 0.390084851 |
| PHE164_CB_C2 | 0.481084896 | 0.496334174 |
| PHE213_CB_C2 | 0.543287031 | 0.352671109 |

**Table S4.** Feature-diverse tICA features and correlations – SUC transport (cont.)

| System | SUC | |
|---|---|---|
| # of features | 1002 | |
| Feature | I tIC1 correlation (*r*) I | I tIC2 correlation (*r*) I |
| PHE24_CB_C2 | 0.416841907 | 0.589544353 |
| PHE43_CB_C2 | 0.581027351 | 0.267069491 |
| PHE90_CB_C2 | 0.593714413 | 0.070489217 |
| PRO149_CB_C2 | 0.535817867 | 0.119504217 |
| PRO166_CB_C2 | 0.445771245 | 0.377225079 |
| PRO195_CB_C2 | 0.256673419 | 0.756947704 |
| PRO27_CB_C2 | 0.551582822 | 0.243493251 |
| PRO47_CB_C2 | 0.527999931 | 0.155759431 |
| SER142_CB_C2 | 0.28912037 | 0.709355682 |
| SER151_CB_C2 | 0.522581027 | 0.393306647 |
| SER161_CB_C2 | 0.503371926 | 0.482523955 |
| SER176_CB_C2 | 0.165333062 | 0.783420809 |
| SER20_CB_C2 | 0.066169122 | 0.785906856 |
| SER39_CB_C2 | 0.503612723 | 0.412353809 |
| SER50_CB_C2 | 0.337755655 | 0.437993753 |
| SER54_CB_C2 | 0.076739663 | 0.759527583 |
| THR159_CB_C2 | 0.46871306 | 0.511132849 |
| THR30_CB_C2 | 0.51361197 | 0.346877176 |
| THR4_CB_C2 | 0.438798682 | 0.319929935 |
| THR40_CB_C2 | 0.53414587 | 0.386908353 |
| THR68_CB_C2 | 0.38711917 | 0.458649482 |
| TRP180_CB_C2 | 0.411822933 | 0.715135853 |
| TRP58_CB_C2 | 0.373521095 | 0.784117898 |
| TYR183_CB_C2 | 0.544943955 | 0.683643068 |
| TYR210_CB_C2 | 0.545717565 | 0.091092688 |
| TYR48_CB_C2 | 0.573422757 | 0.157566412 |
| TYR61_CB_C2 | 0.641637599 | 0.574988632 |
| TYR86_CB_C2 | 0.603400809 | 0.057173674 |
| VAL139_CB_C2 | 0.55883661 | 0.462578432 |
| VAL145_CB_C2 | 0.354658533 | 0.553986355 |
| VAL155_CB_C2 | 0.475108429 | 0.503569506 |

**Table S4.** Feature-diverse tICA features and correlations – SUC transport (cont.)

| System | SUC | |
|---|---|---|
| # of features | 1002 | |
| Feature | l tIC1 correlation (*r*) l | l tIC2 correlation (*r*) l |
| VAL162_CB_C2 | 0.524707293 | 0.373714183 |
| VAL192_CB_C2 | 0.632018715 | 0.55787266 |
| VAL23_CB_C2 | 0.337496038 | 0.641839446 |
| ALA177_CB_C5 | 0.408569419 | 0.667874294 |
| ALA2_CB_C5 | 0.693761253 | 0.081630066 |
| ALA200_CB_C5 | 0.35389835 | 0.745709135 |
| ALA51_CB_C5 | 0.563645088 | 0.453706557 |
| ALA55_CB_C5 | 0.085580296 | 0.66596805 |
| ALA62_CB_C5 | 0.800938978 | 0.268853575 |
| ARG160_CB_C5 | 0.708774453 | 0.291165114 |
| ARG33_CB_C5 | 0.610659261 | 0.36655206 |
| ASN17_CB_C5 | 0.730817069 | 0.443254621 |
| ASN196_CB_C5 | 0.1264263 | 0.76958521 |
| ASN5_CB_C5 | 0.709050232 | 0.149271836 |
| ASN76_CB_C5 | 0.449162467 | 0.615528555 |
| ASP189_CB_C5 | 0.776048832 | 0.357219557 |
| CYS138_CB_C5 | 0.634164143 | 0.505693243 |
| GLN44_CB_C5 | 0.611381571 | 0.378833285 |
| GLU131_CB_C5 | 0.816075303 | 0.263723722 |
| GLU163_CB_C5 | 0.658057994 | 0.388811987 |
| GLU41_CB_C5 | 0.631981345 | 0.336748038 |
| GLU83_CB_C5 | 0.567228885 | 0.436937512 |
| GLY13_CA_C5 | 0.839044225 | 0.312343053 |
| GLY135_CA_C5 | 0.748002552 | 0.396482799 |
| GLY16_CA_C5 | 0.646869632 | 0.563417714 |
| GLY184_CA_C5 | 0.641106018 | 0.542572033 |
| GLY199_CA_C5 | 0.484476416 | 0.680272543 |
| GLY203_CA_C5 | 0.647284571 | 0.602620503 |
| GLY42_CA_C5 | 0.666358943 | 0.326334559 |
| GLY79_CA_C5 | 0.153478869 | 0.701019555 |
| ILE152_CB_C5 | 0.742783994 | 0.22407676 |

**Table S4.** Feature-diverse tICA features and correlations – SUC transport (cont.)

| System | SUC | |
|---|---|---|
| # of features | 1002 | |
| Feature | \| tIC1 correlation (*r*) \| | \| tIC2 correlation (*r*) \| |
| ILE187_CB_C5 | 0.817404735 | 0.262311091 |
| ILE34_CB_C5 | 0.66930179 | 0.287865978 |
| ILE75_CB_C5 | 0.555775483 | 0.514373058 |
| LEU169_CB_C5 | 0.702566944 | 0.005740881 |
| LEU173_CB_C5 | 0.637586643 | 0.586059351 |
| LEU209_CB_C5 | 0.807891109 | 0.079461516 |
| LEU3_CB_C5 | 0.769222165 | 0.105806825 |
| LEU71_CB_C5 | 0.68473489 | 0.270614127 |
| LEU72_CB_C5 | 0.747497279 | 0.194716744 |
| LYS132_CB_C5 | 0.693617079 | 0.384530442 |
| LYS37_CB_C5 | 0.629731854 | 0.374115247 |
| LYS38_CB_C5 | 0.653646544 | 0.302477281 |
| LYS65_CB_C5 | 0.843652932 | 0.174204596 |
| MET1_CB_C5 | 0.658001595 | 0.078966795 |
| MET165_CB_C5 | 0.698396754 | 0.212776394 |
| PHE146_CB_C5 | 0.681561516 | 0.588243931 |
| PHE164_CB_C5 | 0.62196733 | 0.430833802 |
| PHE213_CB_C5 | 0.768082218 | 0.176496229 |
| PHE24_CB_C5 | 0.466484272 | 0.545872104 |
| PHE43_CB_C5 | 0.683233027 | 0.266644055 |
| PHE90_CB_C5 | 0.714205204 | 0.097686984 |
| PRO149_CB_C5 | 0.775321956 | 0.184390808 |
| PRO166_CB_C5 | 0.646755123 | 0.251930192 |
| PRO195_CB_C5 | 0.165262443 | 0.761486515 |
| PRO27_CB_C5 | 0.644661684 | 0.187649655 |
| PRO47_CB_C5 | 0.71142905 | 0.019786713 |
| SER142_CB_C5 | 0.172955605 | 0.687408801 |
| SER151_CB_C5 | 0.626492155 | 0.316911767 |
| SER161_CB_C5 | 0.687994999 | 0.373047726 |
| SER176_CB_C5 | 0.41264755 | 0.713735292 |
| SER20_CB_C5 | 0.061309364 | 0.739561691 |

**Table S4.** Feature-diverse tICA features and correlations – SUC transport (cont.)

| System | SUC | |
|---|---|---|
| # of features | 1002 | |
| Feature | I tIC1 correlation (*r*) I | I tIC2 correlation (*r*) I |
| SER39_CB_C5 | 0.626539869 | 0.385600975 |
| SER50_CB_C5 | 0.580301887 | 0.456772678 |
| SER54_CB_C5 | 0.265483966 | 0.689451354 |
| THR159_CB_C5 | 0.644741804 | 0.416396276 |
| THR30_CB_C5 | 0.611690267 | 0.30843571 |
| THR4_CB_C5 | 0.679512907 | 0.153501294 |
| THR40_CB_C5 | 0.663556305 | 0.367332662 |
| THR68_CB_C5 | 0.730634351 | 0.200957874 |
| TRP180_CB_C5 | 0.370875472 | 0.621509937 |
| TRP58_CB_C5 | 0.697013406 | 0.383621415 |
| TYR183_CB_C5 | 0.705189403 | 0.476590084 |
| TYR210_CB_C5 | 0.788977191 | 0.122117571 |
| TYR48_CB_C5 | 0.697845033 | 0.149786291 |
| TYR61_CB_C5 | 0.797612399 | 0.336157856 |
| TYR86_CB_C5 | 0.706014415 | 0.032482951 |
| VAL139_CB_C5 | 0.430862174 | 0.645620802 |
| VAL145_CB_C5 | 0.568587447 | 0.597267451 |
| VAL155_CB_C5 | 0.624948838 | 0.429221801 |
| VAL162_CB_C5 | 0.732778775 | 0.221297112 |
| VAL192_CB_C5 | 0.719083219 | 0.414431331 |
| VAL23_CB_C5 | 0.495525702 | 0.677155264 |
| ALA177_CB_C6 | 0.406534415 | 0.67436181 |
| ALA2_CB_C6 | 0.713535432 | 0.074530219 |
| ALA200_CB_C6 | 0.338366654 | 0.750417526 |
| ALA51_CB_C6 | 0.598225434 | 0.443815483 |
| ALA55_CB_C6 | 0.152813173 | 0.668618778 |
| ALA62_CB_C6 | 0.782525844 | 0.231934863 |
| ARG160_CB_C6 | 0.722090191 | 0.25343511 |
| ARG33_CB_C6 | 0.648822757 | 0.343538279 |
| ASN17_CB_C6 | 0.664515423 | 0.489074489 |
| ASN196_CB_C6 | 0.054158101 | 0.782630185 |

**Table S4.** Feature-diverse tICA features and correlations – SUC transport (cont.)

| System | SUC | |
|---|---|---|
| # of features | 1002 | |
| Feature | I tIC1 correlation (*r*) I | I tIC2 correlation (*r*) I |
| ASN5_CB_C6 | 0.723444571 | 0.129547867 |
| ASN76_CB_C6 | 0.405763047 | 0.645080569 |
| ASP189_CB_C6 | 0.822755856 | 0.292178287 |
| CYS138_CB_C6 | 0.7085358 | 0.428413649 |
| GLN44_CB_C6 | 0.599336446 | 0.392211326 |
| GLU131_CB_C6 | 0.850951112 | 0.214993129 |
| GLU163_CB_C6 | 0.686038108 | 0.352425464 |
| GLU41_CB_C6 | 0.669946456 | 0.318162493 |
| GLU83_CB_C6 | 0.62437161 | 0.409087322 |
| GLY13_CA_C6 | 0.815286791 | 0.29246222 |
| GLY135_CA_C6 | 0.805289717 | 0.321734377 |
| GLY16_CA_C6 | 0.616743009 | 0.59568559 |
| GLY184_CA_C6 | 0.723505515 | 0.48418378 |
| GLY199_CA_C6 | 0.456953106 | 0.655854206 |
| GLY203_CA_C6 | 0.641534868 | 0.612420419 |
| GLY42_CA_C6 | 0.688837378 | 0.317150666 |
| GLY79_CA_C6 | 0.263239197 | 0.707993554 |
| ILE152_CB_C6 | 0.770622587 | 0.181090849 |
| ILE187_CB_C6 | 0.849261706 | 0.221100011 |
| ILE34_CB_C6 | 0.702554775 | 0.273961164 |
| ILE75_CB_C6 | 0.485462369 | 0.582929275 |
| LEU169_CB_C6 | 0.746534213 | 0.078302361 |
| LEU173_CB_C6 | 0.628239637 | 0.597591954 |
| LEU209_CB_C6 | 0.81489391 | 0.121847454 |
| LEU3_CB_C6 | 0.765476478 | 0.092060146 |
| LEU71_CB_C6 | 0.679522839 | 0.28138362 |
| LEU72_CB_C6 | 0.763341671 | 0.203636108 |
| LYS132_CB_C6 | 0.766288053 | 0.320578956 |
| LYS37_CB_C6 | 0.665709885 | 0.351690682 |
| LYS38_CB_C6 | 0.692841767 | 0.283801777 |
| LYS65_CB_C6 | 0.844193137 | 0.140347426 |

**Table S4.** Feature-diverse tICA features and correlations – SUC transport (cont.)

| System | SUC | |
|---|---|---|
| # of features | 1002 | |
| Feature | \| tIC1 correlation (*r*) \| | \| tIC2 correlation (*r*) \| |
| MET1_CB_C6 | 0.680607601 | 0.074439735 |
| MET165_CB_C6 | 0.724933659 | 0.16350898 |
| PHE146_CB_C6 | 0.677393881 | 0.595133483 |
| PHE164_CB_C6 | 0.641530723 | 0.402627974 |
| PHE213_CB_C6 | 0.785025355 | 0.129917751 |
| PHE24_CB_C6 | 0.558824571 | 0.552786738 |
| PHE43_CB_C6 | 0.674478636 | 0.293220087 |
| PHE90_CB_C6 | 0.739301965 | 0.116814756 |
| PRO149_CB_C6 | 0.792948337 | 0.232173421 |
| PRO166_CB_C6 | 0.699610229 | 0.189775159 |
| PRO195_CB_C6 | 0.06092373 | 0.747346994 |
| PRO27_CB_C6 | 0.675204629 | 0.180372235 |
| PRO47_CB_C6 | 0.737605685 | 0.054713433 |
| SER142_CB_C6 | 0.082858769 | 0.712032444 |
| SER151_CB_C6 | 0.683096901 | 0.295094859 |
| SER161_CB_C6 | 0.708426572 | 0.335109162 |
| SER176_CB_C6 | 0.445379863 | 0.704298743 |
| SER20_CB_C6 | 0.092422807 | 0.753659115 |
| SER39_CB_C6 | 0.665904022 | 0.370490292 |
| SER50_CB_C6 | 0.621836022 | 0.448112128 |
| SER54_CB_C6 | 0.281433133 | 0.690112763 |
| THR159_CB_C6 | 0.660707614 | 0.382344235 |
| THR30_CB_C6 | 0.63912663 | 0.292740432 |
| THR4_CB_C6 | 0.676596482 | 0.130458025 |
| THR40_CB_C6 | 0.686913385 | 0.366221692 |
| THR68_CB_C6 | 0.75461095 | 0.186277157 |
| TRP180_CB_C6 | 0.453198966 | 0.636050564 |
| TRP58_CB_C6 | 0.724091534 | 0.392418932 |
| TYR183_CB_C6 | 0.757813707 | 0.460373903 |
| TYR210_CB_C6 | 0.803924879 | 0.175119159 |
| TYR48_CB_C6 | 0.729676799 | 0.11272061 |

**Table S4.** Feature-diverse tICA features and correlations – SUC transport (cont.)

| System | SUC | |
|---|---|---|
| # of features | 1002 | |
| Feature | I tIC1 correlation (*r*) I | I tIC2 correlation (*r*) I |
| TYR61_CB_C6 | 0.816165665 | 0.283138376 |
| TYR86_CB_C6 | 0.736147771 | 0.013660636 |
| VAL139_CB_C6 | 0.544244223 | 0.545409333 |
| VAL145_CB_C6 | 0.584330867 | 0.613337748 |
| VAL155_CB_C6 | 0.642515585 | 0.400781844 |
| VAL162_CB_C6 | 0.762408759 | 0.171379631 |
| VAL192_CB_C6 | 0.771222133 | 0.343258699 |
| VAL23_CB_C6 | 0.531080718 | 0.642885251 |
| ALA177_CB_C9 | 0.04733647 | 0.649034677 |
| ALA2_CB_C9 | 0.540677803 | 0.206893912 |
| ALA200_CB_C9 | 0.152434939 | 0.656620019 |
| ALA51_CB_C9 | 0.429885724 | 0.351074608 |
| ALA55_CB_C9 | 0.307596167 | 0.751032098 |
| ALA62_CB_C9 | 0.263598721 | 0.763267584 |
| ARG160_CB_C9 | 0.432261529 | 0.438743475 |
| ARG33_CB_C9 | 0.462252752 | 0.379062286 |
| ASN17_CB_C9 | 0.181684124 | 0.748825865 |
| ASN196_CB_C9 | 0.372558969 | 0.652286181 |
| ASN5_CB_C9 | 0.403294369 | 0.279355189 |
| ASN76_CB_C9 | 0.208000309 | 0.845012941 |
| ASP189_CB_C9 | 0.626819886 | 0.465952056 |
| CYS138_CB_C9 | 0.633957837 | 0.357558198 |
| GLN44_CB_C9 | 0.447961248 | 0.387084573 |
| GLU131_CB_C9 | 0.577890497 | 0.360046283 |
| GLU163_CB_C9 | 0.435966502 | 0.454715049 |
| GLU41_CB_C9 | 0.431091262 | 0.335509424 |
| GLU83_CB_C9 | 0.486774552 | 0.467207401 |
| GLY13_CA_C9 | 0.417390373 | 0.635716481 |
| GLY135_CA_C9 | 0.631066806 | 0.38381735 |
| GLY16_CA_C9 | 0.296273327 | 0.767038097 |
| GLY184_CA_C9 | 0.585093756 | 0.55623489 |

**Table S4.** Feature-diverse tICA features and correlations – SUC transport (cont.)

| System | SUC | |
|---|---|---|
| # of features | 1002 | |
| Feature | \| tIC1 correlation (*r*) \| | \| tIC2 correlation (*r*) \| |
| GLY199_CA_C9 | 0.011396847 | 0.583797356 |
| GLY203_CA_C9 | 0.251660551 | 0.416742309 |
| GLY42_CA_C9 | 0.482427326 | 0.297104553 |
| GLY79_CA_C9 | 0.393010673 | 0.731535351 |
| ILE152_CB_C9 | 0.490141748 | 0.333864725 |
| ILE187_CB_C9 | 0.55737977 | 0.458520506 |
| ILE34_CB_C9 | 0.534834395 | 0.271660936 |
| ILE75_CB_C9 | 0.180464261 | 0.777328525 |
| LEU169_CB_C9 | 0.477413315 | 0.171972205 |
| LEU173_CB_C9 | 0.268166712 | 0.332372993 |
| LEU209_CB_C9 | 0.464424634 | 0.134528539 |
| LEU3_CB_C9 | 0.572571221 | 0.247529582 |
| LEU71_CB_C9 | 0.06295305 | 0.708939443 |
| LEU72_CB_C9 | 0.155581585 | 0.748272369 |
| LYS132_CB_C9 | 0.576115253 | 0.377803231 |
| LYS37_CB_C9 | 0.474964754 | 0.377094825 |
| LYS38_CB_C9 | 0.508381709 | 0.282566662 |
| LYS65_CB_C9 | 0.583588183 | 0.40850517 |
| MET1_CB_C9 | 0.58313615 | 0.217783257 |
| MET165_CB_C9 | 0.490012746 | 0.341962017 |
| PHE146_CB_C9 | 0.178472124 | 0.319809552 |
| PHE164_CB_C9 | 0.442376038 | 0.467780768 |
| PHE213_CB_C9 | 0.465885399 | 0.343466771 |
| PHE24_CB_C9 | 0.458535491 | 0.549694884 |
| PHE43_CB_C9 | 0.524000503 | 0.246401336 |
| PHE90_CB_C9 | 0.566474344 | 0.054495656 |
| PRO149_CB_C9 | 0.457230253 | 0.102180443 |
| PRO166_CB_C9 | 0.407407427 | 0.353247146 |
| PRO195_CB_C9 | 0.471634326 | 0.557701672 |
| PRO27_CB_C9 | 0.514279607 | 0.233052056 |
| PRO47_CB_C9 | 0.463050298 | 0.133650277 |

**Table S4.** Feature-diverse tICA features and correlations – SUC transport (cont.)

| System | SUC | |
|---|---|---|
| # of features | 1002 | |
| Feature | \| tIC1 correlation (*r*) \| | \| tIC2 correlation (*r*) \| |
| SER142_CB_C9 | 0.496877509 | 0.512647598 |
| SER151_CB_C9 | 0.456543728 | 0.364330637 |
| SER161_CB_C9 | 0.455773156 | 0.471097909 |
| SER176_CB_C9 | 0.085919247 | 0.711239416 |
| SER20_CB_C9 | 0.166544428 | 0.724575717 |
| SER39_CB_C9 | 0.473220656 | 0.396164237 |
| SER50_CB_C9 | 0.248446202 | 0.371536889 |
| SER54_CB_C9 | 0.064666722 | 0.708407774 |
| THR159_CB_C9 | 0.422276229 | 0.501005156 |
| THR30_CB_C9 | 0.476314461 | 0.330932067 |
| THR4_CB_C9 | 0.357798146 | 0.345387381 |
| THR40_CB_C9 | 0.50177752 | 0.371790125 |
| THR68_CB_C9 | 0.257387418 | 0.490339717 |
| TRP180_CB_C9 | 0.446041772 | 0.55768261 |
| TRP58_CB_C9 | 0.136914758 | 0.830233386 |
| TYR183_CB_C9 | 0.509359372 | 0.627816536 |
| TYR210_CB_C9 | 0.440274654 | 0.075159149 |
| TYR48_CB_C9 | 0.539618281 | 0.151137709 |
| TYR61_CB_C9 | 0.535533256 | 0.630341843 |
| TYR86_CB_C9 | 0.580289544 | 0.067540689 |
| VAL139_CB_C9 | 0.594252866 | 0.347560825 |
| VAL145_CB_C9 | 0.216832057 | 0.45583228 |
| VAL155_CB_C9 | 0.438347039 | 0.483570483 |
| VAL162_CB_C9 | 0.459046838 | 0.35672278 |
| VAL192_CB_C9 | 0.636179003 | 0.470568923 |
| VAL23_CB_C9 | 0.339429622 | 0.584528641 |
| ALA177_CB_C10 | 0.058133761 | 0.596499544 |
| ALA2_CB_C10 | 0.540213678 | 0.20004048 |
| ALA200_CB_C10 | 0.183681064 | 0.609220025 |
| ALA51_CB_C10 | 0.471643547 | 0.342427326 |
| ALA55_CB_C10 | 0.361627104 | 0.696816135 |

**Table S4.** Feature-diverse tICA features and correlations – SUC transport (cont.)

| System | SUC | |
|---|---|---|
| # of features | 1002 | |
| Feature | l tIC1 correlation (*r*) l | l tIC2 correlation (*r*) l |
| ALA62_CB_C10 | 0.253559114 | 0.74287705 |
| ARG160_CB_C10 | 0.427270946 | 0.427822346 |
| ARG33_CB_C10 | 0.482212679 | 0.357006895 |
| ASN17_CB_C10 | 0.079186541 | 0.759701164 |
| ASN196_CB_C10 | 0.40445499 | 0.6075174 |
| ASN5_CB_C10 | 0.383121713 | 0.251630028 |
| ASN76_CB_C10 | 0.265519249 | 0.818940324 |
| ASP189_CB_C10 | 0.663076076 | 0.406132081 |
| CYS138_CB_C10 | 0.700165855 | 0.313807693 |
| GLN44_CB_C10 | 0.469470674 | 0.365414457 |
| GLU131_CB_C10 | 0.626945251 | 0.327131178 |
| GLU163_CB_C10 | 0.442386765 | 0.439662496 |
| GLU41_CB_C10 | 0.453924058 | 0.315864151 |
| GLU83_CB_C10 | 0.529335 | 0.469385466 |
| GLY13_CA_C10 | 0.34570823 | 0.620924856 |
| GLY135_CA_C10 | 0.688955931 | 0.325106868 |
| GLY16_CA_C10 | 0.183590464 | 0.762325538 |
| GLY184_CA_C10 | 0.615120445 | 0.512489952 |
| GLY199_CA_C10 | 0.042909827 | 0.544584475 |
| GLY203_CA_C10 | 0.237953414 | 0.39631924 |
| GLY42_CA_C10 | 0.501086036 | 0.273814282 |
| GLY79_CA_C10 | 0.43836698 | 0.700103041 |
| ILE152_CB_C10 | 0.489539037 | 0.322212352 |
| ILE187_CB_C10 | 0.570805171 | 0.424346801 |
| ILE34_CB_C10 | 0.564012561 | 0.24009064 |
| ILE75_CB_C10 | 0.228484237 | 0.756408473 |
| LEU169_CB_C10 | 0.494691987 | 0.140909142 |
| LEU173_CB_C10 | 0.272362664 | 0.343541964 |
| LEU209_CB_C10 | 0.454080944 | 0.139598153 |
| LEU3_CB_C10 | 0.556622029 | 0.238378758 |
| LEU71_CB_C10 | 0.017381187 | 0.676983512 |

**Table S4.** Feature-diverse tICA features and correlations – SUC transport (cont.)

| System | SUC | |
|---|---|---|
| # of features | 1002 | |
| Feature | \| tIC1 correlation (*r*) \| | \| tIC2 correlation (*r*) \| |
| LEU72_CB_C10 | 0.116899278 | 0.718164027 |
| LYS132_CB_C10 | 0.641884959 | 0.332159365 |
| LYS37_CB_C10 | 0.49674988 | 0.351228487 |
| LYS38_CB_C10 | 0.532915511 | 0.248125011 |
| LYS65_CB_C10 | 0.59230413 | 0.370360785 |
| MET1_CB_C10 | 0.577249626 | 0.212305515 |
| MET165_CB_C10 | 0.495534016 | 0.325828658 |
| PHE146_CB_C10 | 0.149583937 | 0.261580719 |
| PHE164_CB_C10 | 0.442819159 | 0.447028175 |
| PHE213_CB_C10 | 0.46399927 | 0.3347855 |
| PHE24_CB_C10 | 0.51567941 | 0.565262049 |
| PHE43_CB_C10 | 0.540194607 | 0.216880337 |
| PHE90_CB_C10 | 0.599284677 | 0.017160303 |
| PRO149_CB_C10 | 0.450482032 | 0.109428897 |
| PRO166_CB_C10 | 0.410077247 | 0.330657214 |
| PRO195_CB_C10 | 0.584951724 | 0.460851953 |
| PRO27_CB_C10 | 0.541517023 | 0.207890244 |
| PRO47_CB_C10 | 0.478425244 | 0.141428713 |
| SER142_CB_C10 | 0.594168641 | 0.433687194 |
| SER151_CB_C10 | 0.456516085 | 0.350730211 |
| SER161_CB_C10 | 0.456925512 | 0.456066341 |
| SER176_CB_C10 | 0.12835997 | 0.687905646 |
| SER20_CB_C10 | 0.243397868 | 0.725961068 |
| SER39_CB_C10 | 0.494117917 | 0.371099783 |
| SER50_CB_C10 | 0.291804246 | 0.348766827 |
| SER54_CB_C10 | 0.148579471 | 0.694575586 |
| THR159_CB_C10 | 0.421309753 | 0.485057044 |
| THR30_CB_C10 | 0.488013011 | 0.30714695 |
| THR4_CB_C10 | 0.338792668 | 0.325398131 |
| THR40_CB_C10 | 0.524969896 | 0.347383693 |
| THR68_CB_C10 | 0.24414478 | 0.469207231 |

**Table S4.** Feature-diverse tICA features and correlations – SUC transport (cont.)

| System | SUC | |
|---|---|---|
| # of features | 1002 | |
| Feature | \| tIC1 correlation (*r*) \| | \| tIC2 correlation (*r*) \| |
| **TRP180_CB_C10** | 0.401204385 | 0.586708614 |
| **TRP58_CB_C10** | 0.10708208 | 0.817353655 |
| **TYR183_CB_C10** | 0.470732084 | 0.624438036 |
| **TYR210_CB_C10** | 0.44281972 | 0.073123973 |
| **TYR48_CB_C10** | 0.57338241 | 0.170237023 |
| **TYR61_CB_C10** | 0.563761636 | 0.594181756 |
| **TYR86_CB_C10** | 0.615676596 | 0.09965519 |
| **VAL139_CB_C10** | 0.641339178 | 0.265227508 |
| **VAL145_CB_C10** | 0.234022343 | 0.397760941 |
| **VAL155_CB_C10** | 0.443961519 | 0.469122443 |
| **VAL162_CB_C10** | 0.463182441 | 0.343949844 |
| **VAL192_CB_C10** | 0.669271287 | 0.408502916 |
| **VAL23_CB_C10** | 0.401799679 | 0.602729741 |
| **ALA177_CB_C11** | 0.033324592 | 0.664572298 |
| **ALA2_CB_C11** | 0.556771589 | 0.194181227 |
| **ALA200_CB_C11** | 0.077295731 | 0.668472575 |
| **ALA51_CB_C11** | 0.503391697 | 0.385735567 |
| **ALA55_CB_C11** | 0.334317474 | 0.728824269 |
| **ALA62_CB_C11** | 0.366345055 | 0.686385634 |
| **ARG160_CB_C11** | 0.463829531 | 0.437727148 |
| **ARG33_CB_C11** | 0.505015727 | 0.362685085 |
| **ASN17_CB_C11** | 0.151667546 | 0.738699438 |
| **ASN196_CB_C11** | 0.3055156 | 0.703228241 |
| **ASN5_CB_C11** | 0.395932671 | 0.243088152 |
| **ASN76_CB_C11** | 0.191915633 | 0.837618672 |
| **ASP189_CB_C11** | 0.664280957 | 0.451127088 |
| **CYS138_CB_C11** | 0.723301982 | 0.292697587 |
| **GLN44_CB_C11** | 0.510056492 | 0.374162038 |
| **GLU131_CB_C11** | 0.676820004 | 0.335825558 |
| **GLU163_CB_C11** | 0.448000738 | 0.448044678 |
| **GLU41_CB_C11** | 0.482940185 | 0.32749574 |

**Table S4.** Feature-diverse tICA features and correlations – SUC transport (cont.)

| System | SUC | |
|---|---|---|
| # of features | 1002 | |
| Feature | l tIC1 correlation (*r*) l | l tIC2 correlation (*r*) l |
| GLU83_CB_C11 | 0.534471748 | 0.502300484 |
| GLY13_CA_C11 | 0.379943857 | 0.611571185 |
| GLY135_CA_C11 | 0.705599867 | 0.321544067 |
| GLY16_CA_C11 | 0.210164698 | 0.770090146 |
| GLY184_CA_C11 | 0.587399761 | 0.584165938 |
| GLY199_CA_C11 | 0.06127667 | 0.632522106 |
| GLY203_CA_C11 | 0.336987168 | 0.403379982 |
| GLY42_CA_C11 | 0.531213789 | 0.289248884 |
| GLY79_CA_C11 | 0.412916467 | 0.750737543 |
| ILE152_CB_C11 | 0.511583934 | 0.356461125 |
| ILE187_CB_C11 | 0.571016616 | 0.429007909 |
| ILE34_CB_C11 | 0.586353606 | 0.242413953 |
| ILE75_CB_C11 | 0.135187554 | 0.743031096 |
| LEU169_CB_C11 | 0.509973441 | 0.165812471 |
| LEU173_CB_C11 | 0.328779745 | 0.36808442 |
| LEU209_CB_C11 | 0.514858789 | 0.169069688 |
| LEU3_CB_C11 | 0.573082127 | 0.231188646 |
| LEU71_CB_C11 | 0.115539108 | 0.647175977 |
| LEU72_CB_C11 | 0.234724949 | 0.680802649 |
| LYS132_CB_C11 | 0.662511409 | 0.348284682 |
| LYS37_CB_C11 | 0.516521389 | 0.357073658 |
| LYS38_CB_C11 | 0.550798257 | 0.249917097 |
| LYS65_CB_C11 | 0.648756852 | 0.354636064 |
| MET1_CB_C11 | 0.575800258 | 0.199410484 |
| MET165_CB_C11 | 0.519479822 | 0.352287358 |
| PHE146_CB_C11 | 0.261248402 | 0.276288365 |
| PHE164_CB_C11 | 0.453640198 | 0.465413253 |
| PHE213_CB_C11 | 0.502515573 | 0.353691725 |
| PHE24_CB_C11 | 0.494299819 | 0.586061711 |
| PHE43_CB_C11 | 0.573239595 | 0.219020927 |
| PHE90_CB_C11 | 0.618176925 | 0.012369869 |

**Table S4.** Feature-diverse tICA features and correlations – SUC transport (cont.)

| System | SUC | |
|---|---|---|
| # of features | 1002 | |
| Feature | \| tIC1 correlation (*r*) \| | \| tIC2 correlation (*r*) \| |
| PRO149_CB_C11 | 0.492304359 | 0.139177983 |
| PRO166_CB_C11 | 0.404203415 | 0.349333135 |
| PRO195_CB_C11 | 0.514946235 | 0.559036113 |
| PRO27_CB_C11 | 0.561303143 | 0.212359713 |
| PRO47_CB_C11 | 0.503731929 | 0.190334148 |
| SER142_CB_C11 | 0.529590838 | 0.51909219 |
| SER151_CB_C11 | 0.485591568 | 0.377863844 |
| SER161_CB_C11 | 0.473784334 | 0.467421629 |
| SER176_CB_C11 | 0.174925181 | 0.699869236 |
| SER20_CB_C11 | 0.203800359 | 0.716038508 |
| SER39_CB_C11 | 0.516569354 | 0.373852842 |
| SER50_CB_C11 | 0.343864621 | 0.376320121 |
| SER54_CB_C11 | 0.170890069 | 0.708794569 |
| THR159_CB_C11 | 0.43566293 | 0.490810635 |
| THR30_CB_C11 | 0.525986711 | 0.310498688 |
| THR4_CB_C11 | 0.363671354 | 0.306388736 |
| THR40_CB_C11 | 0.55449006 | 0.348519262 |
| THR68_CB_C11 | 0.30561502 | 0.453401305 |
| TRP180_CB_C11 | 0.350655729 | 0.671316742 |
| TRP58_CB_C11 | 0.221286423 | 0.77172664 |
| TYR183_CB_C11 | 0.458082517 | 0.666632646 |
| TYR210_CB_C11 | 0.500180848 | 0.108277182 |
| TYR48_CB_C11 | 0.593582568 | 0.17295088 |
| TYR61_CB_C11 | 0.632854257 | 0.532885866 |
| TYR86_CB_C11 | 0.630801325 | 0.099519288 |
| VAL139_CB_C11 | 0.635866954 | 0.292072736 |
| VAL145_CB_C11 | 0.299531171 | 0.443970389 |
| VAL155_CB_C11 | 0.45925942 | 0.48509205 |
| VAL162_CB_C11 | 0.493355372 | 0.36751551 |
| VAL192_CB_C11 | 0.661480493 | 0.479114542 |
| VAL23_CB_C11 | 0.390484674 | 0.60957781 |

**Table S4.** Feature-diverse tICA features and correlations – SUC transport (cont.)

| System | SUC | |
|---|---|---|
| # of features | 1002 | |
| Feature | \| tIC1 correlation (*r*) \| | \| tIC2 correlation (*r*) \| |
| ALA177_CB_C12 | 0.091840966 | 0.634605682 |
| ALA2_CB_C12 | 0.579426882 | 0.164057845 |
| ALA200_CB_C12 | 0.044902753 | 0.643168218 |
| ALA51_CB_C12 | 0.562069931 | 0.365239833 |
| ALA55_CB_C12 | 0.354341945 | 0.660920118 |
| ALA62_CB_C12 | 0.453207321 | 0.56998906 |
| ARG160_CB_C12 | 0.500788523 | 0.391740894 |
| ARG33_CB_C12 | 0.543911282 | 0.33726707 |
| ASN17_CB_C12 | 0.090521383 | 0.674791419 |
| ASN196_CB_C12 | 0.28847837 | 0.634952013 |
| ASN5_CB_C12 | 0.417119151 | 0.193240077 |
| ASN76_CB_C12 | 0.176988481 | 0.793560365 |
| ASP189_CB_C12 | 0.715074575 | 0.357984743 |
| CYS138_CB_C12 | 0.729223196 | 0.215170884 |
| GLN44_CB_C12 | 0.557548872 | 0.348450851 |
| GLU131_CB_C12 | 0.731649255 | 0.262976169 |
| GLU163_CB_C12 | 0.476185719 | 0.41331641 |
| GLU41_CB_C12 | 0.523257091 | 0.305502292 |
| GLU83_CB_C12 | 0.591652939 | 0.454627154 |
| GLY13_CA_C12 | 0.386909105 | 0.546502868 |
| GLY135_CA_C12 | 0.737826037 | 0.240152611 |
| GLY16_CA_C12 | 0.162127252 | 0.708043168 |
| GLY184_CA_C12 | 0.621140593 | 0.489647161 |
| GLY199_CA_C12 | 0.104937033 | 0.6226049 |
| GLY203_CA_C12 | 0.384306802 | 0.437248956 |
| GLY42_CA_C12 | 0.562706253 | 0.267555008 |
| GLY79_CA_C12 | 0.455210994 | 0.706008141 |
| ILE152_CB_C12 | 0.554602772 | 0.305031108 |
| ILE187_CB_C12 | 0.626472286 | 0.352133384 |
| ILE34_CB_C12 | 0.636615525 | 0.224382231 |
| ILE75_CB_C12 | 0.147360856 | 0.67244448 |

**Table S4.** Feature-diverse tICA features and correlations – SUC transport (cont.)

| System | SUC | |
|---|---|---|
| # of features | 1002 | |
| Feature | \| tIC1 correlation (*r*) \| | \| tIC2 correlation (*r*) \| |
| LEU169_CB_C12 | 0.557026766 | 0.103941927 |
| LEU173_CB_C12 | 0.391388314 | 0.40843257 |
| LEU209_CB_C12 | 0.558625513 | 0.111333134 |
| LEU3_CB_C12 | 0.591881423 | 0.19933511 |
| LEU71_CB_C12 | 0.146789266 | 0.586869299 |
| LEU72_CB_C12 | 0.309902801 | 0.60171427 |
| LYS132_CB_C12 | 0.717242143 | 0.281334959 |
| LYS37_CB_C12 | 0.565085977 | 0.333359587 |
| LYS38_CB_C12 | 0.60187827 | 0.227106196 |
| LYS65_CB_C12 | 0.682025855 | 0.269547261 |
| MET1_CB_C12 | 0.587146701 | 0.168572737 |
| MET165_CB_C12 | 0.560952557 | 0.300416346 |
| PHE146_CB_C12 | 0.309344876 | 0.330210863 |
| PHE164_CB_C12 | 0.476134139 | 0.428549011 |
| PHE213_CB_C12 | 0.54503495 | 0.301716827 |
| PHE24_CB_C12 | 0.563227926 | 0.550018622 |
| PHE43_CB_C12 | 0.609842996 | 0.19466071 |
| PHE90_CB_C12 | 0.668650343 | 0.012822456 |
| PRO149_CB_C12 | 0.539098874 | 0.063722645 |
| PRO166_CB_C12 | 0.45657139 | 0.303787779 |
| PRO195_CB_C12 | 0.509598967 | 0.510868589 |
| PRO27_CB_C12 | 0.58373804 | 0.206162991 |
| PRO47_CB_C12 | 0.533875279 | 0.202820347 |
| SER142_CB_C12 | 0.487371572 | 0.498191232 |
| SER151_CB_C12 | 0.508116823 | 0.336351904 |
| SER161_CB_C12 | 0.505223268 | 0.426342353 |
| SER176_CB_C12 | 0.254028711 | 0.626729597 |
| SER20_CB_C12 | 0.269968506 | 0.632789484 |
| SER39_CB_C12 | 0.563023166 | 0.349645203 |
| SER50_CB_C12 | 0.4328661 | 0.36408068 |
| SER54_CB_C12 | 0.263002206 | 0.645625562 |

**Table S4.** Feature-diverse tICA features and correlations – SUC transport (cont.)

| System | SUC | |
|---|---|---|
| # of features | 1002 | |
| Feature | \| tIC1 correlation (*r*) \| | \| tIC2 correlation (*r*) \| |
| THR159_CB_C12 | 0.467388856 | 0.453441945 |
| THR30_CB_C12 | 0.566076157 | 0.289259025 |
| THR4_CB_C12 | 0.397041571 | 0.255481711 |
| THR40_CB_C12 | 0.595340621 | 0.327776769 |
| THR68_CB_C12 | 0.365180983 | 0.388658201 |
| TRP180_CB_C12 | 0.336012513 | 0.589195336 |
| TRP58_CB_C12 | 0.310661896 | 0.666240661 |
| TYR183_CB_C12 | 0.472721932 | 0.586603625 |
| TYR210_CB_C12 | 0.553369503 | 0.049085732 |
| TYR48_CB_C12 | 0.640810972 | 0.135581542 |
| TYR61_CB_C12 | 0.672689641 | 0.395641465 |
| TYR86_CB_C12 | 0.675122478 | 0.076111273 |
| VAL139_CB_C12 | 0.693512747 | 0.266580596 |
| VAL145_CB_C12 | 0.366985535 | 0.45843449 |
| VAL155_CB_C12 | 0.494664961 | 0.445903922 |
| VAL162_CB_C12 | 0.53659866 | 0.318601207 |
| VAL192_CB_C12 | 0.694227812 | 0.379071645 |
| VAL23_CB_C12 | 0.437858977 | 0.548818562 |
| ALA177_CB_O5 | 0.400637885 | 0.640808241 |
| ALA2_CB_O5 | 0.727197726 | 0.066457004 |
| ALA200_CB_O5 | 0.331653131 | 0.73832693 |
| ALA51_CB_O5 | 0.573553426 | 0.383842112 |
| ALA55_CB_O5 | 0.170239977 | 0.619033785 |
| ALA62_CB_O5 | 0.772247635 | 0.224256321 |
| ARG160_CB_O5 | 0.717838497 | 0.20047707 |
| ARG33_CB_O5 | 0.642743447 | 0.340649215 |
| ASN17_CB_O5 | 0.613377166 | 0.442873842 |
| ASN196_CB_O5 | 0.016095153 | 0.755802684 |
| ASN5_CB_O5 | 0.750700453 | 0.120464503 |
| ASN76_CB_O5 | 0.371081488 | 0.623347498 |
| ASP189_CB_O5 | 0.814093159 | 0.238165392 |

**Table S4.** Feature-diverse tICA features and correlations – SUC transport (cont.)

| System | SUC | |
|---|---|---|
| # of features | 1002 | |
| Feature | I tIC1 correlation (*r*) I | I tIC2 correlation (*r*) I |
| CYS138_CB_O5 | 0.688959632 | 0.395941164 |
| GLN44_CB_O5 | 0.567122565 | 0.403294773 |
| GLU131_CB_O5 | 0.832169875 | 0.181700174 |
| GLU163_CB_O5 | 0.705127623 | 0.30267888 |
| GLU41_CB_O5 | 0.676983121 | 0.314768525 |
| GLU83_CB_O5 | 0.624068561 | 0.316723052 |
| GLY13_CA_O5 | 0.784468304 | 0.240578757 |
| GLY135_CA_O5 | 0.781911545 | 0.277135288 |
| GLY16_CA_O5 | 0.592152143 | 0.550821975 |
| GLY184_CA_O5 | 0.742775269 | 0.414435974 |
| GLY199_CA_O5 | 0.434713149 | 0.628294342 |
| GLY203_CA_O5 | 0.615486189 | 0.625191627 |
| GLY42_CA_O5 | 0.680570289 | 0.310132225 |
| GLY79_CA_O5 | 0.293381981 | 0.657122694 |
| ILE152_CB_O5 | 0.789533904 | 0.083275622 |
| ILE187_CB_O5 | 0.852945409 | 0.184687455 |
| ILE34_CB_O5 | 0.68977813 | 0.298347283 |
| ILE75_CB_O5 | 0.441621986 | 0.550973329 |
| LEU169_CB_O5 | 0.733508164 | 0.134612434 |
| LEU173_CB_O5 | 0.581566025 | 0.578496813 |
| LEU209_CB_O5 | 0.807456783 | 0.204552152 |
| LEU3_CB_O5 | 0.765147081 | 0.079672007 |
| LEU71_CB_O5 | 0.655953714 | 0.255912249 |
| LEU72_CB_O5 | 0.766728471 | 0.204191824 |
| LYS132_CB_O5 | 0.761710885 | 0.275815111 |
| LYS37_CB_O5 | 0.66872634 | 0.354165876 |
| LYS38_CB_O5 | 0.695188244 | 0.301790804 |
| LYS65_CB_O5 | 0.816632117 | 0.118571783 |
| MET1_CB_O5 | 0.662735138 | 0.061582505 |
| MET165_CB_O5 | 0.754563513 | 0.085401401 |
| PHE146_CB_O5 | 0.646286219 | 0.59619251 |

**Table S4.** Feature-diverse tICA features and correlations – SUC transport (cont.)

| System | SUC | |
|---|---|---|
| # of features | 1002 | |
| Feature | \| tIC1 correlation (*r*) \| | \| tIC2 correlation (*r*) \| |
| PHE164_CB_O5 | 0.670815269 | 0.348621514 |
| PHE213_CB_O5 | 0.791882337 | 0.057324521 |
| PHE24_CB_O5 | 0.578205899 | 0.486469319 |
| PHE43_CB_O5 | 0.641697942 | 0.329605529 |
| PHE90_CB_O5 | 0.719609843 | 0.179606379 |
| PRO149_CB_O5 | 0.765216247 | 0.307960339 |
| PRO166_CB_O5 | 0.708418051 | 0.122198912 |
| PRO195_CB_O5 | 0.009399076 | 0.672876208 |
| PRO27_CB_O5 | 0.613664232 | 0.217220016 |
| PRO47_CB_O5 | 0.714714313 | 0.064950672 |
| SER142_CB_O5 | 0.026180526 | 0.63138814 |
| SER151_CB_O5 | 0.693243518 | 0.234409359 |
| SER161_CB_O5 | 0.719676616 | 0.282486805 |
| SER176_CB_O5 | 0.440640008 | 0.630189822 |
| SER20_CB_O5 | 0.138221116 | 0.708160091 |
| SER39_CB_O5 | 0.666580028 | 0.377235285 |
| SER50_CB_O5 | 0.598779876 | 0.386647242 |
| SER54_CB_O5 | 0.249802009 | 0.556160341 |
| THR159_CB_O5 | 0.657797138 | 0.33713214 |
| THR30_CB_O5 | 0.597970684 | 0.301410582 |
| THR4_CB_O5 | 0.693152539 | 0.117191414 |
| THR40_CB_O5 | 0.668948648 | 0.386705912 |
| THR68_CB_O5 | 0.763402219 | 0.16889999 |
| TRP180_CB_O5 | 0.478146964 | 0.567237467 |
| TRP58_CB_O5 | 0.688071339 | 0.405205371 |
| TYR183_CB_O5 | 0.766654585 | 0.399942435 |
| TYR210_CB_O5 | 0.799553854 | 0.2614379 |
| TYR48_CB_O5 | 0.709435232 | 0.030451556 |
| TYR61_CB_O5 | 0.796496971 | 0.274221382 |
| TYR86_CB_O5 | 0.713674852 | 0.06056232 |
| VAL139_CB_O5 | 0.549971596 | 0.476736458 |

**Table S4.** Feature-diverse tICA features and correlations – SUC transport (cont.)

| System | SUC | |
|---|---|---|
| # of features | 1002 | |
| Feature | \| tIC1 correlation (*r*) \| | \| tIC2 correlation (*r*) \| |
| VAL145_CB_O5 | 0.541013918 | 0.561613441 |
| VAL155_CB_O5 | 0.646707754 | 0.342760317 |
| VAL162_CB_O5 | 0.7808552 | 0.095244446 |
| VAL192_CB_O5 | 0.752235137 | 0.275243088 |
| VAL23_CB_O5 | 0.535712796 | 0.609260082 |
| ALA177_CB_O6 | 0.395684985 | 0.659674764 |
| ALA2_CB_O6 | 0.661510413 | 0.082914582 |
| ALA200_CB_O6 | 0.359471328 | 0.713826939 |
| ALA51_CB_O6 | 0.508546647 | 0.44804395 |
| ALA55_CB_O6 | 0.029368758 | 0.656477774 |
| ALA62_CB_O6 | 0.792417372 | 0.257295386 |
| ARG160_CB_O6 | 0.700328673 | 0.330852034 |
| ARG33_CB_O6 | 0.60366609 | 0.358333797 |
| ASN17_CB_O6 | 0.730938831 | 0.407558295 |
| ASN196_CB_O6 | 0.159475482 | 0.712760728 |
| ASN5_CB_O6 | 0.665096155 | 0.147637481 |
| ASN76_CB_O6 | 0.487903315 | 0.541684537 |
| ASP189_CB_O6 | 0.698161753 | 0.391667267 |
| CYS138_CB_O6 | 0.543633686 | 0.502438955 |
| GLN44_CB_O6 | 0.634541988 | 0.345615603 |
| GLU131_CB_O6 | 0.790518098 | 0.285161329 |
| GLU163_CB_O6 | 0.632727358 | 0.416804308 |
| GLU41_CB_O6 | 0.616656454 | 0.327731477 |
| GLU83_CB_O6 | 0.509801085 | 0.483060348 |
| GLY13_CA_O6 | 0.80007552 | 0.322161923 |
| GLY135_CA_O6 | 0.697612382 | 0.425925041 |
| GLY16_CA_O6 | 0.614642608 | 0.520423624 |
| GLY184_CA_O6 | 0.538852994 | 0.576197344 |
| GLY199_CA_O6 | 0.481317342 | 0.657422277 |
| GLY203_CA_O6 | 0.649522503 | 0.545115044 |
| GLY42_CA_O6 | 0.661922996 | 0.316669269 |

**Table S4.** Feature-diverse tICA features and correlations – SUC transport (cont.)

| System | SUC | |
|---|---|---|
| # of features | 1002 | |
| Feature | l tIC1 correlation (*r*) l | l tIC2 correlation (*r*) l |
| GLY79_CA_O6 | 0.066346353 | 0.683102456 |
| ILE152_CB_O6 | 0.687074952 | 0.294930225 |
| ILE187_CB_O6 | 0.75878019 | 0.282064743 |
| ILE34_CB_O6 | 0.660844846 | 0.256762345 |
| ILE75_CB_O6 | 0.592585195 | 0.456402376 |
| LEU169_CB_O6 | 0.682022643 | 0.048440793 |
| LEU173_CB_O6 | 0.624032624 | 0.535470336 |
| LEU209_CB_O6 | 0.786605786 | 0.005378513 |
| LEU3_CB_O6 | 0.741292914 | 0.109694388 |
| LEU71_CB_O6 | 0.696763164 | 0.265902864 |
| LEU72_CB_O6 | 0.733117773 | 0.177683713 |
| LYS132_CB_O6 | 0.649277639 | 0.419006688 |
| LYS37_CB_O6 | 0.614492145 | 0.364058283 |
| LYS38_CB_O6 | 0.634383382 | 0.280344731 |
| LYS65_CB_O6 | 0.822549456 | 0.181355227 |
| MET1_CB_O6 | 0.634854902 | 0.083390737 |
| MET165_CB_O6 | 0.664642687 | 0.269343848 |
| PHE146_CB_O6 | 0.681732031 | 0.55531075 |
| PHE164_CB_O6 | 0.585164779 | 0.455225822 |
| PHE213_CB_O6 | 0.744215432 | 0.235987567 |
| PHE24_CB_O6 | 0.369095772 | 0.556671172 |
| PHE43_CB_O6 | 0.690895553 | 0.217467298 |
| PHE90_CB_O6 | 0.700129421 | 0.029045642 |
| PRO149_CB_O6 | 0.733027251 | 0.097306474 |
| PRO166_CB_O6 | 0.605776034 | 0.284083453 |
| PRO195_CB_O6 | 0.233089066 | 0.729641621 |
| PRO27_CB_O6 | 0.642676296 | 0.137455845 |
| PRO47_CB_O6 | 0.689526681 | 0.022515053 |
| SER142_CB_O6 | 0.227923249 | 0.670402496 |
| SER151_CB_O6 | 0.597548751 | 0.346674166 |
| SER161_CB_O6 | 0.667102999 | 0.407633955 |

**Table S4.** Feature-diverse tICA features and correlations – SUC transport (cont.)

| System | SUC | |
|---|---|---|
| # of features | 1002 | |
| Feature | \| tIC1 correlation (*r*) \| | \| tIC2 correlation (*r*) \| |
| SER176_CB_O6 | 0.363198364 | 0.72068671 |
| SER20_CB_O6 | 0.169153748 | 0.648485407 |
| SER39_CB_O6 | 0.600154557 | 0.365511167 |
| SER50_CB_O6 | 0.552792389 | 0.471477252 |
| SER54_CB_O6 | 0.263896619 | 0.661221017 |
| THR159_CB_O6 | 0.636261945 | 0.44547174 |
| THR30_CB_O6 | 0.612665212 | 0.280040878 |
| THR4_CB_O6 | 0.656983566 | 0.158439188 |
| THR40_CB_O6 | 0.668168707 | 0.335999572 |
| THR68_CB_O6 | 0.697283138 | 0.195171086 |
| TRP180_CB_O6 | 0.267295086 | 0.589903335 |
| TRP58_CB_O6 | 0.657046525 | 0.308403527 |
| TYR183_CB_O6 | 0.622664368 | 0.489136627 |
| TYR210_CB_O6 | 0.764687628 | 0.031745767 |
| TYR48_CB_O6 | 0.661902757 | 0.226596217 |
| TYR61_CB_O6 | 0.744001703 | 0.311898555 |
| TYR86_CB_O6 | 0.680827526 | 0.107109135 |
| VAL139_CB_O6 | 0.329302892 | 0.682278054 |
| VAL145_CB_O6 | 0.56235283 | 0.590841145 |
| VAL155_CB_O6 | 0.577274952 | 0.45971949 |
| VAL162_CB_O6 | 0.700751382 | 0.278144799 |
| VAL192_CB_O6 | 0.633308642 | 0.465097994 |
| VAL23_CB_O6 | 0.414033983 | 0.651182518 |
| ALA177_CB_O9 | 0.157681229 | 0.776672799 |
| ALA2_CB_O9 | 0.598135519 | 0.182373784 |
| ALA200_CB_O9 | 0.062771072 | 0.775478992 |
| ALA51_CB_O9 | 0.518671442 | 0.448015703 |
| ALA55_CB_O9 | 0.24784135 | 0.809640394 |
| ALA62_CB_O9 | 0.517615794 | 0.630556293 |
| ARG160_CB_O9 | 0.516215734 | 0.439377557 |
| ARG33_CB_O9 | 0.518594654 | 0.383046957 |

**Table S4.** Feature-diverse tICA features and correlations – SUC transport (cont.)

| System | SUC | |
|---|---|---|
| # of features | 1002 | |
| Feature | \| tIC1 correlation ($r$) \| | \| tIC2 correlation ($r$) \| |
| ASN17_CB_O9 | 0.344924948 | 0.718024608 |
| ASN196_CB_O9 | 0.183465753 | 0.845140808 |
| ASN5_CB_O9 | 0.445769965 | 0.237043506 |
| ASN76_CB_O9 | 0.014423787 | 0.876006084 |
| ASP189_CB_O9 | 0.663936294 | 0.499881218 |
| CYS138_CB_O9 | 0.704534283 | 0.363674161 |
| GLN44_CB_O9 | 0.531245562 | 0.399037058 |
| GLU131_CB_O9 | 0.693899947 | 0.355314314 |
| GLU163_CB_O9 | 0.473516039 | 0.458794778 |
| GLU41_CB_O9 | 0.504247091 | 0.3493407 |
| GLU83_CB_O9 | 0.520501272 | 0.531709651 |
| GLY13_CA_O9 | 0.518884643 | 0.600904766 |
| GLY135_CA_O9 | 0.691224611 | 0.374974927 |
| GLY16_CA_O9 | 0.348755233 | 0.773267748 |
| GLY184_CA_O9 | 0.572064753 | 0.659951909 |
| GLY199_CA_O9 | 0.243143261 | 0.75056883 |
| GLY203_CA_O9 | 0.459469133 | 0.454963713 |
| GLY42_CA_O9 | 0.553065182 | 0.318055396 |
| GLY79_CA_O9 | 0.332818764 | 0.798600874 |
| ILE152_CB_O9 | 0.550717981 | 0.37896868 |
| ILE187_CB_O9 | 0.604110487 | 0.432818564 |
| ILE34_CB_O9 | 0.590165292 | 0.270557474 |
| ILE75_CB_O9 | 0.040521266 | 0.763925925 |
| LEU169_CB_O9 | 0.551441218 | 0.175819808 |
| LEU173_CB_O9 | 0.414617113 | 0.412894297 |
| LEU209_CB_O9 | 0.594347375 | 0.163779992 |
| LEU3_CB_O9 | 0.625650136 | 0.217403642 |
| LEU71_CB_O9 | 0.286408046 | 0.624926131 |
| LEU72_CB_O9 | 0.426465872 | 0.622725674 |
| LYS132_CB_O9 | 0.642822521 | 0.40093676 |
| LYS37_CB_O9 | 0.526711928 | 0.37926119 |

**Table S4.** Feature-diverse tICA features and correlations – SUC transport (cont.)

| System | SUC | |
|---|---|---|
| # of features | 1002 | |
| Feature | \| tIC1 correlation (*r*) \| | \| tIC2 correlation (*r*) \| |
| LYS38_CB_O9 | 0.557069724 | 0.280082087 |
| LYS65_CB_O9 | 0.703122142 | 0.341363268 |
| MET1_CB_O9 | 0.603720563 | 0.180926169 |
| MET165_CB_O9 | 0.556782424 | 0.366217067 |
| PHE146_CB_O9 | 0.421459196 | 0.36692207 |
| PHE164_CB_O9 | 0.483719391 | 0.491650014 |
| PHE213_CB_O9 | 0.55850385 | 0.356200378 |
| PHE24_CB_O9 | 0.448517537 | 0.606045811 |
| PHE43_CB_O9 | 0.605359053 | 0.24945636 |
| PHE90_CB_O9 | 0.625437394 | 0.042123867 |
| PRO149_CB_O9 | 0.545531763 | 0.139090134 |
| PRO166_CB_O9 | 0.428920224 | 0.36591115 |
| PRO195_CB_O9 | 0.293295026 | 0.761888776 |
| PRO27_CB_O9 | 0.58283617 | 0.224610643 |
| PRO47_CB_O9 | 0.533915035 | 0.188537023 |
| SER142_CB_O9 | 0.312804626 | 0.715697784 |
| SER151_CB_O9 | 0.533947202 | 0.400112157 |
| SER161_CB_O9 | 0.510734601 | 0.478330945 |
| SER176_CB_O9 | 0.209015269 | 0.766403721 |
| SER20_CB_O9 | 0.101943619 | 0.785229413 |
| SER39_CB_O9 | 0.529870202 | 0.395255214 |
| SER50_CB_O9 | 0.388715986 | 0.44285672 |
| SER54_CB_O9 | 0.129939043 | 0.755306433 |
| THR159_CB_O9 | 0.473530268 | 0.503367057 |
| THR30_CB_O9 | 0.545385801 | 0.330251047 |
| THR4_CB_O9 | 0.436818612 | 0.295474139 |
| THR40_CB_O9 | 0.568168272 | 0.371419339 |
| THR68_CB_O9 | 0.408311486 | 0.433472122 |
| TRP180_CB_O9 | 0.365391254 | 0.76074082 |
| TRP58_CB_O9 | 0.413704845 | 0.720816266 |
| TYR183_CB_O9 | 0.520546976 | 0.699763857 |

**Table S4.** Feature-diverse tICA features and correlations – SUC transport (cont.)

| System | SUC | |
|---|---|---|
| # of features | 1002 | |
| Feature | \| tIC1 correlation (*r*) \| | \| tIC2 correlation (*r*) \| |
| TYR210_CB_O9 | 0.572130202 | 0.106968282 |
| TYR48_CB_O9 | 0.607840823 | 0.174531454 |
| TYR61_CB_O9 | 0.693653085 | 0.51469256 |
| TYR86_CB_O9 | 0.636003541 | 0.081641476 |
| VAL139_CB_O9 | 0.598859746 | 0.426079439 |
| VAL145_CB_O9 | 0.391453507 | 0.553654346 |
| VAL155_CB_O9 | 0.480089081 | 0.501376884 |
| VAL162_CB_O9 | 0.540005256 | 0.377061396 |
| VAL192_CB_O9 | 0.650251741 | 0.544841798 |
| VAL23_CB_O9 | 0.379888631 | 0.654507068 |

**Table S5.** Feature-diverse tICA features and correlations – GLC transport

| System | GLC | |
|---|---|---|
| # of features | 498 | |
| Feature | l tIC1 correlation (*r*) l | l tIC2 correlation (*r*) l |
| theta_C3_xy | 0.001052478 | 0.002427539 |
| theta_C3_xz | 0.001464052 | 0.001249731 |
| theta_C3_yz | 0.000996129 | 0.004713568 |
| theta_C4_xy | 0.002143311 | 9.94827E-05 |
| theta_C4_xz | 0.002678207 | 0.000625455 |
| theta_C4_yz | 0.000426365 | 0.002094419 |
| theta_O3_xy | 0.005793633 | 0.001369311 |
| theta_O3_xz | 0.007110356 | 0.002671049 |
| theta_O3_yz | 0.000249144 | 0.005023463 |
| theta_O4_xy | 0.003259637 | 0.00118086 |
| theta_O4_xz | 0.003701709 | 0.001785342 |
| theta_O4_yz | 0.001521904 | 0.001095996 |
| GLY203_CA_ALA177_CB | 0.118569167 | 0.065364732 |
| LEU173_CB_ALA177_CB | 0.01654979 | 0.021971227 |
| SER176_CB_ALA177_CB | 0.070462909 | 0.023447591 |
| MET1_CB_ALA2_CB | 0.255022811 | 0.086297402 |
| ALA177_CB_ALA200_CB | 0.081137852 | 0.036535181 |
| GLY199_CA_ALA200_CB | 0.012739735 | 0.004726496 |
| GLY203_CA_ALA200_CB | 0.027869783 | 0.052601268 |
| PRO47_CB_ALA51_CB | 0.000679023 | 0.011437987 |
| SER50_CB_ALA51_CB | 0.056119786 | 0.063441169 |
| TYR48_CB_ALA51_CB | 0.07948126 | 0.004361781 |
| ALA51_CB_ALA55_CB | 0.001542572 | 0.008191417 |
| GLY79_CA_ALA55_CB | 0.01735778 | 0.074534705 |
| SER54_CB_ALA55_CB | 0.052917371 | 0.015898339 |
| LEU72_CB_ALA62_CB | 0.354257995 | 0.137715584 |
| TRP58_CB_ALA62_CB | 0.120653081 | 0.041190982 |
| TYR61_CB_ALA62_CB | 0.057956331 | 0.004560453 |
| GLU41_CB_ARG33_CB | 0.7501723 | 0.010260202 |
| ILE75_CB_ASN17_CB | 0.189993253 | 0.017490625 |
| SER20_CB_ASN17_CB | 0.401673331 | 0.104824599 |

**Table S5.** Feature-diverse tICA features and correlations – GLC transport (cont)

| System | GLC | |
|---|---|---|
| # of features | 498 | |
| Feature | l tIC1 correlation (*r*) l | l tIC2 correlation (*r*) l |
| ALA177_CB_ASN196_CB | 0.036410393 | 0.059058268 |
| ALA200_CB_ASN196_CB | 0.023961074 | 0.000912098 |
| GLY199_CA_ASN196_CB | 0.04339627 | 0.05134446 |
| TRP180_CB_ASN196_CB | 0.005699086 | 0.009233579 |
| ALA2_CB_ASN5_CB | 0.252071622 | 0.139397163 |
| THR4_CB_ASN5_CB | 0.209162772 | 0.087536782 |
| ALA55_CB_ASN76_CB | 0.0074455 | 0.01302181 |
| GLY79_CA_ASN76_CB | 0.033776912 | 0.022635177 |
| GLY184_CA_ASP189_CB | 0.044549086 | 0.000457399 |
| ILE187_CB_ASP189_CB | 0.03887334 | 0.000123747 |
| TRP58_CB_CYS138_CB | 0.336880766 | 0.129478613 |
| PHE43_CB_GLN44_CB | 0.206953418 | 0.001364964 |
| GLY135_CA_GLU131_CB | 0.074182132 | 0.095281384 |
| LYS65_CB_GLU131_CB | 0.207884812 | 0.030961262 |
| SER161_CB_GLU163_CB | 0.361040422 | 0.14371621 |
| SER39_CB_GLU41_CB | 0.398519774 | 0.032798896 |
| THR40_CB_GLU41_CB | 0.644568405 | 0.020224753 |
| ALA51_CB_GLU83_CB | 0.02553114 | 0.007473088 |
| TYR48_CB_GLU83_CB | 0.016391632 | 0.00759257 |
| TYR86_CB_GLU83_CB | 0.028756601 | 0.014949948 |
| GLY16_CA_GLY13_CA | 0.255083964 | 0.067742563 |
| CYS138_CB_GLY135_CA | 0.149511106 | 0.177947212 |
| VAL139_CB_GLY135_CA | 0.092773992 | 0.114930333 |
| ILE75_CB_GLY16_CA | 0.892348952 | 0.198474474 |
| SER20_CB_GLY16_CA | 0.116692584 | 0.03444149 |
| TRP180_CB_GLY184_CA | 0.048413642 | 0.024971341 |
| TYR183_CB_GLY184_CA | 0.042809233 | 0.006194336 |
| ALA177_CB_GLY199_CA | 0.073463668 | 0.044986126 |
| GLY203_CA_GLY199_CA | 0.039744653 | 0.070218523 |
| LEU173_CB_GLY199_CA | 0.05261034 | 0.004052121 |
| PHE146_CB_GLY199_CA | 0.089264059 | 0.207593128 |

**Table S5.** Feature-diverse tICA features and correlations – GLC transport (cont)

| System | GLC | |
|---|---|---|
| # of features | 498 | |
| **Feature** | **\| tIC1 correlation (*r*) \|** | **\| tIC2 correlation (*r*) \|** |
| SER142_CB_GLY199_CA | 0.016336283 | 0.315529866 |
| LEU173_CB_GLY203_CA | 0.269213293 | 0.218647531 |
| GLU41_CB_GLY42_CA | 0.345210863 | 0.022801062 |
| PHE24_CB_GLY79_CA | 0.409622893 | 0.260429782 |
| GLN44_CB_ILE152_CB | 0.444407162 | 0.332856264 |
| MET165_CB_ILE152_CB | 0.270586665 | 0.068830931 |
| PHE164_CB_ILE152_CB | 0.422546857 | 0.076364237 |
| PHE43_CB_ILE152_CB | 0.677275437 | 0.079204184 |
| SER151_CB_ILE152_CB | 0.02841677 | 0.011271247 |
| VAL155_CB_ILE152_CB | 0.050667863 | 0.037065943 |
| LEU3_CB_ILE187_CB | 0.319620529 | 0.203251448 |
| THR68_CB_ILE187_CB | 0.799429001 | 0.154351213 |
| ARG33_CB_ILE34_CB | 0.669432339 | 0.010723572 |
| GLU41_CB_ILE34_CB | 0.725824733 | 0.01431892 |
| SER20_CB_ILE75_CB | 0.421556136 | 0.005374901 |
| PHE43_CB_LEU169_CB | 0.762887631 | 0.093378661 |
| PRO166_CB_LEU169_CB | 0.604199957 | 0.03593678 |
| PRO27_CB_LEU169_CB | 0.58207293 | 0.188384294 |
| PRO47_CB_LEU169_CB | 0.610376844 | 0.245054854 |
| PHE146_CB_LEU173_CB | 0.127507384 | 0.255616558 |
| PRO149_CB_LEU173_CB | 0.086677381 | 0.26654871 |
| PHE213_CB_LEU209_CB | 0.058508373 | 0.062204555 |
| ALA2_CB_LEU3_CB | 0.188591496 | 0.071300559 |
| ASN17_CB_LEU71_CB | 0.039013276 | 0.159411003 |
| GLY13_CA_LEU71_CB | 0.902963764 | 0.205436832 |
| GLY16_CA_LEU71_CB | 0.895043334 | 0.186586979 |
| ILE75_CB_LEU71_CB | 0.349247362 | 0.043900236 |
| LEU72_CB_LEU71_CB | 0.682951037 | 0.178074366 |
| ILE75_CB_LEU72_CB | 0.371079896 | 0.167200081 |
| TRP58_CB_LEU72_CB | 0.339109103 | 0.178139507 |
| GLU131_CB_LYS132_CB | 0.007090973 | 0.031086975 |

**Table S5.** Feature-diverse tICA features and correlations – GLC transport (cont)

| System | GLC | |
|---|---|---|
| # of features | 498 | |
| Feature | ǀ tIC1 correlation (*r*) ǀ | ǀ tIC2 correlation (*r*) ǀ |
| GLY135_CA_LYS132_CB | 0.143355594 | 0.023166509 |
| ARG33_CB_LYS37_CB | 0.555244051 | 0.050493234 |
| GLU41_CB_LYS37_CB | 0.096836112 | 0.021918431 |
| ILE34_CB_LYS37_CB | 0.013784033 | 0.05883934 |
| SER39_CB_LYS37_CB | 0.360001045 | 0.023943666 |
| LYS37_CB_LYS38_CB | 0.249244944 | 0.006816309 |
| SER39_CB_LYS38_CB | 0.036807044 | 0.005305367 |
| ALA2_CB_LYS65_CB | 0.224071136 | 0.089910999 |
| ALA62_CB_LYS65_CB | 0.277590542 | 0.013438761 |
| LEU72_CB_LYS65_CB | 0.033086997 | 0.100993859 |
| GLY13_CA_MET1_CB | 0.564304725 | 0.011480152 |
| LEU71_CB_MET1_CB | 0.458127812 | 0.046921471 |
| GLY42_CA_MET165_CB | 0.717462292 | 0.10446522 |
| PHE164_CB_MET165_CB | 0.122732563 | 0.080848478 |
| VAL162_CB_MET165_CB | 0.439376465 | 0.089464816 |
| PRO149_CB_PHE146_CB | 0.003562505 | 0.165428141 |
| GLU163_CB_PHE164_CB | 0.161443975 | 0.013494106 |
| GLY42_CA_PHE164_CB | 0.275613572 | 0.405552266 |
| PHE43_CB_PHE164_CB | 0.244395071 | 0.240306887 |
| SER161_CB_PHE164_CB | 0.210157204 | 0.06325331 |
| VAL162_CB_PHE164_CB | 0.150925833 | 0.055091313 |
| ARG160_CB_PHE213_CB | 0.133070874 | 0.179301535 |
| SER161_CB_PHE213_CB | 0.172280079 | 0.179898524 |
| VAL162_CB_PHE213_CB | 0.211998748 | 0.021069695 |
| GLY42_CA_PHE43_CB | 0.667193057 | 0.04932232 |
| THR40_CB_PHE43_CB | 0.822930946 | 0.007871825 |
| ILE152_CB_PRO149_CB | 0.008985319 | 0.02404728 |
| PHE43_CB_PRO149_CB | 0.659861807 | 0.080180728 |
| PRO47_CB_PRO149_CB | 0.397872741 | 0.083044604 |
| GLY42_CA_PRO166_CB | 0.831817691 | 0.063166415 |
| MET165_CB_PRO166_CB | 0.041959276 | 0.147452986 |

**Table S5.** Feature-diverse tICA features and correlations – GLC transport (cont)

| System | GLC | |
|---|---|---|
| # of features | 498 | |
| Feature | l tIC1 correlation (*r*) l | l tIC2 correlation (*r*) l |
| THR30_CB_PRO166_CB | 0.621051642 | 0.198362773 |
| CYS138_CB_PRO195_CB | 0.224951633 | 0.436573086 |
| SER142_CB_PRO195_CB | 0.016596445 | 0.231412397 |
| VAL139_CB_PRO195_CB | 0.217544421 | 0.056191625 |
| PHE43_CB_PRO27_CB | 0.270922264 | 0.073821959 |
| THR30_CB_PRO27_CB | 0.110866979 | 0.051035174 |
| PHE43_CB_PRO47_CB | 0.68507126 | 0.016897219 |
| VAL145_CB_SER142_CB | 0.231427532 | 0.040050168 |
| PHE43_CB_SER151_CB | 0.531339586 | 0.056790598 |
| ARG160_CB_SER161_CB | 0.032737445 | 0.103778982 |
| LEU173_CB_SER176_CB | 0.242870294 | 0.030385565 |
| SER20_CB_SER176_CB | 0.502463342 | 0.056316651 |
| VAL23_CB_SER176_CB | 0.203984742 | 0.033770675 |
| VAL23_CB_SER20_CB | 0.375178059 | 0.102035742 |
| PRO47_CB_SER50_CB | 0.058477237 | 0.063124613 |
| ALA51_CB_SER54_CB | 0.127216553 | 0.023054339 |
| SER50_CB_SER54_CB | 0.154557223 | 0.015030266 |
| VAL145_CB_SER54_CB | 0.138762015 | 0.27703972 |
| GLU163_CB_THR159_CB | 0.469915812 | 0.189150838 |
| PHE164_CB_THR159_CB | 0.030922115 | 0.223223646 |
| SER161_CB_THR159_CB | 0.207315894 | 0.070825659 |
| ARG33_CB_THR30_CB | 0.746609437 | 0.007404086 |
| GLU41_CB_THR30_CB | 0.739447559 | 0.02423976 |
| GLY42_CA_THR30_CB | 0.865420685 | 0.007264994 |
| ALA2_CB_THR4_CB | 0.359909499 | 0.035851048 |
| LEU3_CB_THR4_CB | 0.043770359 | 0.014715117 |
| LEU71_CB_THR68_CB | 0.545931697 | 0.113025161 |
| LYS65_CB_THR68_CB | 0.580302799 | 0.163712194 |
| GLY16_CA_TRP180_CB | 0.051942628 | 0.017648765 |
| ALA55_CB_TRP58_CB | 0.036628749 | 0.085160563 |
| ASN76_CB_TRP58_CB | 0.115976218 | 0.063371461 |

**Table S5.** Feature-diverse tICA features and correlations – GLC transport (cont)

| System | GLC | |
|---|---|---|
| # of features | 498 | |
| **Feature** | **I tIC1 correlation (*r*) I** | **I tIC2 correlation (*r*) I** |
| GLY13_CA_TYR183_CB | 0.11722316 | 0.071636728 |
| GLY16_CA_TYR183_CB | 0.038822577 | 0.076205403 |
| TRP180_CB_TYR183_CB | 0.024094994 | 0.036822969 |
| LEU209_CB_TYR210_CB | 0.073483274 | 0.039574823 |
| VAL162_CB_TYR210_CB | 0.167787165 | 0.102226216 |
| PHE90_CB_TYR48_CB | 0.139649541 | 0.045985514 |
| CYS138_CB_TYR61_CB | 0.450764676 | 0.325732428 |
| TRP58_CB_TYR61_CB | 0.244359717 | 0.017518117 |
| PHE90_CB_TYR86_CB | 0.258718212 | 0.021468918 |
| PRO27_CB_TYR86_CB | 0.146760035 | 0.094957695 |
| TYR48_CB_TYR86_CB | 0.201197424 | 0.042683234 |
| CYS138_CB_VAL139_CB | 0.083381694 | 0.161079384 |
| SER142_CB_VAL139_CB | 0.093447899 | 0.162360163 |
| ALA51_CB_VAL145_CB | 0.041932755 | 0.196285137 |
| PRO47_CB_VAL145_CB | 0.111679678 | 0.000571634 |
| SER50_CB_VAL145_CB | 0.360140822 | 0.051899115 |
| MET165_CB_VAL155_CB | 0.240374705 | 0.098610296 |
| PHE164_CB_VAL155_CB | 0.480554047 | 0.219500481 |
| SER161_CB_VAL162_CB | 0.06952473 | 0.085657525 |
| ASP189_CB_VAL192_CB | 0.178174294 | 0.036001388 |
| GLY184_CA_VAL192_CB | 0.119372566 | 0.029627109 |
| TRP180_CB_VAL192_CB | 0.225431933 | 0.01173529 |
| TYR183_CB_VAL192_CB | 0.253546355 | 0.030171378 |
| ALA177_CB_C3 | 0.254530489 | 0.769865189 |
| ALA2_CB_C3 | 0.298298315 | 0.298902523 |
| ALA200_CB_C3 | 0.259944966 | 0.814214506 |
| ALA51_CB_C3 | 0.283258888 | 0.552604471 |
| ALA55_CB_C3 | 0.261821849 | 0.811893696 |
| ALA62_CB_C3 | 0.225927488 | 0.775692196 |
| ARG160_CB_C3 | 0.059989343 | 0.166560291 |
| ARG33_CB_C3 | 0.013345776 | 0.223823989 |

**Table S5.** Feature-diverse tICA features and correlations – GLC transport (cont)

| System | GLC | |
|---|---|---|
| # of features | 498 | |
| **Feature** | **\| tIC1 correlation (*r*) \|** | **\| tIC2 correlation (*r*) \|** |
| **ASN17_CB_C3** | 0.420206348 | 0.692644328 |
| **ASN196_CB_C3** | 0.176421315 | 0.871471685 |
| **ASN5_CB_C3** | 0.064908774 | 0.392921164 |
| **ASN76_CB_C3** | 0.239915948 | 0.835376073 |
| **ASP189_CB_C3** | 0.251233918 | 0.72788253 |
| **CYS138_CB_C3** | 0.339637816 | 0.770163118 |
| **GLN44_CB_C3** | 0.165358819 | 0.269003929 |
| **GLU131_CB_C3** | 0.316026334 | 0.597074128 |
| **GLU163_CB_C3** | 0.003209754 | 0.127343819 |
| **GLU41_CB_C3** | 0.111697694 | 0.140918115 |
| **GLU83_CB_C3** | 0.296952021 | 0.694602036 |
| **GLY13_CA_C3** | 0.317366301 | 0.616424727 |
| **GLY135_CA_C3** | 0.329392007 | 0.724733132 |
| **GLY16_CA_C3** | 0.337834176 | 0.790876124 |
| **GLY184_CA_C3** | 0.222051454 | 0.816637702 |
| **GLY199_CA_C3** | 0.247884205 | 0.739270169 |
| **GLY203_CA_C3** | 0.277536233 | 0.583079664 |
| **GLY42_CA_C3** | 0.170878231 | 0.116092 |
| **GLY79_CA_C3** | 0.251894251 | 0.815150141 |
| **ILE152_CB_C3** | 0.189821634 | 0.275127709 |
| **ILE187_CB_C3** | 0.259338119 | 0.60426558 |
| **ILE34_CB_C3** | 0.106856993 | 0.343879713 |
| **ILE75_CB_C3** | 0.239731732 | 0.796877369 |
| **LEU169_CB_C3** | 0.245241734 | 0.347839044 |
| **LEU173_CB_C3** | 0.328591221 | 0.546483659 |
| **LEU209_CB_C3** | 0.214522523 | 0.446607593 |
| **LEU3_CB_C3** | 0.149863022 | 0.414094403 |
| **LEU71_CB_C3** | 0.221851773 | 0.654857981 |
| **LEU72_CB_C3** | 0.195419372 | 0.730798349 |
| **LYS132_CB_C3** | 0.288170908 | 0.617224714 |
| **LYS37_CB_C3** | 0.009616757 | 0.143383017 |

**Table S5.** Feature-diverse tICA features and correlations – GLC transport (cont)

| System | GLC | |
|---|---|---|
| # of features | 498 | |
| Feature | \| tIC1 correlation (*r*) \| | \| tIC2 correlation (*r*) \| |
| LYS38_CB_C3 | 0.030362614 | 0.157346263 |
| LYS65_CB_C3 | 0.322601443 | 0.584010799 |
| MET1_CB_C3 | 0.429334506 | 0.313075967 |
| MET165_CB_C3 | 0.178980291 | 0.220620513 |
| PHE146_CB_C3 | 0.273936651 | 0.689967093 |
| PHE164_CB_C3 | 0.015924443 | 0.149394973 |
| PHE213_CB_C3 | 0.124672086 | 0.264439175 |
| PHE24_CB_C3 | 0.30197089 | 0.754191929 |
| PHE43_CB_C3 | 0.257254772 | 0.189580938 |
| PHE90_CB_C3 | 0.227195445 | 0.484596167 |
| PRO149_CB_C3 | 0.253543999 | 0.356326084 |
| PRO166_CB_C3 | 0.037143284 | 0.216261603 |
| PRO195_CB_C3 | 0.233991738 | 0.831875062 |
| PRO27_CB_C3 | 0.131072806 | 0.38061222 |
| PRO47_CB_C3 | 0.23926399 | 0.298420507 |
| SER142_CB_C3 | 0.295857143 | 0.770792454 |
| SER151_CB_C3 | 0.15181199 | 0.312150834 |
| SER161_CB_C3 | 0.060020893 | 0.156893699 |
| SER176_CB_C3 | 0.295004754 | 0.791288471 |
| SER20_CB_C3 | 0.312343229 | 0.735891193 |
| SER39_CB_C3 | 0.023879774 | 0.13961193 |
| SER50_CB_C3 | 0.345164555 | 0.487509793 |
| SER54_CB_C3 | 0.344270726 | 0.720414771 |
| THR159_CB_C3 | 0.034110162 | 0.1275464 |
| THR30_CB_C3 | 0.079327373 | 0.282089592 |
| THR4_CB_C3 | 0.091069894 | 0.298265952 |
| THR40_CB_C3 | 0.10807696 | 0.266061955 |
| THR68_CB_C3 | 0.211458538 | 0.404881522 |
| TRP180_CB_C3 | 0.119642409 | 0.848708369 |
| TRP58_CB_C3 | 0.184212226 | 0.837524655 |
| TYR183_CB_C3 | 0.252721602 | 0.805610537 |

**Table S5.** Feature-diverse tICA features and correlations – GLC transport (cont)

| System | GLC | |
|---|---|---|
| # of features | 498 | |
| **Feature** | **\| tIC1 correlation (*r*) \|** | **\| tIC2 correlation (*r*) \|** |
| TYR210_CB_C3 | 0.230704648 | 0.463072636 |
| TYR48_CB_C3 | 0.265335572 | 0.559414263 |
| TYR61_CB_C3 | 0.297259114 | 0.803282071 |
| TYR86_CB_C3 | 0.232277586 | 0.547088273 |
| VAL139_CB_C3 | 0.337457871 | 0.788920751 |
| VAL145_CB_C3 | 0.26366882 | 0.63095545 |
| VAL155_CB_C3 | 0.103438559 | 0.218867494 |
| VAL162_CB_C3 | 0.154413968 | 0.264109669 |
| VAL192_CB_C3 | 0.23225549 | 0.78338912 |
| VAL23_CB_C3 | 0.325337064 | 0.74795983 |
| ALA177_CB_C4 | 0.277099708 | 0.742567582 |
| ALA2_CB_C4 | 0.285309305 | 0.317490319 |
| ALA200_CB_C4 | 0.290532543 | 0.793937773 |
| ALA51_CB_C4 | 0.273549678 | 0.546321928 |
| ALA55_CB_C4 | 0.227135986 | 0.835878644 |
| ALA62_CB_C4 | 0.209418056 | 0.778404623 |
| ARG160_CB_C4 | 0.064405828 | 0.156893342 |
| ARG33_CB_C4 | 0.021906137 | 0.23470122 |
| ASN17_CB_C4 | 0.353901942 | 0.741446509 |
| ASN196_CB_C4 | 0.237253646 | 0.848431574 |
| ASN5_CB_C4 | 0.050965717 | 0.412408809 |
| ASN76_CB_C4 | 0.184806284 | 0.862144085 |
| ASP189_CB_C4 | 0.287048228 | 0.704258383 |
| CYS138_CB_C4 | 0.397284117 | 0.723238319 |
| GLN44_CB_C4 | 0.171100517 | 0.265502145 |
| GLU131_CB_C4 | 0.338658618 | 0.56905677 |
| GLU163_CB_C4 | 0.001984326 | 0.126557863 |
| GLU41_CB_C4 | 0.112736693 | 0.147643042 |
| GLU83_CB_C4 | 0.276679899 | 0.701908994 |
| GLY13_CA_C4 | 0.272697911 | 0.624127563 |
| GLY135_CA_C4 | 0.374279326 | 0.683178975 |

**Table S5.** Feature-diverse tICA features and correlations – GLC transport (cont)

| System | GLC | |
|---|---|---|
| # of features | 498 | |
| Feature | \| tIC1 correlation (*r*) \| | \| tIC2 correlation (*r*) \| |
| GLY16_CA_C4 | 0.266392023 | 0.826765634 |
| GLY184_CA_C4 | 0.262760415 | 0.79519185 |
| GLY199_CA_C4 | 0.286805424 | 0.692528032 |
| GLY203_CA_C4 | 0.2948583 | 0.595962627 |
| GLY42_CA_C4 | 0.178080385 | 0.123433095 |
| GLY79_CA_C4 | 0.190241112 | 0.854985389 |
| ILE152_CB_C4 | 0.193673182 | 0.262487676 |
| ILE187_CB_C4 | 0.264520607 | 0.598003431 |
| ILE34_CB_C4 | 0.097490922 | 0.358485568 |
| ILE75_CB_C4 | 0.136720736 | 0.847527047 |
| LEU169_CB_C4 | 0.237633321 | 0.347905327 |
| LEU173_CB_C4 | 0.32745717 | 0.537464126 |
| LEU209_CB_C4 | 0.221865708 | 0.430817263 |
| LEU3_CB_C4 | 0.130158494 | 0.432982852 |
| LEU71_CB_C4 | 0.167002867 | 0.692290137 |
| LEU72_CB_C4 | 0.161805535 | 0.742434606 |
| LYS132_CB_C4 | 0.319847193 | 0.573511544 |
| LYS37_CB_C4 | 0.004139238 | 0.15382813 |
| LYS38_CB_C4 | 0.0243218 | 0.169805112 |
| LYS65_CB_C4 | 0.328439832 | 0.578123192 |
| MET1_CB_C4 | 0.415226397 | 0.32507658 |
| MET165_CB_C4 | 0.17977408 | 0.225743082 |
| PHE146_CB_C4 | 0.3095296 | 0.644877256 |
| PHE164_CB_C4 | 0.016376207 | 0.146285893 |
| PHE213_CB_C4 | 0.127651399 | 0.258477924 |
| PHE24_CB_C4 | 0.251762676 | 0.784497441 |
| PHE43_CB_C4 | 0.262518133 | 0.195736195 |
| PHE90_CB_C4 | 0.219670223 | 0.490857153 |
| PRO149_CB_C4 | 0.259991989 | 0.334827302 |
| PRO166_CB_C4 | 0.029988852 | 0.214194237 |
| PRO195_CB_C4 | 0.322004317 | 0.786362685 |

**Table S5.** Feature-diverse tICA features and correlations – GLC transport (cont)

| System | GLC | |
|---|---|---|
| # of features | 498 | |
| Feature | l tIC1 correlation (*r*) l | l tIC2 correlation (*r*) l |
| PRO27_CB_C4 | 0.121086152 | 0.390432986 |
| PRO47_CB_C4 | 0.244078182 | 0.290083841 |
| SER142_CB_C4 | 0.37221021 | 0.725630591 |
| SER151_CB_C4 | 0.159351919 | 0.292274496 |
| SER161_CB_C4 | 0.061306847 | 0.151244476 |
| SER176_CB_C4 | 0.264032636 | 0.798903459 |
| SER20_CB_C4 | 0.223577504 | 0.810077002 |
| SER39_CB_C4 | 0.021204596 | 0.151443123 |
| SER50_CB_C4 | 0.352049512 | 0.466427849 |
| SER54_CB_C4 | 0.353390309 | 0.714053375 |
| THR159_CB_C4 | 0.037159814 | 0.121201267 |
| THR30_CB_C4 | 0.072926468 | 0.296827297 |
| THR4_CB_C4 | 0.080787463 | 0.317483165 |
| THR40_CB_C4 | 0.107224691 | 0.275100975 |
| THR68_CB_C4 | 0.209320309 | 0.416644282 |
| TRP180_CB_C4 | 0.148904795 | 0.84659216 |
| TRP58_CB_C4 | 0.17272501 | 0.837979184 |
| TYR183_CB_C4 | 0.245241734 | 0.804819815 |
| TYR210_CB_C4 | 0.232048981 | 0.459213617 |
| TYR48_CB_C4 | 0.254491246 | 0.556956915 |
| TYR61_CB_C4 | 0.325862832 | 0.780135419 |
| TYR86_CB_C4 | 0.216209526 | 0.552835888 |
| VAL139_CB_C4 | 0.401125624 | 0.738623374 |
| VAL145_CB_C4 | 0.291423531 | 0.590282927 |
| VAL155_CB_C4 | 0.108731567 | 0.209571373 |
| VAL162_CB_C4 | 0.153787315 | 0.262626215 |
| VAL192_CB_C4 | 0.283081042 | 0.753142772 |
| VAL23_CB_C4 | 0.275426647 | 0.779747038 |
| ALA177_CB_O3 | 0.245861541 | 0.715123056 |
| ALA2_CB_O3 | 0.281770341 | 0.300869659 |
| ALA200_CB_O3 | 0.235502734 | 0.78842481 |

**Table S5.** Feature-diverse tICA features and correlations – GLC transport (cont)

| System | GLC | |
|---|---|---|
| # of features | 498 | |
| Feature | l tIC1 correlation (*r*) l | l tIC2 correlation (*r*) l |
| ALA51_CB_O3 | 0.303127453 | 0.509852637 |
| ALA55_CB_O3 | 0.295334877 | 0.717947484 |
| ALA62_CB_O3 | 0.207007217 | 0.760059138 |
| ARG160_CB_O3 | 0.06569217 | 0.151448585 |
| ARG33_CB_O3 | 7.22628E-05 | 0.200598814 |
| ASN17_CB_O3 | 0.418604415 | 0.663541863 |
| ASN196_CB_O3 | 0.116154644 | 0.848380532 |
| ASN5_CB_O3 | 0.036527638 | 0.40164609 |
| ASN76_CB_O3 | 0.276077078 | 0.761753366 |
| ASP189_CB_O3 | 0.164169046 | 0.743874334 |
| CYS138_CB_O3 | 0.312349376 | 0.764684047 |
| GLN44_CB_O3 | 0.171492551 | 0.237718346 |
| GLU131_CB_O3 | 0.287284738 | 0.619371233 |
| GLU163_CB_O3 | 0.010490603 | 0.107920051 |
| GLU41_CB_O3 | 0.11760113 | 0.122150217 |
| GLU83_CB_O3 | 0.323595475 | 0.641362389 |
| GLY13_CA_O3 | 0.281874189 | 0.639624756 |
| GLY135_CA_O3 | 0.289082146 | 0.734755798 |
| GLY16_CA_O3 | 0.308823606 | 0.795588902 |
| GLY184_CA_O3 | 0.116000068 | 0.827473495 |
| GLY199_CA_O3 | 0.239724214 | 0.671574743 |
| GLY203_CA_O3 | 0.263665336 | 0.509129199 |
| GLY42_CA_O3 | 0.167535086 | 0.099717898 |
| GLY79_CA_O3 | 0.288051964 | 0.725349142 |
| ILE152_CB_O3 | 0.194682589 | 0.250235191 |
| ILE187_CB_O3 | 0.198193213 | 0.626253897 |
| ILE34_CB_O3 | 0.122640016 | 0.315506096 |
| ILE75_CB_O3 | 0.261568653 | 0.733582977 |
| LEU169_CB_O3 | 0.249769845 | 0.334368065 |
| LEU173_CB_O3 | 0.338463543 | 0.539889954 |
| LEU209_CB_O3 | 0.218696554 | 0.424452726 |

**Table S5.** Feature-diverse tICA features and correlations – GLC transport (cont)

| System | GLC | |
|---|---|---|
| # of features | 498 | |
| **Feature** | **I tIC1 correlation (*r*) I** | **I tIC2 correlation (*r*) I** |
| LEU3_CB_O3 | 0.128376291 | 0.419962141 |
| LEU71_CB_O3 | 0.20632386 | 0.637805014 |
| LEU72_CB_O3 | 0.166528515 | 0.71406905 |
| LYS132_CB_O3 | 0.260571184 | 0.638964611 |
| LYS37_CB_O3 | 0.021359057 | 0.120132953 |
| LYS38_CB_O3 | 0.040558875 | 0.126984219 |
| LYS65_CB_O3 | 0.285576181 | 0.597860062 |
| MET1_CB_O3 | 0.414189339 | 0.312700898 |
| MET165_CB_O3 | 0.179019213 | 0.194810695 |
| PHE146_CB_O3 | 0.283544651 | 0.656973746 |
| PHE164_CB_O3 | 0.005220991 | 0.130691379 |
| PHE213_CB_O3 | 0.128685446 | 0.243158528 |
| PHE24_CB_O3 | 0.328999745 | 0.686893792 |
| PHE43_CB_O3 | 0.255061994 | 0.176578425 |
| PHE90_CB_O3 | 0.245362855 | 0.450389801 |
| PRO149_CB_O3 | 0.256628309 | 0.335518217 |
| PRO166_CB_O3 | 0.046829519 | 0.19787024 |
| PRO195_CB_O3 | 0.208517563 | 0.784697819 |
| PRO27_CB_O3 | 0.142603135 | 0.348852528 |
| PRO47_CB_O3 | 0.237164893 | 0.288271048 |
| SER142_CB_O3 | 0.288777183 | 0.693310607 |
| SER151_CB_O3 | 0.163253797 | 0.287830284 |
| SER161_CB_O3 | 0.066925231 | 0.138602996 |
| SER176_CB_O3 | 0.310250977 | 0.734330173 |
| SER20_CB_O3 | 0.333982812 | 0.658224509 |
| SER39_CB_O3 | 0.035469554 | 0.116825033 |
| SER50_CB_O3 | 0.347610432 | 0.439311398 |
| SER54_CB_O3 | 0.36238353 | 0.617167426 |
| THR159_CB_O3 | 0.041748963 | 0.110418932 |
| THR30_CB_O3 | 0.093663299 | 0.265272626 |
| THR4_CB_O3 | 0.069582492 | 0.304740673 |

**Table S5.** Feature-diverse tICA features and correlations – GLC transport (cont)

| System | GLC | |
|---|---|---|
| # of features | 498 | |
| **Feature** | **\| tIC1 correlation (*r*) \|** | **\| tIC2 correlation (*r*) \|** |
| **THR40_CB_O3** | 0.119962065 | 0.237932431 |
| **THR68_CB_O3** | 0.18207313 | 0.421066016 |
| **TRP180_CB_O3** | 0.004572625 | 0.832777705 |
| **TRP58_CB_O3** | 0.203984833 | 0.788826378 |
| **TYR183_CB_O3** | 0.155658753 | 0.824372736 |
| **TYR210_CB_O3** | 0.232987253 | 0.436965313 |
| **TYR48_CB_O3** | 0.285195554 | 0.522988356 |
| **TYR61_CB_O3** | 0.269385681 | 0.796044414 |
| **TYR86_CB_O3** | 0.251820533 | 0.509068847 |
| **VAL139_CB_O3** | 0.304973513 | 0.780269294 |
| **VAL145_CB_O3** | 0.287264059 | 0.57724965 |
| **VAL155_CB_O3** | 0.112229484 | 0.198315338 |
| **VAL162_CB_O3** | 0.158148142 | 0.241016793 |
| **VAL192_CB_O3** | 0.119144375 | 0.794576662 |
| **VAL23_CB_O3** | 0.347362698 | 0.673324828 |
| **ALA177_CB_O4** | 0.30931222 | 0.626195243 |
| **ALA2_CB_O4** | 0.260974893 | 0.335597597 |
| **ALA200_CB_O4** | 0.327835482 | 0.716410885 |
| **ALA51_CB_O4** | 0.276196133 | 0.516443173 |
| **ALA55_CB_O4** | 0.194737161 | 0.807315931 |
| **ALA62_CB_O4** | 0.112268247 | 0.791734772 |
| **ARG160_CB_O4** | 0.077007532 | 0.131048473 |
| **ARG33_CB_O4** | 0.0132582 | 0.216278671 |
| **ASN17_CB_O4** | 0.326941973 | 0.727440126 |
| **ASN196_CB_O4** | 0.289249456 | 0.754421052 |
| **ASN5_CB_O4** | 0.029743332 | 0.424170184 |
| **ASN76_CB_O4** | 0.10503932 | 0.848668236 |
| **ASP189_CB_O4** | 0.26073281 | 0.68442749 |
| **CYS138_CB_O4** | 0.35307843 | 0.721494791 |
| **GLN44_CB_O4** | 0.17302987 | 0.236092899 |
| **GLU131_CB_O4** | 0.300016509 | 0.581991968 |

**Table S5.** Feature-diverse tICA features and correlations – GLC transport (cont)

| System | GLC | |
|---|---|---|
| # of features | 498 | |
| Feature | l tIC1 correlation (*r*) l | l tIC2 correlation (*r*) l |
| GLU163_CB_O4 | 0.012759408 | 0.105438017 |
| GLU41_CB_O4 | 0.118420228 | 0.134089461 |
| GLU83_CB_O4 | 0.270753548 | 0.678880874 |
| GLY13_CA_O4 | 0.236445544 | 0.607227288 |
| GLY135_CA_O4 | 0.340597447 | 0.685866083 |
| GLY16_CA_O4 | 0.261322114 | 0.807754125 |
| GLY184_CA_O4 | 0.264314798 | 0.75235425 |
| GLY199_CA_O4 | 0.306795287 | 0.575085003 |
| GLY203_CA_O4 | 0.310699125 | 0.519288967 |
| GLY42_CA_O4 | 0.176937971 | 0.108798681 |
| GLY79_CA_O4 | 0.170705723 | 0.81857805 |
| ILE152_CB_O4 | 0.202690556 | 0.246620953 |
| ILE187_CB_O4 | 0.233766821 | 0.60169557 |
| ILE34_CB_O4 | 0.103004963 | 0.340231088 |
| ILE75_CB_O4 | 0.093681585 | 0.816799985 |
| LEU169_CB_O4 | 0.246172687 | 0.318606468 |
| LEU173_CB_O4 | 0.347946668 | 0.499865795 |
| LEU209_CB_O4 | 0.241508816 | 0.396334865 |
| LEU3_CB_O4 | 0.102064635 | 0.451454377 |
| LEU71_CB_O4 | 0.122355412 | 0.692208099 |
| LEU72_CB_O4 | 0.064658002 | 0.735253833 |
| LYS132_CB_O4 | 0.296530099 | 0.569585634 |
| LYS37_CB_O4 | 0.010910656 | 0.14099074 |
| LYS38_CB_O4 | 0.027819703 | 0.158314332 |
| LYS65_CB_O4 | 0.276422418 | 0.603226096 |
| MET1_CB_O4 | 0.390084822 | 0.33512185 |
| MET165_CB_O4 | 0.186567374 | 0.200734862 |
| PHE146_CB_O4 | 0.3340354 | 0.594506025 |
| PHE164_CB_O4 | 0.004546797 | 0.125179869 |
| PHE213_CB_O4 | 0.142258914 | 0.229462957 |
| PHE24_CB_O4 | 0.255353433 | 0.739349373 |

**Table S5.** Feature-diverse tICA features and correlations – GLC transport (cont)

| System | GLC | |
|---|---|---|
| # of features | 498 | |
| Feature | \| tIC1 correlation (*r*) \| | \| tIC2 correlation (*r*) \| |
| PHE43_CB_O4 | 0.258974277 | 0.177995674 |
| PHE90_CB_O4 | 0.219451201 | 0.471669451 |
| PRO149_CB_O4 | 0.270099682 | 0.317952452 |
| PRO166_CB_O4 | 0.045100985 | 0.194949622 |
| PRO195_CB_O4 | 0.338685596 | 0.719961347 |
| PRO27_CB_O4 | 0.121108482 | 0.353883054 |
| PRO47_CB_O4 | 0.241228156 | 0.260547596 |
| SER142_CB_O4 | 0.373437361 | 0.659852918 |
| SER151_CB_O4 | 0.167294437 | 0.265235819 |
| SER161_CB_O4 | 0.07269414 | 0.128586544 |
| SER176_CB_O4 | 0.304259401 | 0.69236126 |
| SER20_CB_O4 | 0.240906791 | 0.748158279 |
| SER39_CB_O4 | 0.027589874 | 0.14013347 |
| SER50_CB_O4 | 0.339714186 | 0.421820353 |
| SER54_CB_O4 | 0.332816235 | 0.665022163 |
| THR159_CB_O4 | 0.048030312 | 0.10059472 |
| THR30_CB_O4 | 0.079864875 | 0.277989339 |
| THR4_CB_O4 | 0.061808381 | 0.340528168 |
| THR40_CB_O4 | 0.112397466 | 0.257188028 |
| THR68_CB_O4 | 0.179228801 | 0.451712891 |
| TRP180_CB_O4 | 0.203722612 | 0.764363009 |
| TRP58_CB_O4 | 0.035750158 | 0.833493476 |
| TYR183_CB_O4 | 0.241298232 | 0.772976636 |
| TYR210_CB_O4 | 0.252784459 | 0.423124753 |
| TYR48_CB_O4 | 0.255569513 | 0.535679716 |
| TYR61_CB_O4 | 0.244602838 | 0.802612484 |
| TYR86_CB_O4 | 0.21674997 | 0.52923809 |
| VAL139_CB_O4 | 0.385390887 | 0.701977212 |
| VAL145_CB_O4 | 0.299302246 | 0.522471186 |
| VAL155_CB_O4 | 0.118573227 | 0.188742998 |
| VAL162_CB_O4 | 0.166623075 | 0.23658557 |

**Table S5.** Feature-diverse tICA features and correlations – GLC transport (cont)

| System | GLC | |
|---|---|---|
| # of features | 498 | |
| Feature | \| tIC1 correlation (*r*) \| | \| tIC2 correlation (*r*) \| |
| VAL192_CB_O4 | 0.264337295 | 0.710818379 |
| VAL23_CB_O4 | 0.299084098 | 0.70361835 |
